# A free energy landscape screen reveals the disordered conformational ensemble of tropoelastin

**DOI:** 10.64898/2026.02.03.703507

**Authors:** Sean E. Reichheld, Lisa D. Muiznieks, Zi Hao Liu, Brandon J. Payliss, Fred W. Keeley, Simon Sharpe

**Affiliations:** Molecular Medicine Program, Hospital for Sick Children, Toronto, Ontario, Canada; Department of Biochemistry, University of Toronto, Toronto, Ontario, Canada

**Keywords:** Intrinsic disorder, conformational ensemble, protein structure, NMR, tropoelastin

## Abstract

Understanding how proteins explore their conformational energy landscapes is essential for linking sequence to function, yet current ensemble methods are limited by sampling inefficiency and poor scalability to large disordered systems. Here we introduce a free energy landscape screen (FELS), a conceptually different approach that replaces sampling-centric ensemble fitting with broad exploration of energy landscapes, screening thousands of landscape shapes—from highly funneled to flat and rugged. By systematically biasing and evaluating large conformer pools according to contact propensities derived from experiment, FELS efficiently identifies sets of conformers that best reproduce experimental data and highlights candidates for structural refinement, without being restricted by chain length or amount of disorder. To demonstrate the power of this approach we applied it to a previously intractable system, human tropoelastin (hTE), a ∼700-residue precursor of elastin. FELS provides the first experimentally defined atomistic view of the hTE conformational ensemble, revealing that this protein is intrinsically disordered yet exhibits distinct local secondary structure and specific, transient medium- and long-range contacts that organize its ensemble. These findings reconcile long-standing conflicting models and demonstrate that FELS provides a general, experimentally driven framework for mapping conformational energy landscapes of large proteins across the continuum between structural order and disorder.

## Introduction

Understanding protein structure is central to uncovering biological mechanisms and designing therapeutics ^1^. Most progress has focused on globular proteins, which fold into a single, stable three-dimensional structure. As a result, the Protein Data Bank (PDB) now contains over 200,000 entries ^2,3^, the vast majority of which represent compact, folded domains. In contrast, intrinsically disordered proteins (IDPs) and regions (IDRs), which lack stable tertiary structure and instead populate broad, interconverting ensembles^4^, remain underrepresented, with fewer than 700 ensembles reported^5^. Yet more than 30% of eukaryotic proteins are predicted to contain long disordered regions involved in many cellular processes, where flexibility enables function ^6–9^.

Ensemble modeling has become a central strategy for interpreting experimental data on disordered proteins ^10–13^. These methods typically generate a conformer pool from statistical coil models or PDB-based loop libraries ^14–18^, then apply sampling strategies such as Monte Carlo optimization or Bayesian reweighting to fit ensembles to experimental restraints ^10–13^. Despite their utility, these approaches are fundamentally limited by the quality of the initial pool, the challenge of efficiently sampling sufficient conformational space, and the number and quality of experimental restraints^12,19–23^. If the pool lacks relevant structures, or sampling is biased or limited, the resulting ensembles may fit the data in aggregate yet fail to capture the true structural preferences of individual protein conformations. Mean distance restraints often lack the sensitivity to capture short-range molecular forces that stabilize conformations, leading to under-restrained ensembles that are either over-averaged or artificially heterogeneous^19,20,24^. These limitations are compounded as chain length increases because the number of potential conformations grows exponentially ^25,26^ and atomistic-level restraints are harder to acquire^27^, helping explain why nearly all published ensembles are for IDPs shorter than ∼200 residues^5^.

A useful way to describe protein structural space is as a free energy landscape, in which folded proteins occupy funneled landscapes with a deep global minimum corresponding to the native structure ^26^, while disordered proteins populate broad, shallow landscapes with many similar-energy minima, reflecting high conformational entropy and structural diversity ^28^. In reality, most proteins lie along a conformational continuum, adopting a mixture of defined and flexible regions whose energy landscapes shift with sequence context, solvent conditions, post-translational modifications, or binding partners ^28,29^. Crucially, because free energy landscapes describe conformational preferences in probabilistic terms^30,31^, they are agnostic to the degree of structural order in the system being studied.

To model structure across this conformational spectrum, we developed a free energy landscape screen (FELS) method that is guided by experimental data. Contact propensities are first derived from paramagnetic relaxation enhancement (PRE) nuclear magnetic resonance (NMR) probes and used to guide tunable weighting schemes applied to a large pool of pre-generated conformers. These schemes bias conformer selection while generating landscapes ranging from highly funneled to effectively flat. By reweighting and evaluating an entire conformer pool comprising hundreds of thousands of structures, we estimate the effective energy of each conformation and assess how well the pool reflects the experimental data. This evaluation enables ensemble construction without the need to exhaustively sample conformational space, and identifies opportunities for pool refinement. By avoiding random selection, iterative reweighting, and long molecular dynamics (MD) simulations, FELS is computationally efficient and avoids overfitting, even for large proteins.

In this study, we apply FELS, along with a modular NMR chemical shift assignment strategy, to a previously intractable system, human tropoelastin (hTE). hTE is a large ∼700 amino acid protein that serves as the soluble precursor to elastin and forms the core of elastic fibers in vertebrate tissues ^32–34^. While some studies have proposed a relatively stable global fold based on small-angle x-ray scattering (SAXS) and MD simulations ^35–37^, others have suggested that hTE is an IDP with extensive conformational heterogeneity, which is required for its assembly and function ^38,39,32^. Our analysis provides the first atomistic view of this protein and reveals that hTE exists as a dynamic conformational ensemble with limited global order but distinct local structure and specific yet transient medium- and long-range contacts. These features suggest a flexible, modular architecture with varying levels of local compaction, which may facilitate elastin assembly and function. More broadly, FELS provides a general strategy for mapping diverse conformational landscapes, with potential for future integration with machine learning to enhance inference and expand capabilities.

## Results and Discussion

### Modular and disordered architecture of tropoelastin

NMR spectroscopy is ideally suited for examining the structure and dynamics of IDPs with atomistic precision^23,27^. However, the combination of disorder and low-complexity, repetitive sequences causes severe spectral overlap, making large IDPs like hTE particularly challenging^27^. Tropoelastin is distinctive among IDPs because its sequence is highly hydrophobic and arranged into alternating hydrophobic and crosslinking domains (Figure 1A), a modular organization thought to underlie its self-assembly and elasticity^32–34,39^. Hydrophobic domains (HDs), enriched in proline and glycine and other hydrophobic residues, promote both elasticity, and the phase separation required for self-assembly (often referred to as coacervation), while lysine-containing crosslinking domains (CLDs) facilitate covalent crosslinking into polymeric materials. Prior studies on elastin-like proteins (ELPs) and isolated domains of hTE have suggested strong domain independence, with local sequence motifs dictating structural and functional properties irrespective of global context^40,41,32^. Indeed, synthetic ELPs composed of simple repeats mimicking this modular arrangement are able to recapitulate hTE’s phase separation, self-assembly and elastic properties^40,42,43^.

**Figure 1.**
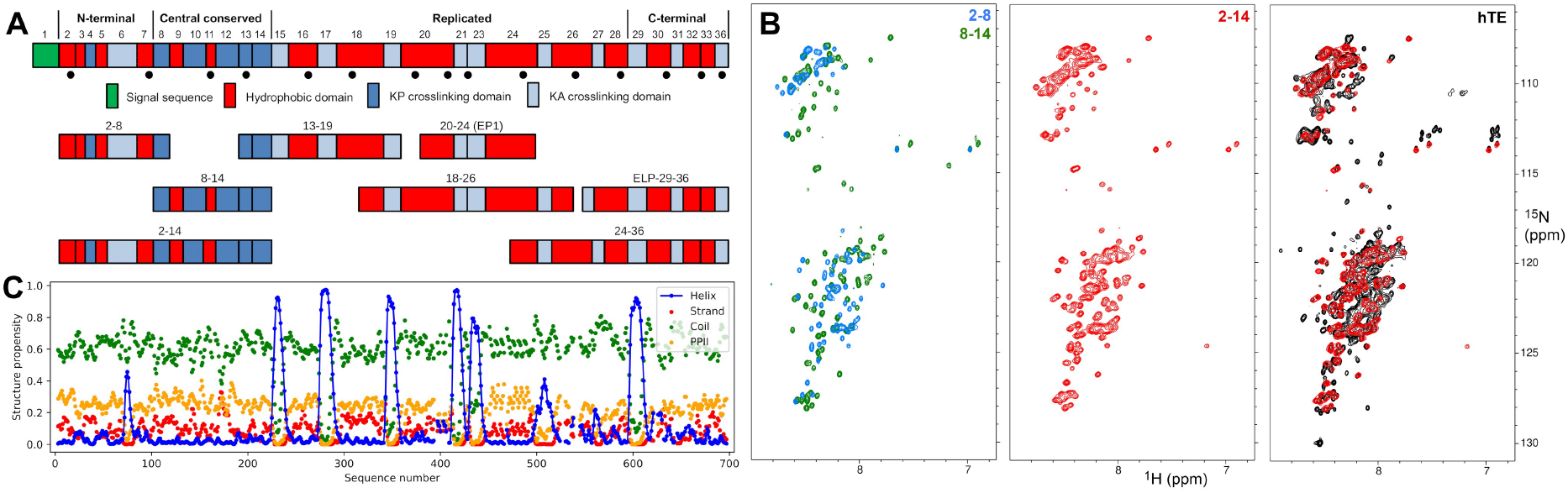
Sequence-specific secondary structure assignment of hTE. (**A**) Schematic representation of full-length hTE and fragment constructs used for NMR chemical shift assignment. Distinct regions are separated by vertical lines and domains are numbered according to their exons, consistent with historical nomenclature for hTE. Dots below the full-length hTE schematic indicate cysteine mutation sites used for MTSL PRE labeling. (**B**) ^1^H-^15^N HSQC spectra of 2-8, 8-14, 2-14 fragment constructs, and full-length hTE. (**C**) Sequence-specific secondary structure propensities derived from chemical shifts using δ2D^44^.

To facilitate NMR resonance assignments in full-length hTE, we used a strategy that takes advantage of this modular behaviour. Chemical shifts from smaller, overlapping fragments of hTE (Figure 1A; SI Table 1) were used as reference points to aid assignment of corresponding regions in full-length hTE. ^1^H-^15^N HSQCs of hTE fragments and full-length hTE were typical of IDPs, with poor ^1^H spectral dispersion and narrow linewidths, irrespective of construct size. The presence of a single set of amide resonances suggests that the hTE structural ensemble undergoes rapid ns to μs timescale exchange between conformers (Figure 1B; SI Figure 1). Overlays of spectra from smaller fragments, such as constructs 2-8 and 8-14, closely matched those from sequentially overlapping fragments (e.g. 2-14; Figure 1B; SI Figure 2), supporting our fragment-based approach and the lack of persistent contacts between the fragments, which would be expected to impact chemical shifts. The sequential backbone ^15^N and ^13^C chemical shifts from all constructs, including full-length hTE were in close agreement with each other (SI Figure 3), suggesting that structural propensities are largely defined by local sequence properties rather than by long-range contacts or cooperative folding.

Using fragment and full-length data, we compiled a master set of chemical shift assignments for hTE (Assignments in SI Table 2; Assignment statistics in SI Table 3). δ2D^44^ analysis identified predominant α-helical propensity in the KA crosslinking domains, with limited secondary structure in hydrophobic and KP crosslinking domains (Figure 1C). Helical propensity varied between KA domains (SI Figure 4), with longer polyalanine stretches and those immediately preceded by a proline exhibiting increased α-helix stability.

### Building tropoelastin structural models with experimentally biased secondary structure

NMR chemical shift data indicate that hTE is highly disordered, requiring structurally diverse ensembles for accurate modeling. To this end, a pool of ∼200,000 3D models of hTE was generated using IDPConformerGenerator^14,45^. A C688S/C693S mutant sequence was used to match the constructs used in PRE experiments and conformation sampling was biased using δ2D secondary structure propensities.

The initial conformer pool was highly diverse but also markedly expanded, with a mean radius of gyration (*R*_g_) of 81.7 Å (SI Figure 5A). This is roughly 15–20 Å larger than expected for an IDP of comparable length and 20–25 Å larger than the envelope size estimated from SAXS (SI Figure 6), which was consistent with previous measurements^35^. The scattering profiles and *P*(*r*) distributions of WT hTE and the C688S/C693S mutant were very similar, supporting the use of this mutant in PRE experiments (SI Figure 6). Backbone torsion angles sampled the expected Ramachandran space for each amino acid type (SI Figure 7), and back-calculated secondary structure propensities agreed well with δ2D inputs (SI Figure 5B). HDs displayed features consistent with previous findings^40,46,32^, including a high proportion of bends, hydrogen-bonded turns and random coil, while CLDs contained varying amounts of α-helix.

### A free energy landscape screen (FELS) for building protein ensembles

Despite starting from a conformer pool that recapitulated experimental secondary structure propensities, exhaustive sampling of the 3D conformational space of large IDPs such as hTE (698 residues) is computationally infeasible because the number of possible conformations (estimated at > 10^330^ different backbone conformations for hTE) grows exponentially with chain length^25,26^. To overcome this barrier, we use contact propensities from PRE NMR data to bias the conformational search (Figure 2, SI Figure 8). PRE data is commonly used to obtain interatomic distances for structure determination of proteins and protein complexes by measuring the increased relaxation and concurrent signal attenuation of NMR active nuclei in close proximity to a paramagnetic center^47,48^.

**Figure 2.**
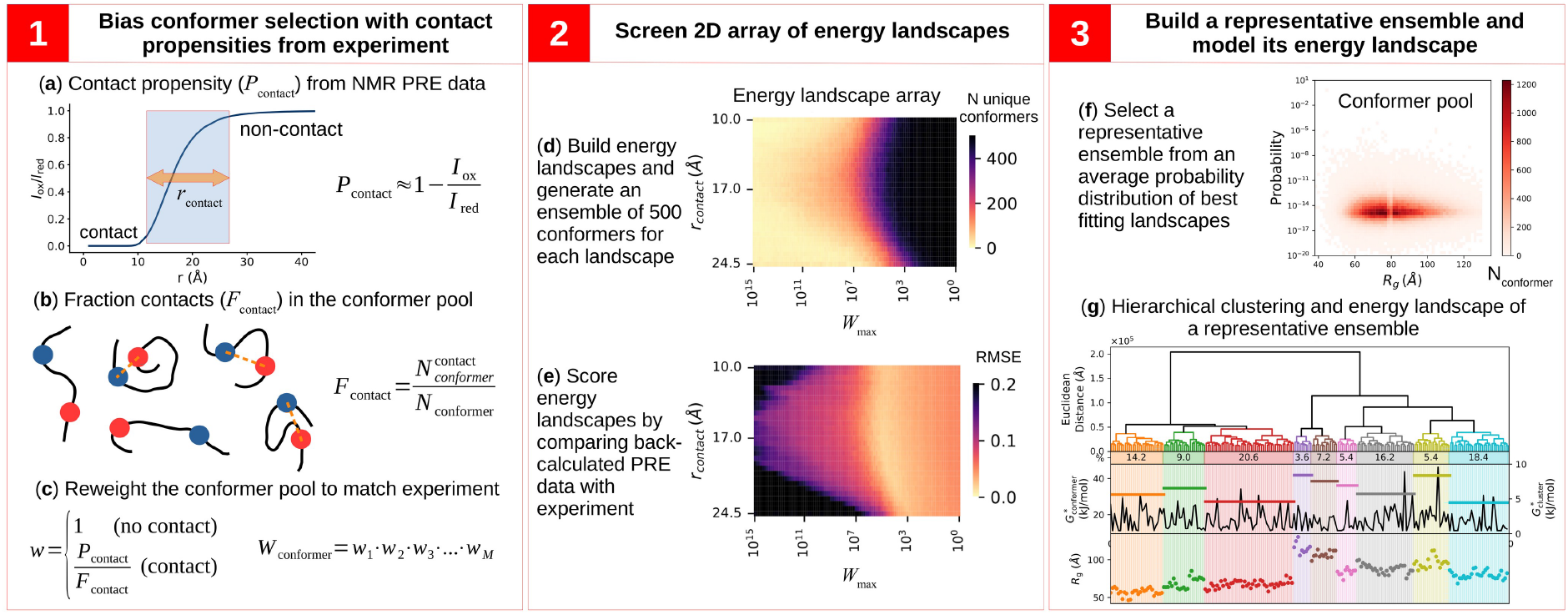
Free energy landscape screen (FELS). **Step 1: Bias conformer selection with contact propensities from experiment**. In (**a**), experimental peak-intensity ratios (*I*_ox_/*I*_red_) are converted to contact propensities (*P*_contact_). In (**b**), a pool of conformer models is generated and the fraction of contacts (*F*_contact_) is calculated 200across a range of contact distances (*r*_contact_) that span the linear region of the *I*_ox_/*I*_red_ vs. distance (*r*) relationship shown in (**a**). In the simplified structural models shown in (**b**), the paramagnetic center is represented as a blue circle and the amide proton as a red circle, with contacts indicated by orange dashed lines. In this example, three of five conformers satisfy the contact criterion, yielding an *F*_contact_ value of 3/5. In (**c**), pairwise reweighting factors (*w*) are derived to resolve discrepancies between experimental contact propensities and contact proportions in the conformer pool. Weights for non-contact and contact sites are set to 1 and *P*_contact_/*F*_contact_, respectively. For a given conformer, the overall conformer weight (*W*_conformer_) is computed as the product of the individual contact weights across all *M* restraints. **Step 2: Screen 2D array of energy landscapes**. In (**d**), a single ensemble of 500 conformers is generated for each landscape in a two-dimensional matrix using weighted random sampling. The horizontal dimension (coarse adjustment) corresponds to the maximum weight (*W*_max_) allowed for any one conformer, while the vertical dimension (fine adjustment) corresponds to variations in *r*_contact_. Ensemble diversity is assessed by counting the number of unique conformers in each ensemble and is visualized as a heat map across the landscape array. High *W*_max_ values produce funneled landscapes with low conformer diversity, whereas low *W*_max_ values produce flatter landscapes with higher diversity. In (**e**), ensemble-averaged PRE curves are generated from back-calculated data and compared with experiment to compute RMSE values, which are likewise displayed as a heat map over the same two-dimensional landscape matrix. **Step 3: Build a representative ensemble and model its energy landscape**. In (**f**), weights from the best-fitting energy landscapes (within 2.5% RMSE of the global minimum) are averaged to generate a master probability distribution, visualized as a heat map of conformer probability vs. radius of gyration (*R*_g_). From this distribution, representative ensembles of the energy landscape are generated. In (**g**), hierarchical clustering of a representative ensemble is performed using a C_a_ distance matrices. A pseudo-free-energy is calculated for individual conformers (*G*^*^_conformer_) and for clusters of structurally related conformers (*G*^*^_cluster_). Refer to SI Figure 8 for a detailed flow chart of the FELS methodology.

To collect PRE data across the entire hTE sequence, we generated 15 single-cysteine mutant constructs for site-specific labeling with a methanethiosulfonate spin label (MTSL) (Figure 1A). One construct retained the natural cysteine at position C688 (with a C693S mutation), while the others were in a double mutant (C688S/C693S) background. ^1^H-^15^N HSQC spectra were recorded for all MTSL-hTE constructs in the presence or absence of ascorbic acid. Proximity to the unpaired electron of oxidized MTSL increases spin relaxation, decreasing protein NMR signal intensity (*I*_ox_) relative to the reduced state (*I*_red_). For an IDP, the underlying distribution of structures is assumed to be an ensemble of rapidly interconverting conformations such that *I*_ox_/*I*_red_ provides a measure of the population-weighted and time-averaged distance between the paramagnetic center and the nuclear spins of interest^47,24^.

As expected for an IDP, strong signal attenuation was observed at sites sequentially proximal to the spin label, with weaker attenuation extending ∼150–200 residues from the labeling site (SI Figure 9A). However, the decay in signal attenuation as a function of sequence distance from the PRE site was not uniform across datasets, suggesting the presence of transient, yet specific, medium- and long-range contacts within a disordered ensemble (SI Figure 9A). In all constructs, the upper baseline of *I*_ox_/*I*_red_ values fell slightly below 1.0, indicative of intermolecular PRE effects, which were corrected for by subtracting an average baseline derived from distant residues across all datasets (SI Figure 9B; SI Figure 10).

Most approaches for PRE-based ensemble modeling treat *I*_ox_/*I*_red_ values as population-weighted distance restraints^10–13^. However, PRE data suffer from limited resolution and a nonlinear relationship between signal attenuation and distance (see Figure 2, step 1). At short distances (≤10–12 Å), maximal signal attenuation is observed, making shorter distances indistinguishable. Beyond this range, signal attenuation falls off rapidly due to the 1/*r*^6^ distance dependence, limiting the ability to distinguish between long distances (>20–25 Å). As a consequence of ensemble averaging, intermediate *I*_ox_/*I*_red_ values may reflect an ambiguous combination of short, intermediate and long distances. These limitations suggest that transforming PRE data into precise population-weighted distance restraints overstates its resolving power. Moreover, using PRE data solely as a post hoc evaluation tool is inherently inefficient due to the large number of conformational ensembles that may need to be generated and tested before arriving at a representative fit. These challenges motivated us to instead treat *I*_ox_/*I*_red_ values as estimates of contact propensities, with interpretation roughly analogous to established semi-quantitative treatment of nuclear Overhauser effect (NOE) data. As detailed in SI Methods 1.7, use of PRE data to estimate contact propensities provides an efficient framework for generating dynamic IDP ensembles.

Our FELS implements this idea by systematically testing a broad set of possible energy landscapes, ranging from funneled (folded-like) to flat and rugged (disordered-like). This is achieved by using *I*_ox_/*I*_red_ PRE data to guide the conformational search and evaluate the entire conformer pool, rather than using it merely as a retrospective fitting tool. As shown in Figure 2, step 1, our workflow begins by converting PRE-derived intensity ratios into contact propensities (*P*_contact_: Eq. S1) and comparing them to the fraction of contacts (*F*_contact_: Eq. S2) observed in the conformer pool. Discrepancies between all pairwise contact ratios (*w*_contact_: Eq. S3) are used to derive reweighting factors for each conformer (*W*_conformer_: Eq. S4), which increase the likelihood of selecting conformers that reflect the experimental data.

*W*_conformer_ values are computed across a two-dimensional matrix of conditions by systematically varying the contact distance threshold (*r*_contact_) and the maximum allowable weight (*W*_max_) assigned to any individual conformer. *W*_max_ sets an upper bound on *W*_conformer_, limiting the influence of highly favored structures and thereby controlling the steepness and ruggedness of the resulting energy landscape. Together, these parameters define the shape of each tested landscape (Figure 2, step 2). For each unique combination of *r*_contact_ and *W*_max_, an ensemble of 500 conformers is generated using random weighted sampling. This process is highly efficient, enabling the screening of tens of thousands of diverse candidate landscapes within minutes, with each yielding a distinct ensemble. By design, this approach can recover both highly specific, funneled energy landscapes containing only a few strongly upweighted conformers and flatter, more structurally diverse landscapes consistent with disordered ensembles.

For each ensemble, the average *I*_ox_/*I*_red_ values are back-calculated and compared with experimental data to give a global root-mean-square error (RMSE) value across all data points. The best-fit energy landscapes (those with RMSE values within 2.5% of the global minimum) are then combined into a master probability distribution (Figure 2, step 3), visualized as a heat map of conformer probability (*P*_conformer_) vs. *R*_g_. From this distribution, multiple ensembles are generated and the one with the median RMSE is selected, resulting in a representative ensemble that reflects both a best-fit to structural features and the appropriate degree of conformational diversity, avoiding overfitting to a narrow subset of the conformer pool.

To visualize and interpret this representative energy landscape, structurally similar conformers are grouped using hierarchical clustering based on pairwise differences in backbone C_α_ positions^20^. Each conformer’s free energy (*G*^*^_conformer_) is estimated from its probability using the standard Boltzmann relation (Eqs. S7, S8). Aggregate pseudo-free energies are then calculated for each structurally distinct cluster (*G*^*^_cluster_) by summing the probabilities of its member conformers and normalizing over the total probability of the representative ensemble (Eqs. S9–11). This approach not only identifies dominant structural clusters but moves toward a quantitative model of the conformational free energy surface.

### Initial screen of the tropoelastin’s energy landscape

Ensembles of hTE were generated from the pool of ∼200,000 IDPConformerGenerator structures using the FELS approach. PRE restraint sites were selected to capture key features in the *I*_ox_/*I*_red_ curves indicative of medium- and long-range contacts, while minimizing interdependence between neighboring residues in our reweighting factor calculations. Despite restraint sites comprising only ∼5% of all data points, RMSE at restraint sites correlated strongly with global RMSE (Figure 3A), validating the use of minimal restraints to guide the search while avoiding over-weighting of individual contacts.

**Figure 3.**
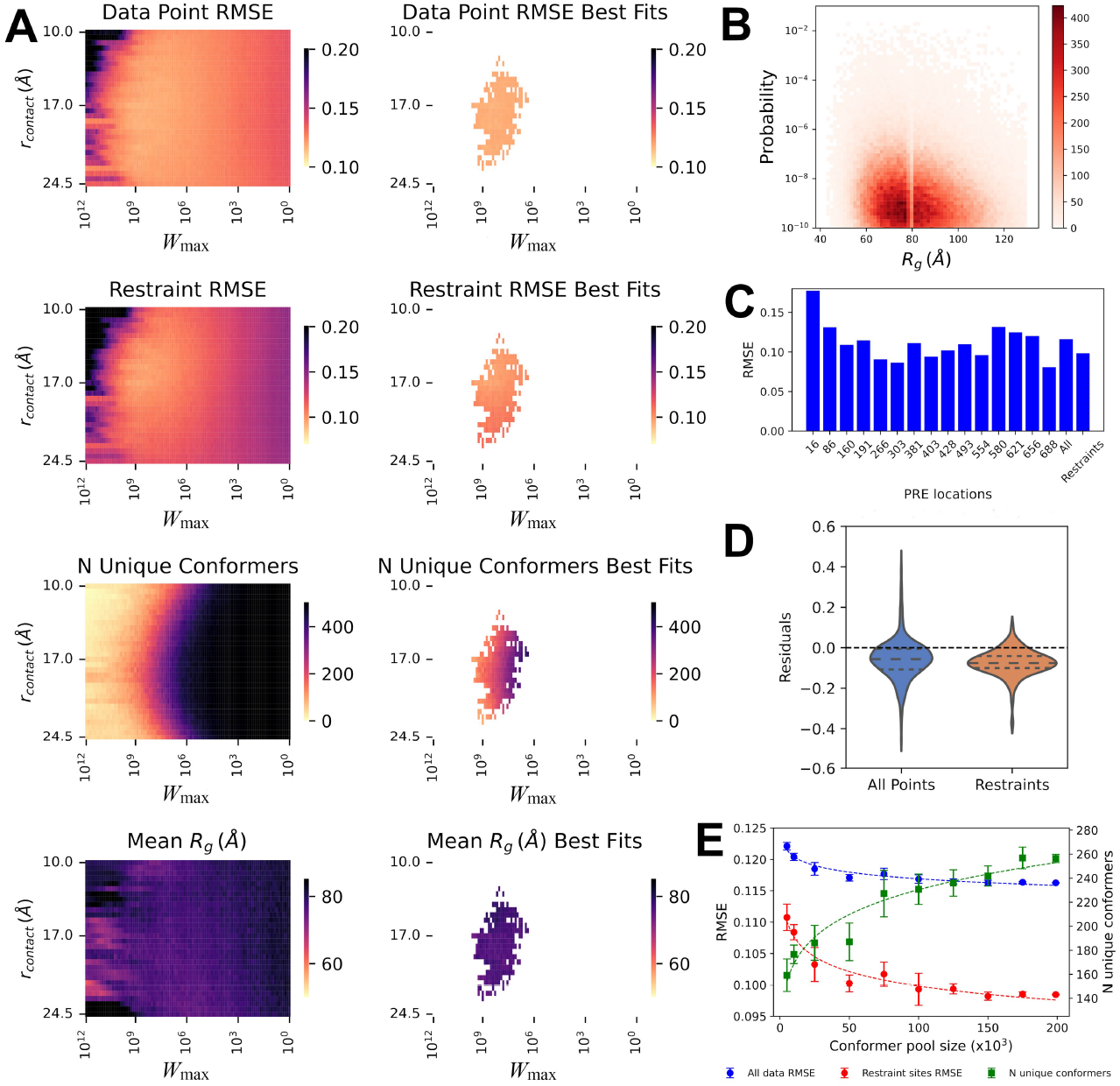
Initial application of FELS. Experimentally derived PRE data served as input for FELS runs against the hTE conformer pool generated by IDPConformerGenerator. Averaged back-calculated *I*_ox_/*I*_red_ curves were generated using a fixed-point PRE model^47^ and compared to experiment to compute RMSE values for each ensemble. (**A**) Heat maps of all-data-point RMSE, restraint-site RMSE, ensemble diversity and *R*_g_ are shown for the full screen (left) and best-fitting ensembles (right). (**B**) Heat map of conformer probability vs. *R*_g_, averaged across the top-scoring energy landscapes (within 2.5% RMSE of the global minimum). The resulting distribution was then used to generate the final representative ensemble. (**C**) Bar chart of RMSE values for each PRE site, for all PRE data points (All), and for all restraint sites (Restraints). (**D**) Violin plots of residuals. (**E**) Plots showing the change in RMSE values and ensemble diversity (N unique conformers) with increasing conformer-pool size.

The initial application of FELS revealed limitations of the hTE conformer pool. RMSE heat maps showed that no energy landscape yielded strong fits, and even the best-fitting ensembles exhibited only moderate agreement (Figure 3A). The best landscapes remained highly diverse (>200 unique conformers) but also contained strongly upweighted conformers (*W*_max_ 10^6^–10^9^) that skewed toward more compact structures (Figure 3B). A representative ensemble derived from the FELS yielded a global RMSE of 0.115 (Figure 3C), above the theoretical minimum of ∼0.08, which was calculated by fitting a moving average of the data. This high baseline error reflects both the experimental variability and sequence-specific relaxation effects inherent to PRE datasets (See SI Figure 11 for a breakdown of datasets and fits). Residual distributions (Figure 3D) show a negative skew, with predictions generally exceeding experimental values, suggesting a lack of medium- and long-range contacts in the conformer pool. The mean *R*_g_ of 77.5 Å, was ∼20 Å larger than our estimate from SAXS experiments (SI Figure 6), confirming that compact states were underrepresented.

Adjusting the conformer pool size showed that RMSE improvements plateaued beyond ∼50,000 conformers (Figure 3E), indicating that brute-force expansion cannot replace better structural sampling. Notably, ensemble diversity increased in step with better fits, emphasizing that hTE must be modeled as a structurally heterogeneous ensemble.

### Validating FELS with synthetic data

To test whether the limitations in our initial hTE fitting reflected shortcomings in the conformer pool or weaknesses in the FELS methodology itself, we performed a proof-of-principle screen using synthetic PRE data from ensembles of known composition. This allowed direct assessment of FELS in recovering defined input populations and quantifying the relationship between ensemble diversity and fitting accuracy.

Random input ensembles of varying sizes were selected from 41 conformers identified in the original ∼200,000-member IDPConformerGenerator pool, each with at least five contacts at restraint sites. Contacts were defined as having back-calculated *I*_ox_/*I*_red_ values of < 0.5, corresponding to distances of < 16 Å in a fixed-point PRE model. For each input ensemble, synthetic *I*_ox_/*I*_red_ curves were generated for all 15 MTSL-labeled sites and used as input for FELS against the original conformer pool.

Despite the size and complexity of the pool, FELS accurately recovered the input structures in the best-fit ensembles. Even with inputs of 15 or more conformers, ∼40% of the representative ensembles consisted of input conformers, and ∼90% of the input set was recovered at least once (Figure 4A). Ensemble diversity scaled with the number of unique input conformers (Figure 4B), and although global RMSE increased slightly with diversity, it remained low overall (∼0.04 for ensembles with >= 5 conformers; Figure 4C). A strong correlation (*R*^2^ = 0.93) between RMSE values calculated across all data points and those from restraint sites alone (Figure 4D) further validated the use of minimal restraints to drive accurate ensemble selection.

**Figure 4.**
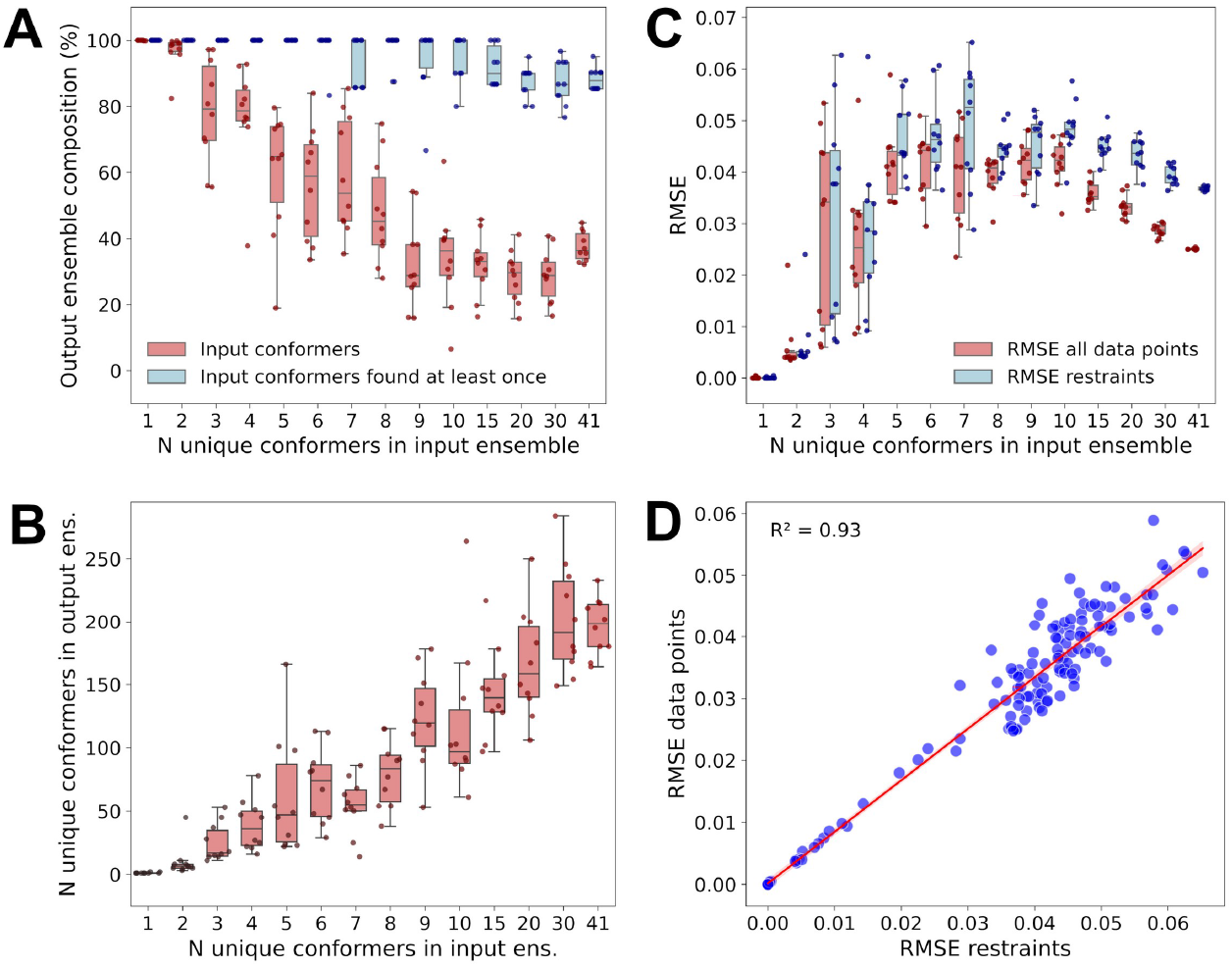
FELS proof-of-principle. Averaged back-calculated *I*_ox_/*I*_red_ curves, computed using a fixed-point PRE model^47^, from each subset served as synthetic input data for FELS runs against the IDPConformerGenerator pool. Ten unique input ensembles were tested at each level of diversity—quantified by the number of unique conformers. (**A**) Percentage of conformers in the output ensemble that were present in the input ensemble (red) and the percentage of input conformers recovered at least once in the output ensemble (blue). (**B**) Output-ensemble diversity as a function of input-ensemble diversity. (**C**) RMSE of global fits to synthetic input data as a function of input-ensemble diversity. (**D**) Correlation between global RMSE calculated from all data points and RMSE from restraint sites alone (*R*^2^ = 0.93).

### Refining ensemble models with MD

While the proof-of-principle results demonstrated that FELS can faithfully reconstruct ensembles when representative conformers are present, the initial hTE conformer pool lacked enough compact structures to satisfy the experimental PRE contact propensities. To address this, ∼1000 conformers consistently upweighted in the initial hTE screen (*R*_g_ < 85 Å) were refined by short all-atom MD simulations in explicit solvent. The goal was to enrich medium- and long-range contacts by promoting local sidechain packing and hydrophobic collapse—features difficult to capture using stepwise conformer generation alone. Each 4-ns trajectory generated 10 snapshots, producing ∼10,000 new conformers. A second round of refinement on 120 upweighted structures from this pool yielded an additional 1,200 conformers.

MD refinement produced only modest decreases in *R*_g_ but significantly reduced sidechain solvent-accessible surface area (SASA), consistent with improved hydrophobic packing (SI Figure 12). Secondary structure propensities were not significantly altered by MD refinement, and remained consistent with the NMR data. The combined pool of refined models spanned a broad size range (∼40– 85 Å) with an average *R*_g_ of 56.2 Å (Figure 5A), close to our size estimate from SAXS (SI Figure 6) and similar to previous estimates^35^. MD also drove a marked increase in medium-to long-range contacts (Figure 5B), accompanied by a modest increase in backbone bends (Figure 5C; SI Figure 12E, F).

**Figure 5.**
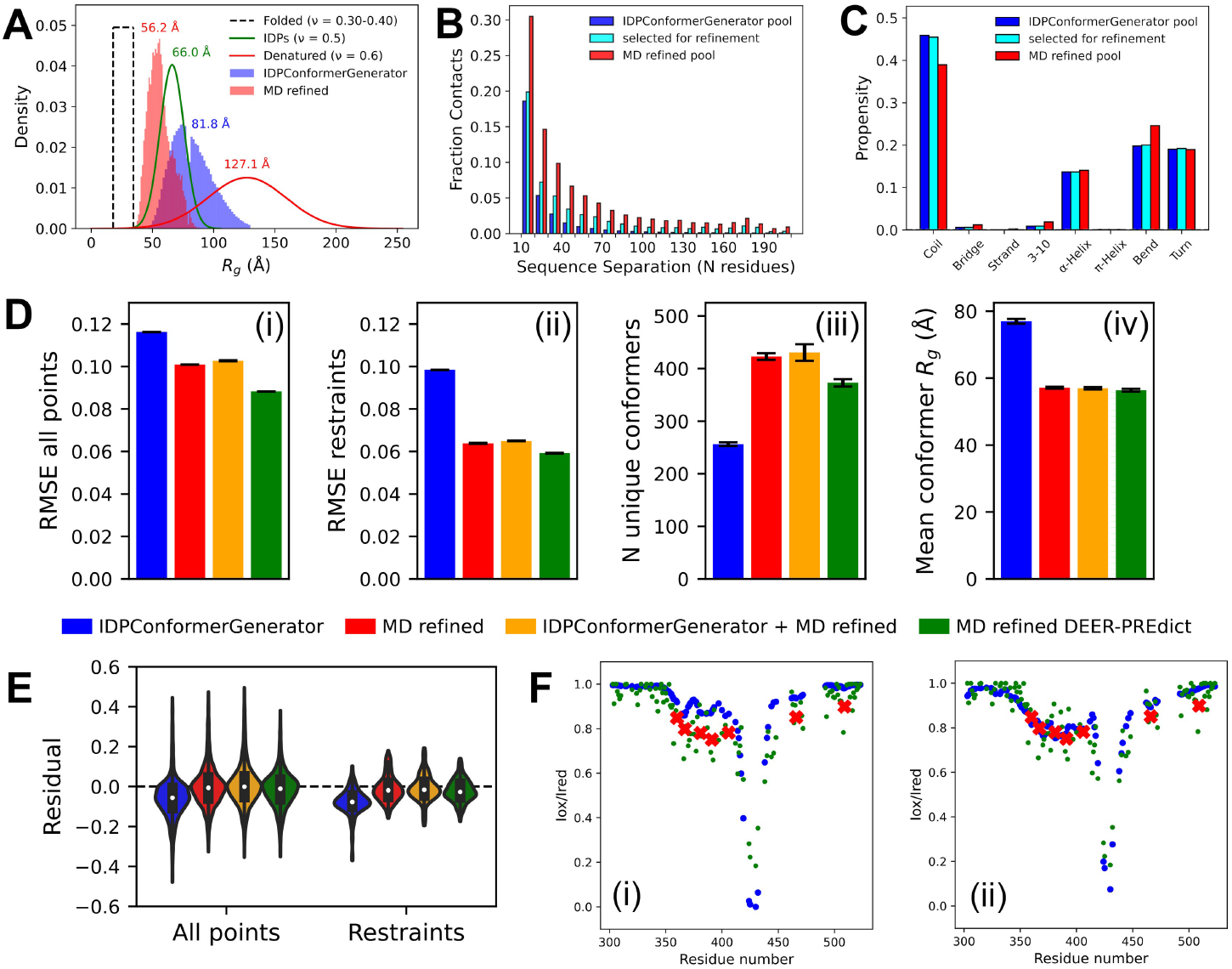
Tropoelastin ensemble refinement through MD simulations. (**A**) Comparison of conformer size distributions for the hTE MD-refined pool (round 1 + round 2) with the IDPConformerGenerator pool and those expected for folded, intrinsically disordered, and denatured proteins of the same chain length (698 residues). Mean conformer sizes are shown above each distribution. Size distributions for model protein types were calculated as in SI Figure 5. (**B**) Fraction of contacts as a function of sequence distance from all PRE sites and (**C**) secondary-structure content calculated using DSSP^61^ for each pool and for conformers selected from the IDPConformerGenerator pool for refinement during the initial screen. (**D**) Representative ensemble summaries from FELS runs against the two separate pools and the combined pool using a fixed-point PRE model^47^, compared to runs against the MD-refined pool using the flexible PRE model from DEER-PREdict^49^. Shown are (i) RMSE for all data points, (ii) RMSE at restraint sites, (iii) ensemble diversity, and (iv) mean *R*_g_. Error bars represent standard deviations from five independent runs. Additional results for FELS runs using different combinations of conformer pools and PRE back-calculation models are shown in SI Figure 13. (**E**) Violin plots of *I*_ox_/*I*_red_ residuals for representative ensembles. (**F**) Examples of individual PRE-site fits (larger blue circles) to experimental data (smaller green circles) and restraints (red Xs). Single-site PRE fits labeled at residue 428 in the hTE hinge region using (i) the IDPConformerGenerator pool with a fixed-point PRE model and (ii) MD-refined pool with the flexible PRE model from DEER-PREdict. Single site fits for all sites for these two screens are provided in SI Figures 11 and 15.

FELS runs using the refined pools showed markedly improved fits to experimental PRE data. Global and restraint-site RMSE values decreased relative to the original pool, with the best results obtained using the combined refined pool with the flexible PRE model from DEER-PREdict^49^ (Figure 5D; SI Figures 13–14). Residuals between experimental and back-calculated *I*_ox_/*I*_red_ values were more symmetrically distributed around zero (Figure 5E), indicating reduced systematic fitting bias. Importantly, the best-fitting energy landscapes required much less conformer upweighting (*W*_max_ 10^3^– 10^5^) than the initial screen (*W*_max_ 10^6^–10^9^), showing that the refined pool more closely matched the experimental data. When screening combined pools of IDPConformerGenerator and MD-refined conformers, fits remained strong, confirming the screen’s ability to identify relevant conformers even within large, diverse pools containing tens of thousands of higher energy structures.

Notably, improved fits were not solely driven by MD-induced compaction or a loss of conformer diversity, as ensemble diversity increased when using MD-refined conformer pools (Figure 5Diii; SI Figure 13C), and final fits showed consistent mean *R*_g_ values near 57 Å (Figure 5Div; SI Figure 13D) irrespective of the pool composition. Some MD-refined conformers were even downweighted, preventing overrepresentation of specific contacts, reflecting FELS’s ability to adjust for overrepresented features in the conformer pool. Nearly all single-site PRE fits showed better agreement when using the global FELS fitting method applied to pools containing MD-refined conformers (Figure 5F; SI Figure 13E; SI Figure 15), indicating that MD refinement broadened the accessible conformational landscape and enhanced fit quality across the full length of hTE. A second proof-of-principle test using the refined pool further confirmed FELS’s ability to recover input ensembles with varying amounts of conformer diversity (SI Figure 16).

As independent validation, back-calculated SAXS curves generated from representative ensembles were compared with experimental scattering curves (SI Figure 16). All back-calculated SAXS profiles were in excellent agreement with experiment. Importantly, our experimental SAXS data are closely aligned with previously reported scattering profiles^35,36,50^, whose authors interpreted hTE as adopting a relatively stable globular fold—highlighting the known limitations of using scattering data alone to unambiguously resolve the degree of structural disorder in proteins^51,52^.

Together, these results demonstrate that FELS successfully identified a large set of suitable candidates for MD refinement, leading to enhanced conformer realism by improving sidechain packing and intramolecular contacts. This, in turn, enabled FELS to recover ensembles that more closely matched the PRE data while preserving structural heterogeneity.

### Modeling the free-energy landscape of hTE

Having established representative conformational ensembles that recapitulate experimental PRE data, we next sought to model the conformational free-energy surface of hTE. Our ensemble fitting yields weights for each conformer, allowing estimation of the free-energy landscape by converting these weights into probabilities and then into energies and organizing the conformational space through hierarchical clustering of backbone positions. This approach quantifies the ensemble’s diversity and structure while identifying the basins of the underlying energy landscape.

Since structural alignment of diverse IDP conformers is not possible, we clustered the SAXS-validated ensembles using Euclidean distances derived from intramolecular C_α_ distance matrices^20^ with one example shown in Figure 6A. A pseudo-free energy was calculated for each conformer (Eq. S8) and each structural cluster (Eq. S11). The resulting dendrogram revealed a diverse distribution of conformers across the ensemble, spanning collapsed and expanded states, with multiple low-energy basins consistent with hTE existing as a structurally heterogeneous ensemble. As expected given hTE’s high hydrophobicity, the ensemble was slightly biased toward collapsed states. Backbone ^15^N *R*_2_ relaxation rates (SI Figure 18B) were generally in the 1-20 s^-1^ range, consistent with rapid conformational exchange and highlighting the disconnect between local chain dynamics and overall molecular tumbling. Higher *R*_2_ values were observed for regions anticipated to have slowed conformational exchange, such as CLDs with high helical propensity (e.g. residues 227–238, 276–289, 345–355, 414–426, 432–444, 499–513 and 598–612) as well as sites of longer-lived local collapsed states (outlined below).

**Figure 6.**
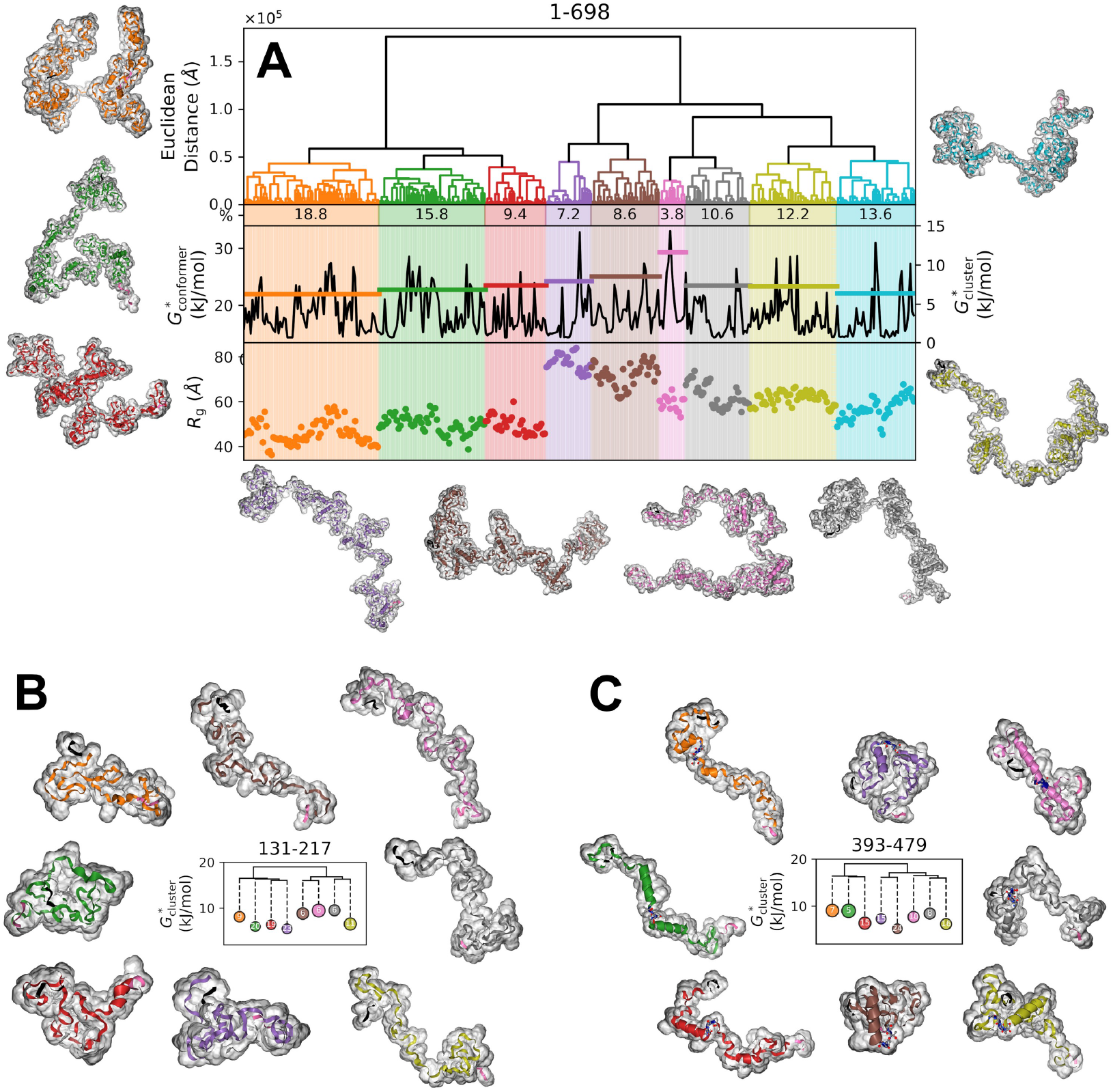
Hierarchical clustering and energy landscape of tropoelastin. (**A**) Dendrogram of hierarchical clustering for a representative ensemble from a FELS run against the MD-refined pool using the flexible PRE model from DEER-PREdict^49^. Euclidean distances were calculated from differences between intramolecular C_α_ distance matrices^20^. Shaded regions denote distinct structural clusters, matching the colors of the surrounding representative structures, which are shown with a gray solvent-exposed shell. The percentage contribution of each cluster to the representative ensemble is shown below the dendrogram. Pseudo-free energies of individual conformers (*G*^*^_conformer_) and structural clusters (*G*^*^_cluster_) are shown, with *R*_g_ of individual conformers plotted below the energy profiles. Clustering of discrete sequence regions spanning overlapping one-eighth ranges within the full-length hTE representative ensemble was also performed with complete results shown in SI Figure 19. These are represented as simplified hybrid clustering-energy diagrams, where solid lines indicate structural relatedness proportional to Euclidean distances and dashed lines represent cluster energy levels. In these simplified diagrams, circle size is proportional to the mean *R*_g_ of the cluster, and the number within each circle indicates the percent contribution of that cluster to the representative ensemble. Examples of local clustering are shown in (**B**) and (**C**): (**B**) Residues 131–217, spanning the central conserved region. (**C**) Residues 393–479, spanning the hinge (residues 427– 431, shown as balls and sticks) region. Representative structures of each cluster show only the portion of full-length hTE corresponding to the sequence range used for clustering. For all structures (full-length and sequence-range), N-terminal residues are shown in black and C-terminal residues are shown in hot pink.

Although our chemical shift data suggested structural independence between different sequence regions, PRE *I*_ox_/*I*_red_ profiles revealed asymmetry and discrete contact propensities along the sequence. To examine local structural variation, we clustered overlapping windows of 1/8 sequence length (SI Figure 19). We observed notable differences in conformer compactness and cluster energies across regions, with the N- and C-termini, residues 131–217, and residues 350–436 exhibiting the highest degree of compaction (SI Figure 20A). Importantly, no region was restricted to a narrow set of conformations, with each displaying a range from collapsed to expanded states. When longer sequence windows were used, no notable differences in compaction were observed (SI Figure 20B and C) and there was very little correlation between the *R*_g_ of shorter and longer segments (SI Figure 20D–F), suggesting that most structural differences are driven by transient interactions between residues separated by less than ∼80–100 amino acids.

Our results significantly advance the structural understanding of hTE by providing site-specific insights that go well beyond the limited resolution provided by previous SAXS^35,36,50^ and NMR^53^–55 studies. Residues 131–217 span the central conserved portion of hTE, an evolutionarily well-conserved hydrophobic region^56^,57 with a high proportion of aromatic residues. Our structural analysis reveals that this region adopts collapsed states in more than 70% of conformers without forming a specific globular structure (Figure 6B), reflecting a locally collapsed but dynamically exchanging segment. Another sequence block of hTE that has been of particular interest is the “hinge region” which spans residues 413–444 and has been proposed to form an anti-parallel helix-turn-helix^58^. Counter to this expectation, our clustering of the hinge (including portions of adjacent domains) reveals high conformational diversity and varying helical content (Figure 6C).

Using FELS, we demonstrate that hTE adopts a heterogeneous and highly dynamic ensemble containing local regions of transient compaction. Importantly, our findings reconcile previous SAXS and NMR observations by showing that the apparent compactness of hTE arises from transient, sequence-specific interactions rather than persistent tertiary structures. Furthermore, the presence of locally collapsed states within an overall flexible ensemble provides a structural rationale for the heterogeneity in crosslinking patterns observed in mass spectrometry studies of natural elastin^38^. Finally, while atomistic MD simulations have provided valuable insights into local structural features of tropoelastin^37,59,60^, the extensive conformational diversity indicated by our ensembles suggests that readily accessible MD timescales are insufficient to fully capture the breadth of the conformational landscape. Our results thus provide the first detailed, experimentally grounded structural framework for understanding tropoelastin’s behavior in solution and its role in elastin assembly, offering a benchmark for future studies aiming to capture its conformational ensemble with greater fidelity.

## Conclusions and Future Directions

The methodology developed in this work advances the structural analysis of large IDPs by enabling the generation of experimentally consistent, atomistic ensembles that capture both local structural propensities and the extensive conformational diversity characteristic of these systems. By combining PRE-derived contact propensities with SAXS validation within an efficient FELS framework, we have demonstrated a practical approach for mapping the conformational free-energy surface of a large IDP, tropoelastin, with site-specific resolution that extends well beyond what is achievable by SAXS or NMR chemical shift data alone.

We find that tropoelastin exists as a heterogeneous structural ensemble, but with preferred locally compact states that are transiently populated. This resolves the long-standing tension between models describing tropoelastin as either folded or disordered by showing that hTE occupies multiple shallow basins within an overall disordered ensemble. This landscape helps explain why highly simplified and repetitive ELPs or smaller tropoelastin fragments can recapitulate the elastic function of full-length tropoelastin, yet exhibit variations in mechanical properties such as stiffness, energy loss, and stress relaxation^40,62–67^. It also provides a mechanistic basis for understanding the crosslinking behavior reported by Schräder *et al*.^38^, who observed largely stochastic intermolecular crosslinking—consistent with a dynamic, disordered protein—but a higher abundance of certain intramolecular crosslinks. Together, these results establish a framework for probing how preferred conformational states influence early stages of tropoelastin self-assembly and elastic function, and how interactions with other extracellular matrix components, such as fibrillin, the fibulins, and lysyl oxidase, contribute to organizing the elastic fiber.

This methodology provides a foundation for addressing important challenges in protein structural biology. While atomistic MD simulations continue to provide valuable insights into local conformational preferences, much longer simulation timescales are required to sample the conformational landscape of large systems, particularly for transient, medium- and long-range contacts. Our FELS approach complements MD by efficiently exploring diverse conformational states while remaining grounded in experimental data. Furthermore, this strategy can be readily integrated with other experimental methods used to probe distance distributions and dynamics, such as single-molecule Forster resonance energy transfer (FRET) and double electron-electron resonance (DEER) electron paramagnetic resonance (EPR) measurements and should be adaptable to machine learning environments for greater efficiency and to generate more complete energy landscapes.

Crucially, our approach is not limited to fully disordered proteins. By systematically and rapidly screening a broad range of energy landscapes—from highly funneled, folded-like basins to flatter, rugged, disordered-like surfaces—FELS can identify and characterize ensembles across the continuum of protein structural states, including folded proteins with local disordered regions, molten globule-like states, and fully disordered proteins. The ability to rapidly obtain detailed conformation ensembles for large IDPs or large IDR containing proteins represents a significant advancement, as demonstrated here for the 698 amino acid IDP tropoelastin.

## Acknowledgements

The authors thank Julie Forman-Kay for insightful discussions during development of the FELS methodology. This work was funded by CIHR Project Grant FRN399475 and HSF Grant in Aid G-18-0022249 (to SS). SAXS measurements were carried out in the SickKids Structural and Biophysical Core Facility. This research was enabled in part by support provided by Compute Ontario and the Digital Research Alliance of Canada.

## Author Contributions

SS, SR, FWK and LM conceptualized the project. SS and SR designed experiments, analyzed data and wrote the manuscript. SR, LM and BP carried out experiments. SR developed the equations and code for the FELS methodology. ZHL developed software and provided computational and theoretical support for conformer generation.

## Conflict of Interest Statement

The authors declare no competing interests.

## Data Availability

NMR chemical shift data have been deposited in the Biological Magnetic Resonance Data Bank (BMRB) under accession numbers listed in SI Table 1. Structural ensembles generated in this study have been deposited in the Protein Ensemble Database (PED) and will be released upon publication. The FELS code and all additional data and analysis scripts supporting the findings of this study are available from the authors upon reasonable request.

## Notes

### Competing Interest Statement

The authors have declared no competing interest.

