## Supplemental Information for "A free energy landscape screen reveals the disordered conformational ensemble of tropoelastin"

### 1. Supplementary Methods

#### 1.1 Protein expression and purification

All protein expression constructs were purchased from GenScript and verified by DNA sequencing. In designing tropoelastin fragment constructs, care was taken to avoid placing construct termini within KA-type crosslinking domains (CLDs), where truncation could disrupt extended α-helical secondary structure. Amino acid sequences of full-length hTE and fragments used for chemical shift assignments are shown in SI Table 1. Expression and purification of recombinant proteins followed previously described methods[1]. Briefly, proteins were expressed in *Escherichia coli* BL21(DE3) using a pET32b plasmid (Invitrogen) in which the target sequence was fused to an N-terminal thioredoxin tag. The thioredoxin was removed by cyanogen bromide digestion in 70% formic acid under denaturing conditions. Proteins were purified by ion-exchange FPLC followed by reverse-phase HPLC. For isotopic enrichment, proteins were expressed in *E. coli* grown in M9 minimal medium supplemented with $^{15}N$-$NH_4Cl$ and $^{13}C$-glucose as the sole nitrogen and carbon sources. For chemical shift assignment, tropoelastin was expressed in M9 minimal medium prepared in 70% $D_2O$.

#### 1.2 NMR spectroscopy

All NMR experiments were performed on 500–600 µL samples using a triple-resonance 5-mm TXI MicroProbe equipped with Z-gradient and temperature control on Bruker Avance III spectrometers operating at a $^1H$ frequency of 600 or 700 MHz. Samples were buffered in 50 mM phosphate, pH 7.0, with 10% $D_2O$ and varying concentrations of sodium chloride. $^1H$ chemical shifts were referenced to DSS, prepared separately in 50 mM sodium phosphate, 50 mM NaCl, and 10% $D_2O$ at pH 7.0, with the methyl resonance set to 0.00 ppm. $^{13}C$ and $^{15}N$ chemical shifts were referenced indirectly using the IUPAC-recommended frequency ratios relative to DSS. Data were acquired with Bruker TopSpin using standard 2D and 3D pulse sequences with water suppression. Spectra were processed with NMRPipe[2] and analyzed with CcpNmr Analysis[3]. Figures were generated in CcpNmr Analysis and with the Python package nmrglue[4].

#### 1.3 Chemical shift assignment

Chemical shift assignments of tropoelastin and its fragments were carried out on solution samples at 10 °C. $^1H$, $^{13}C$, and $^{15}N$ resonances were assigned using standard 2D, 3D, and 4D NMR methods. Non-uniform sampling (NUS) was employed at 12% for 3D experiments and 4% for 4D experiments, and spectra were reconstructed using the iterative soft thresholding (ist) algorithm implemented in NMRPipe[2]. The following experiments (pulse sequences provided by Bruker BioSpin) were used for sequential backbone assignment of full-length human tropoelastin (hTE) and its fragments: $^1H$-$^{15}N$ HSQC, HSQC-TOCSY, NOESY-HSQC, HNCA, HN(CO)CA, HNCO, HN(CA)CO, CBCANH, (H)CC(CO)NH, HCAN, HNN, and HN(CO)N. Additional 4D HNCACO, HNCOCA, and HNNH experiments were required for sequential assignments of full-length hTE. Overlaying assigned fragment spectra with the full-length hTE spectra was necessary to resolve assignment ambiguities.

#### 1.4 Secondary structure prediction and conformer generation

Secondary structure propensities for hTE were predicted from $^1HN$, $^{15}N$, $^{13}C_\alpha$, $^{13}CO$, and $^1H_\alpha$ chemical shifts using the δ2D web server for intrinsically disordered proteins[5]. Since chemical shifts from smaller hTE fragments correlated strongly with those of full-length hTE (SI Figure 3), fragment chemical shifts were pooled (SI Table 2) to provide more comprehensive input for secondary structure predictions.

Approximately 200,000 full-length hTE conformers were generated on the Beluga Digital Research Alliance of Canada cluster[6] using the IDPConformerGenerator software package[7,8], with parameters summarized in SI Table 3. δ2D-derived secondary structure propensities were provided as input to bias conformer generation toward locally preferred backbone geometries. Conformers were built from the C688S/C693S sequence background, which was the mutant background used for PRE experiments. Conformers were generated in batches of 1000-2000 structures, each submitted to a full compute node (40 CPUs, maximum RAM). Structures were assembled by fragment selection from loop and helix regions of the Protein Data Bank using segment sizes of 3–8 residues. The --long and --long-ranges flags were enabled to accelerate conformer construction. Approximately half of the conformers were generated by stitching together six fragments of roughly equal size, with an average rate of 13.8 conformers $h^{-1}core^{-1}$. The remaining half were built by stitching together twelve fragments at an average rate of 45.5 conformers $h^{-1}core^{-1}$. Initial conformer pools were filtered to remove structures with steric clashes, chain breaks, or improper geometries. Conformers with a radius of gyration ($R_g$) greater than ~20 nm were also removed due to incompatibility with CRYSOL[9] small-angle X-ray scattering (SAXS) back-calculations. Protons were added to conformers using the pdb2gmx utility in the GROMACS[10] molecular dynamics software suite.

**1.5 SAXS analysis**

WT hTE and hTE C688S/C693 at 100 µM and 200 µM in 50 mM sodium phosphate buffer (pH 7) and 50 mM NaCl were analyzed in a capillary cell at 10 °C using a benchtop Automated SAXSpace instrument (Anton Paar, Montréal, Canada) with a temperature-controlled sample stage. Data represent the average of 60 × 120 s acquisitions, collected in high-intensity mode with line collimation. Background subtraction, desmearing, and primary processing were performed with SAXS Treat and SAXS Analysis software (Anton Paar). 200 µM samples provided stronger high-$q$ signal but exhibited a low-$q$ upturn consistent with attractive, monomer-monomer interactions. This upturn was largely alleviated in 100 µM samples, which showed increase noise at medium–high $q$. Pair-distance distribution functions, $P(r)$, were calculated in GNOM[11] using $q$ ranges of 0.0210–0.1502 $Å^{-1}$ (200 µM) and 0.0173–0.1502 $Å^{-1}$ (100 µM). $D_{max}$ was adjusted iteratively until the $P(r)$ function decayed smoothly to zero without oscillations or truncation artifacts, which was achieved for a $D_{max}$ of 200 Å. The GNOM fits for $D_{max}$ values spanning 150–240 Å are summarized in SI Table 5.

Back-calculated SAXS profiles were generated for each conformer with CRYSOL[9] 3.0 using default solvent and hydration parameters ($\Delta\rho = 0.03$ e/Å³, $\rho = 0.334$ e/Å³, spherical harmonics order lmax = 75, 100 points per curve). Output curves were compiled and averaged across ensembles to obtain ensemble $P(r)$ distributions. Experimental and ensemble-derived $P(r)$ curves were compared directly, and radii of gyration ($R_g$) were extracted from the $P(r)$ distributions for quantitative comparison. Comparisons were restricted to $P(r)$ curves and $R_g$ values because these representations are less sensitive to absolute intensity scaling, more robust to differences in concentration or background subtraction, and provide a direct structural basis for evaluating disordered protein ensembles.

**1.6 Paramagnetic relaxation enhancement (PRE) labeling and data collection**

PRE experiments were performed on site-directed cysteine mutants of hTE generated in the C688S/C693S background. Individual residues were mutated to cysteine for site-specific attachment of the paramagnetic spin label MTSL (Toronto Research Chemicals). Cysteine mutation sites for MTSL labeling were chosen to minimize disruption of local structural propensity, solvation, and intramolecular contacts when introducing the hydrophobic MTSL tag. Purified and lyophilized protein was dissolved to ~100–150 µM in 50 mM sodium phosphate, pH 7.0, containing 2 mM DTT, and incubated for 1 h at room temperature to reduce the cysteine site. DTT was removed using a PD-10 desalting spin column (Cytiva). MTSL, prepared as a fresh 100 mM stock in acetonitrile, was immediately added to the protein effluent from the spin column to a final concentration of 2 mM, with the protein concentration maintained at ~100–150 µM in 50 mM sodium phosphate, pH 7.0. Labeling was carried out at room temperature for ~18–24 h with moderate shaking in the absence of light. Excess free label was removed by multiple rounds of dialysis against water, and labeled samples were lyophilized for future use.

$^{1}$H-$^{15}$N HSQC spectra were acquired at 10 °C on ~100 µM protein samples for each mutant in both oxidized (paramagnetic) and reduced (diamagnetic) states. Complete or near-complete labeling with active MTSL was verified by the loss of NMR signal intensity at or adjacent to the labeling site. Diamagnetic samples were generated by adding ascorbic acid to a final concentration of 2 mM directly to the oxidized sample, followed by 2 h incubation at room temperature. PRE effects were quantified as the intensity ratio ($I_{ox}/I_{red}$) of backbone amide cross-peaks.

**1.7 Free energy landscape screen (FELS)**

Given the sigmoidal nature of the $I_{ox}/I_{red}$-distance relationship and the inherent measurement errors associated with PRE data, we propose interpreting $I_{ox}/I_{red}$ values as loose indicators of proximity rather than precise distance measurements. When applied to a single structure, strong signal attenuation ($I_{ox}/I_{red}$ values close to 0) suggests that the PRE label is close (contact) to the amide, while weak or no signal ($I_{ox}/I_{red}$ values close to 1) indicates that the PRE is far (non-contact) from the amide. This interpretation aligns with long-established NMR practices, where NOE intensities are categorized into broad distance ranges based on loosely defined signal intensity strengths of strong, medium, and weak due to known error and uncertainty in these measurements[12]. When this interpretation of the $I_{ox}/I_{red}$ is applied to an average of a conformational ensemble the intensity ratio naturally becomes an estimate of the likelihood or propensity of two sites being close or far from each other. The estimate of contact propensity $P_{\text{contact}}$ between a paramagnetic center at residue $i$ and an amide proton at residue $j$ is expressed as:

$$P_{\text{contact}}(i,j) = 1 - \frac{I_{\text{ox},i,j}}{I_{\text{red},i,j}} \tag{S1}$$

where $I_{ox,i,j}$ is the intensity of the NMR signal for the HN at residue $j$ in the oxidized (paramagnetic) state due to the paramagnetic label at residue $i$, and $I_{\text{red},i,j}$ is the intensity of the NMR signal for the amide proton at residue $j$ in the reduced (diamagnetic) state (See SI Figure 8 for a detailed workflow diagram of the FELS methodology).

While contact propensities provide an estimate of contacts between residue $i$ and residue $j$ derived from NMR data, we can also directly quantify the fraction of contacts across our pool of conformer models. The fraction of contacts, $F_{\text{contact}}(i,j)$, is computed as the fraction of conformers where the distance between residue $i$ and residue $j$ is less than or equal to the contact threshold $r_{\text{contact}}$. This is expressed as:

$$F_{\text{contact}}(i,j) = \frac{\sum_{k=1}^{N_{\text{conformer}}} I\left(r_{i,j,k} \le r_{\text{contact}}\right)}{N_{\text{conformer}}} \tag{S2}$$

where $N_{\text{conformer}}$ is the total number of conformers in the pool of structures, and $I(r_{i,j,k} \le r_{\text{contact}})$ is an indicator function that equals 1 if the distance between residues $i$ and $j$ in conformer $k$ is less than or equal to the contact threshold $r_{\text{contact}}$, and 0 otherwise.

To reconcile the difference between the observed contact propensity derived from experimental data and the calculated fraction of contacts in the conformer pool, a reweighting factor is applied to conformers that contain a contact while those that contain no such contact maintain a weighting of 1. This reweighting adjusts the likelihood of conformers being picked when generating conformational ensembles, ensuring that the weighting of conformers more accurately reflects the experimental observations. Conformers with contacts that are overrepresented will be down-weighted, while those with contacts that are underrepresented will be upweighted. The pairwise reweighting factor $w(i,j)$ is defined as the ratio of the contact propensity from our NMR data to the fraction of contacts in the conformer pool for residues i and j:

$$w_{\text{contact}}(i,j) = \frac{P_{\text{contact}}(i,j)}{F_{\text{contact}}(i,j)} \tag{S3}$$

To refine the probability distribution of the conformer pool and ensure that it accurately reflects all the experimental PRE data for each restraint, a combined weight is calculated for each conformation by multiplying the individual weights from each restraint site. Importantly, not all HN sites are used as restraints; instead, selected sites are chosen to avoid the effects of interdependence, such as those within close proximity in the sequence or near the PRE site. The aggregate conformer weight $W_k$ for a given conformation $k$ is calculated as the product of all pairwise reweighting factors (See Eq. S3) for that conformer across all PRE-HN restraint combinations:

$$W_{\text{conformer},k} = \prod_{(i,j)\in S} w(i,j) \tag{S4}$$

where the product is taken over the set $S$ of selected PRE-HN restraint sites. After reweighting the entire conformer pool, ensembles can be generated through random weighted sampling. However, before attempting to generate ensembles there are two important considerations that must be addressed.

First, the PRE effect is not a perfect binary relationship, as we have modeled it so far. Intermediate distances between the paramagnetic center and the amide protons will still contribute partial signal attenuation. The correct distance cutoff for a contact is, therefore, an unknown variable that may vary depending on the NMR dataset, the protein being studied, and the specific conformer pool used. To account for uncertainties in the exact distances between residues and to explore the sensitivity of our model to different distance cutoff values, we introduce the concept of varying the threshold distance $r_{contact}$. This approach allows us to test different definitions of what constitutes a contact within an ensemble, capturing a broader range of conformational possibilities that might still satisfy the experimental data. The typical range tested spans the ambiguous intermediate distance regime of approximately 10–25 Å.

Second, our combined weighting formula assumes perfect sampling of all possible configurations in the conformer pool. For a protein of 698 residues like hTE, the number of possible backbone conformations is immense, estimated at > $10^{330}$, using a conservative estimate of 2 accessible torsion states for prolines (119 residues), 4 for glycines (175 residues), and 3 for all other residues (404 residues). However, since it is impossible to sample all these combinations when generating conformer models, some conformers in the pool will inevitably be overweighted due to insufficient sampling of alternative contact combinations and configurations. This imbalance can cause the overrepresentation of a small number of conformers, particularly those that perfectly satisfy specific contact restraints that are underrepresented in the pool due to random chance. Consequently, this could lead to poor agreement between back-calculated and experimental data or a lack of conformer diversity due to overfitting.

To mitigate overfitting, we introduced a simple weight-clipping approach, which caps individual $W_{conformer}$ values. By systematically testing a range of maximum weights ($W_{max}$), we explore a vast array of energy landscapes ranging from highly funneled (high $W_{max}$), which favor specific conformers, to flatter landscapes (low $W_{max}$) where a wider range of conformers contribute to the ensemble. Crucially, and unlike current Bayesian maximum entropy (BME) approaches for ensemble building, which actively penalize a lack of conformer diversity, this methodology allows us to efficiently test and validate ensembles whose underlying structures may differ significantly from purely random selection of the conformer pool, while also avoiding problematic overfitting.

The energy landscape screen was performed by recalculating conformer weights across a two-dimensional matrix of $r_{contact}$ and $W_{max}$ values, with no iterative reweighting. For each parameter combination, conformer weights were updated according to Eqs. S1–S4, and a single ensemble was generated by random weighted sampling using the built-in Python 3 module random (function random.choices) with probabilities proportional to the normalized weights. Each ensemble was then evaluated by calculating the root-mean-square error (RMSE) between the experimental and back-calculated $I_{ox}/I_{red}$ ratios using either a fixed-point PRE model[13] or a flexible PRE model from DEER-PREdict[14], producing one RMSE value per cell in the matrix.

To identify the most representative regions of the landscape, the parameter combinations with the lowest RMSE values (within 2.5% of the global minimum RMSE value) were selected from the global matrix. The conformer probabilities from these best-fitting ensembles were combined into an averaged probability matrix, with each ensemble weighted inversely to its RMSE so that better-fitting ensembles contributed proportionally more. From this averaged probability distribution, 1,001 ensembles were generated by probabilistic sampling, and the ensemble whose RMSE corresponded to the median of this distribution was defined as the representative ensemble.

##### 1.8 Molecular dynamics refinement and conformer processing

All simulations were run on Digital Research Alliance of Canada[6] clusters. Conformers selected for refinement by molecular dynamics simulations were the most upweighted structures from the energy landscape screen with $R_g$ < 85 Å. Each conformer was prepared for all-atom simulation using the GROMACS 2022.3 molecular dynamics package[10] with the CHARMM27 force field and TIP3P explicit water model. Each conformer was centered in a cubic water box with 1.0 nm padding, solvated with SPC216 water, and neutralized with $Na^+$ and $Cl^-$ ions to a final concentration of 150 mM.

Energy minimization was performed using the steepest descent algorithm until the maximum force was <$10^3$ kJ $mol^{-1}$ $nm^{-1}$. Equilibration was carried out for 200 ps under NPT conditions (298 K, 1 bar) using the leap-frog integrator with a 2 fs time step. Bonds to hydrogens were constrained with LINCS (order 6, one iteration), and water geometry with SETTLE. Long-range electrostatics were treated with PME (real-space cutoff 0.95 nm; Fourier grid spacing 0.12 nm), and van der Waals interactions with a 0.95 nm cutoff. Temperature was maintained with the V-rescale thermostat (τ = 0.1 ps; separate coupling groups for protein and solvent), and pressure with the Berendsen barostat (τ = 2 ps; isotropic coupling).

Production runs were performed for 4 ns total, split into two consecutive 2 ns simulations with the Parrinello-Rahman barostat (τ = 2 ps, isotropic, 1 bar), with coordinates saved every 100 ps. These short refinements at 298 K were used to enhance local compaction and side-chain packing; they were not intended to represent structures at equilibrium or provide a comprehensive conformational search. Importantly, we deliberately leveraged the known over-compaction bias of the CHARMM27 force field for IDPs[15] as a means of improving local packing during short refinements.

After simulations, trajectories were processed with GROMACS tools. Production runs were concatenated (trjcat) and reimaged to generate protein-centered trajectories without solvent or ions (trjconv). Conformers were extracted every 0.4 ns as PDB snapshots (trjconv -dump), yielding 10 conformers per trajectory. Standard trajectory analyses were also performed, including protein radius of gyration (gmx gyrate), backbone RMSD (gmx rms), and solvent accessible surface area (SASA) for the whole protein, backbone, and sidechains (gmx sasa). Refined conformers were then compiled to create a new pool for energy landscape screening and ensemble generation, which was also integrated into the IDPConformerGenerator pool.

**1.9 Clustering and structural analysis**

After applying the PRE-derived reweighting described in Eqs. S1–S4, the resulting conformer pool can be analyzed in terms of structural and thermodynamic quantities. In the following equations (S5–S11), we define how the radius of gyration, conformer probabilities, and pseudo-free energies were calculated for both individual conformers and structural clusters.

Clustering and radius of gyration ($R_g$) calculations were performed on full-length hTE and overlapping subsequences spanning one-half, one-quarter, and one-eighth of the sequence length. Pairwise distances between backbone $C_\alpha$ atoms were used to compute conformer similarity as in Tsangaris *et al.*[16], and hierarchical clustering was applied with the Ward linkage criterion using SciPy[17] (function scipy.cluster.hierarchy). Subsets of residues corresponding to each sequence window were isolated, and the radius of gyration was computed for each conformer within that window using only $C_\alpha$ coordinates[18]:

$$R_g = \sqrt{\frac{1}{N}\sum_{i=1}^{N}\left|\mathrm{r}_i - \mathrm{r}_{\mathrm{cm}}\right|^2} \tag{S5}$$

where N is the number of $C_\alpha$ atoms in the subsequence, $\mathbf{r}_i$ are the Cartesian coordinates ($x$,$y$,$z$) of the $C_\alpha$ atoms taken from the PDB structure, and $\mathbf{r}_{cm}$ is the center of mass, approximated as the positional average of all $C_\alpha$ atoms[18]:

$$\mathrm{r}_{\mathrm{cm}} = \frac{1}{N}\sum_{i=1}^{N}\mathrm{r}_i \tag{S6}$$

To determine the pseudo-free energy of each conformer the reweigthing factors must first be converted to probabilities. The probability of a single conformer $k$ being selected is defined as:

$$P_{\mathrm{conformer},k} = \frac{W_k}{\sum_{m=1}^{N_{\mathrm{conformer}}} W_m} \tag{S7}$$

where $W_k$ is the aggregate weight of conformer $k$, $N_{\mathrm{conformer}}$ is the total number of conformers in the pool, and $m$ is the summation index over all conformers.

Each conformer's pseudo-free energy $G^*_{\mathrm{conformer}}$ was estimated from its normalized probability across the full conformer pool using the Boltzmann relation[19]:

$$G^*_{\mathrm{conformer},k} = -RT\ln\left(P_{\mathrm{conformer},k}\right) \tag{S8}$$

where $P_{\mathrm{conformer}}$ is the normalized conformer probability from Eq. S7, $R$ is the gas constant, and $T$ is the absolute temperature.

To compare the relative stabilities of conformational clusters, we calculated cluster-level pseudo-free energies from the probabilities of their constituent conformers. The probability of a given cluster *c* is defined as the sum of the probabilities of all conformers assigned to that cluster:

$$P_{\text{cluster},c} = \sum_{k \in c} P_{\text{conformer},k} \tag{S9}$$

where *k* indexes all conformers belonging to cluster *c*, and $P_{\text{conformer},k}$ is the normalized probability of conformer *k* from Eq. S7.

The total probability of all conformers in the representative ensemble *e* is then:

$$P_{\text{ensemble},e} = \sum_{k \in e} P_{\text{conformer},k} \tag{S10}$$

The pseudo-free energy of cluster *c* was estimated from its relative probability using the Boltzmann relation[19]:

$$G^{*}_{\text{cluster},c} = -RT \ln\left(\frac{P_{\text{cluster},c}}{P_{\text{ensemble},e}}\right) \tag{S11}$$

where *R* is the gas constant and *T* is the absolute temperature.

**1.10 Dynamics measurements**

Longitudinal ($^{15}$N $T_1$) and transverse ($^{15}$N $T_2$) relaxation times were determined for uniformly $^{15}$N-labeled full-length hTE at 200 µM in 50 mM sodium phosphate buffer (pH 7.0) containing 50 mM NaCl at 10 °C. Pseudo-3D experiments were used with standard inversion-recovery ($T_1$)[20] and CPMG spin-echo ($T_2$)[21] pulse sequences . For $T_1$, 10 $^1$H-$^{15}$N HSQC spectra were acquired incrementally with relaxation delays ranging from 100 to 1,500 ms. For $T_2$, 10 spectra were acquired in an inter-leaved manner with delays ranging from 17 to 340 ms. Signal decays were fit to a mono-exponential decay function by least-squares analysis in CcpNmr[3] using default parameters. Relaxation rates ($R_1$ and $R_2$) are reported as the inverse of the fitted relaxation times, with errors estimated from fitting uncertainties.

#### 2. Supplementary Figures

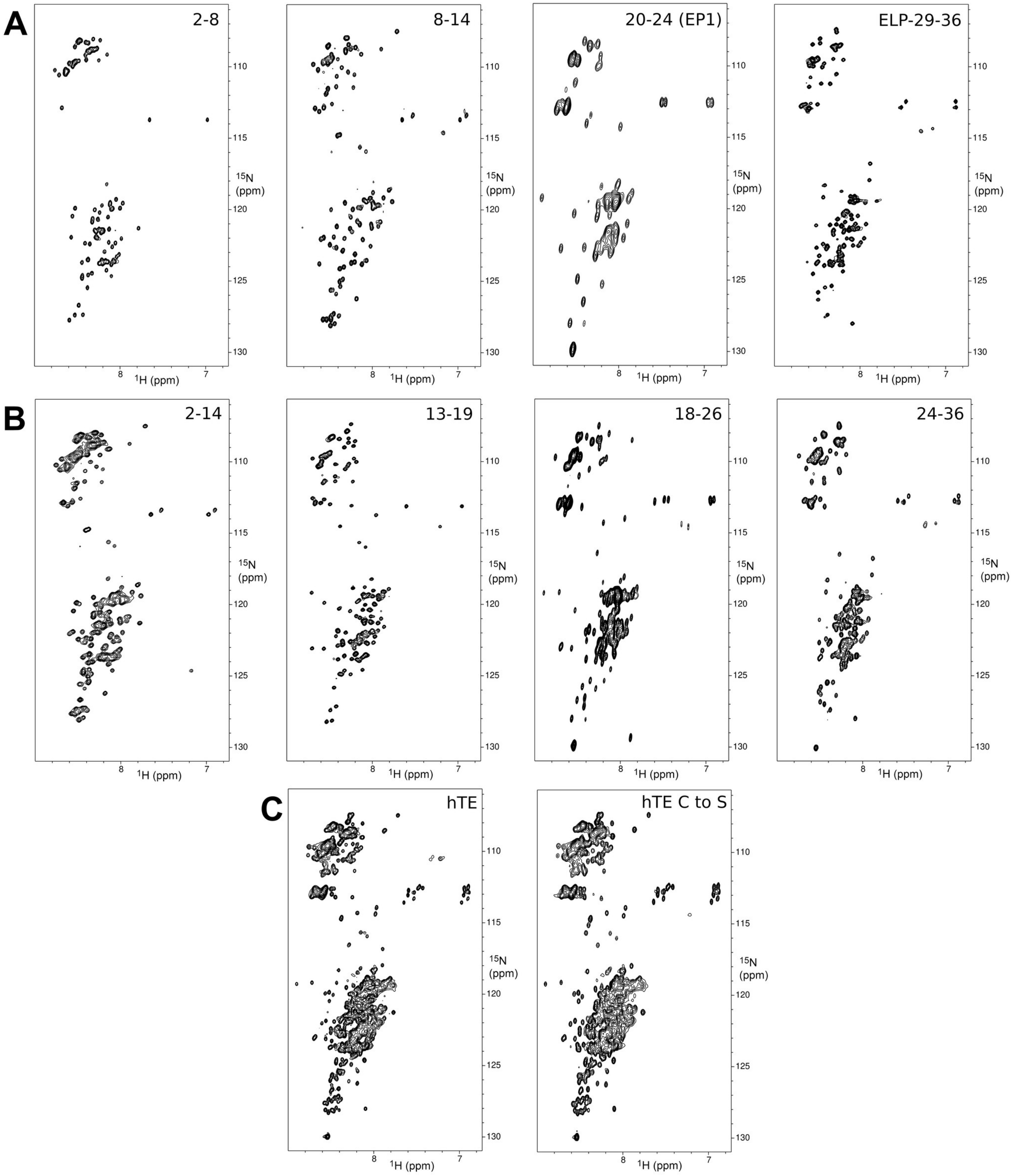

**SI Figure 1. $^{1}$H-$^{15}$N HSQC spectra of hTE constructs.**
(**A**) Short hTE fragments (~150 residues or fewer). (**B**) Medium-sized hTE fragments (~170 to 240 residues). (**C**) Full-length hTE WT (hTE) and C688S/C693S (hTE C to S) mutant. Amino acid sequences of hTE constructs are in SI Table 1.

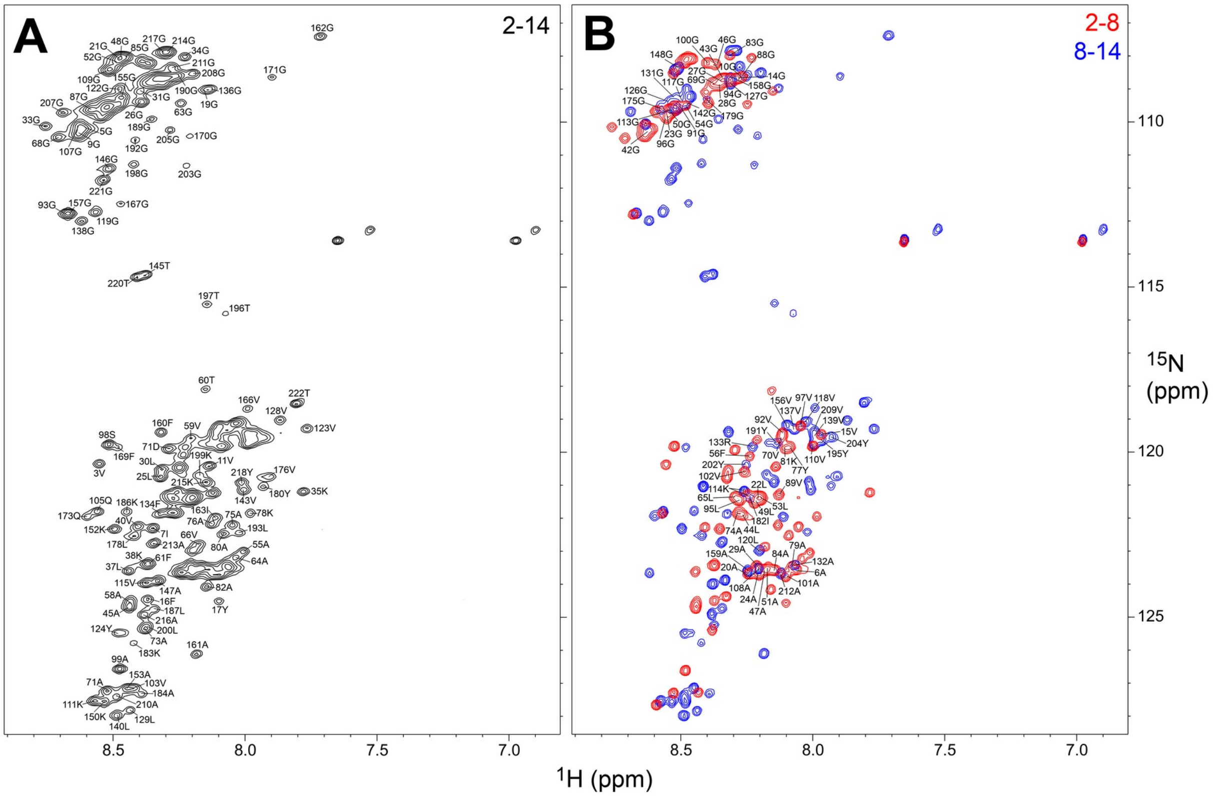

**SI Figure 2. Assigned $^{1}H$-$^{15}N$ HSQC spectra of overlapping hTE fragments.**
(**A**) Spectrum of the 2-14 domain construct. (**B**) Overlay of 2-8 and 8-14 domain construct spectra.

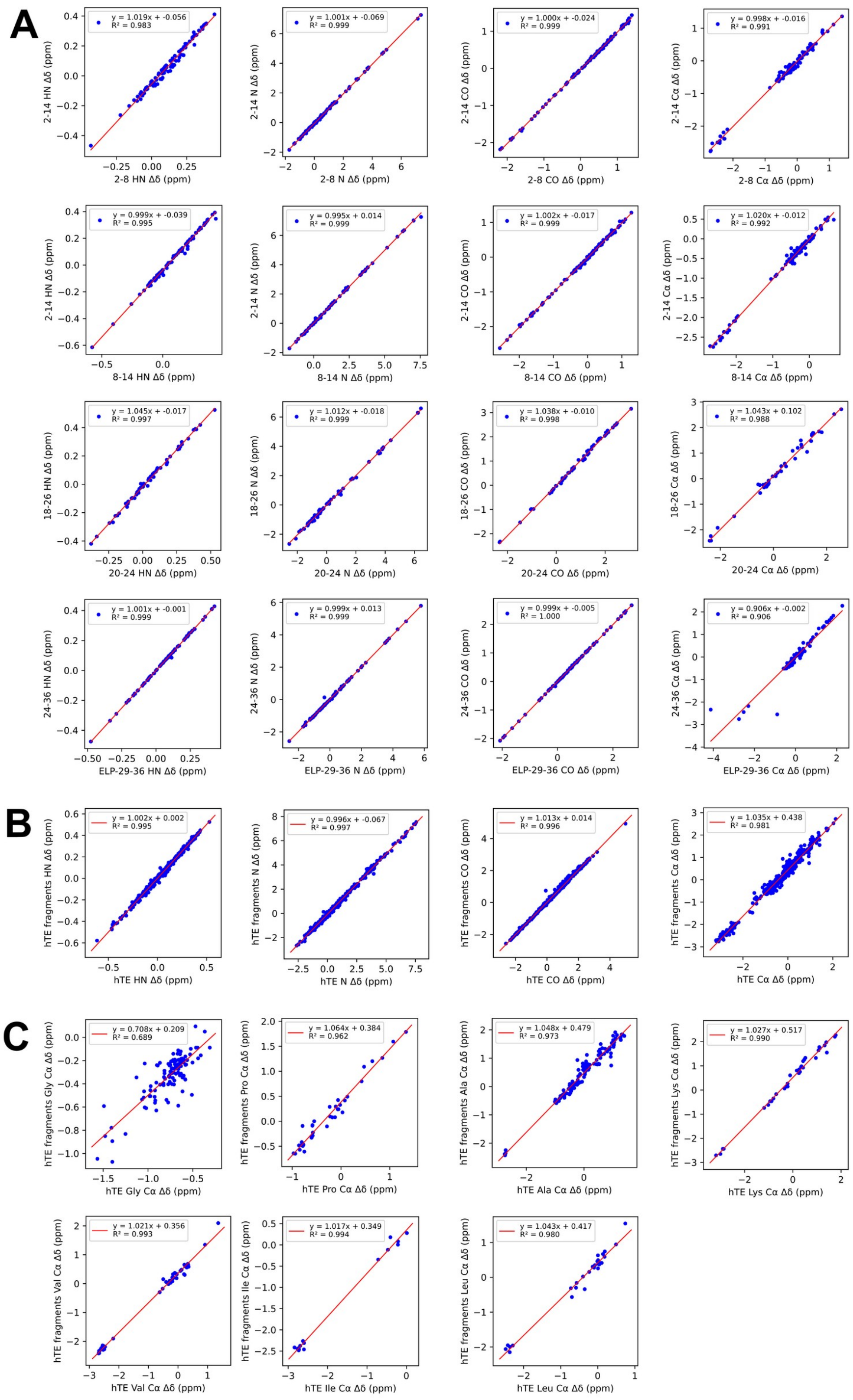

**SI Figure 3. Secondary chemical shift correlations.**
$^{1}$HN, $^{15}$N, $^{13}$CO, and $^{13}$Cα secondary chemical shifts (Δδ) for backbone nuclei are shown for overlapping sequence constructs. (**A**) Correlation between medium- and short-length fragments. (**B**) Correlation between all fragments and full-length

hTE. (**C**) Correlation between all fragments and full-length hTE for specific amino acid residue types. Secondary chemical shifts were calculated as the difference from random-coil values in Zhang *et al.*[22].

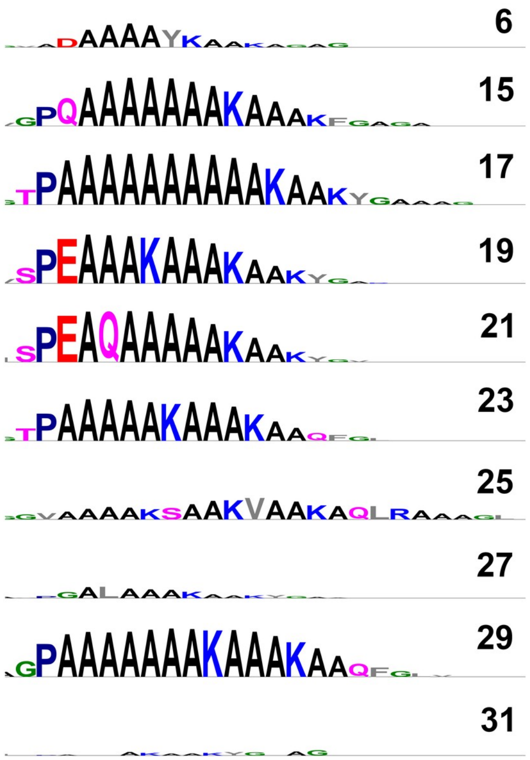

**SI Figure 4. Sequence-specific α-helical propensity of KA CLDs.**
Logo plots of CLD sequences, with the height of each amino acid proportional to α-helical propensity. Labels represent domain numbering from Figure 1. CLDs with prolines preceding polyalanine regions have significantly greater α-helical propensity. The total amount of α-helical content is also proportional to the length of the polyalanine region.

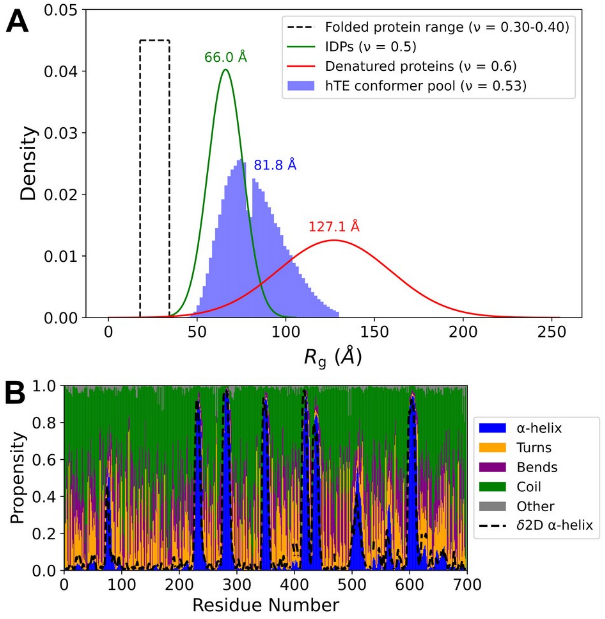

**SI Figure 5. Structural analysis of the initial hTE conformer pool.**
(**A**) Comparison of the radius of gyration ($R_g$) distribution of the ~ 200, 000 initial hTE conformers generated using IDP-ConformerGenerator to the size distributions expected for folded, IDPs or denatured proteins of the same chain length (698 amino acids). Expected $R_g$ values were scaled according to the Flory equation ($R_g = R_0 \cdot N^{\nu}$), where $R_0$ is a general proportionality constant (2.5 Å) for all protein types, $N$ is the chain length in amino acids, and $\nu$ is the scaling factor (0.30 to 0.4 for folded/globular proteins in a poor solvent; 0.5 for IDPs in an ideal solvent, 0.6 for denatured proteins in a good solvent)[23-25]. Standard deviations were approximated as $\sigma_{Rg} = f \cdot R_g$ , where $f$ is a fractional width (0.15 for IDPs in an ideal solvent; 0.25 for denatured proteins in a good solvent) based on $R_g$ distributions from simulations using force fields with different compaction tendencies described by Rauscher *et al*[15]. These size distributions are intended to be heuristic estimates rather than exact measures of conformer diversity. The initial pool of conformers had a mean $R_g$ of 81.8 Å, corresponding to a ν value of 0.53. (**B**) Back-calculated secondary structure from DSSP[26] shown as stacked bars. δ2D α-helical propensity from chemical shifts (dashed line) is in good agreement with α-helical content of the conformer pool.

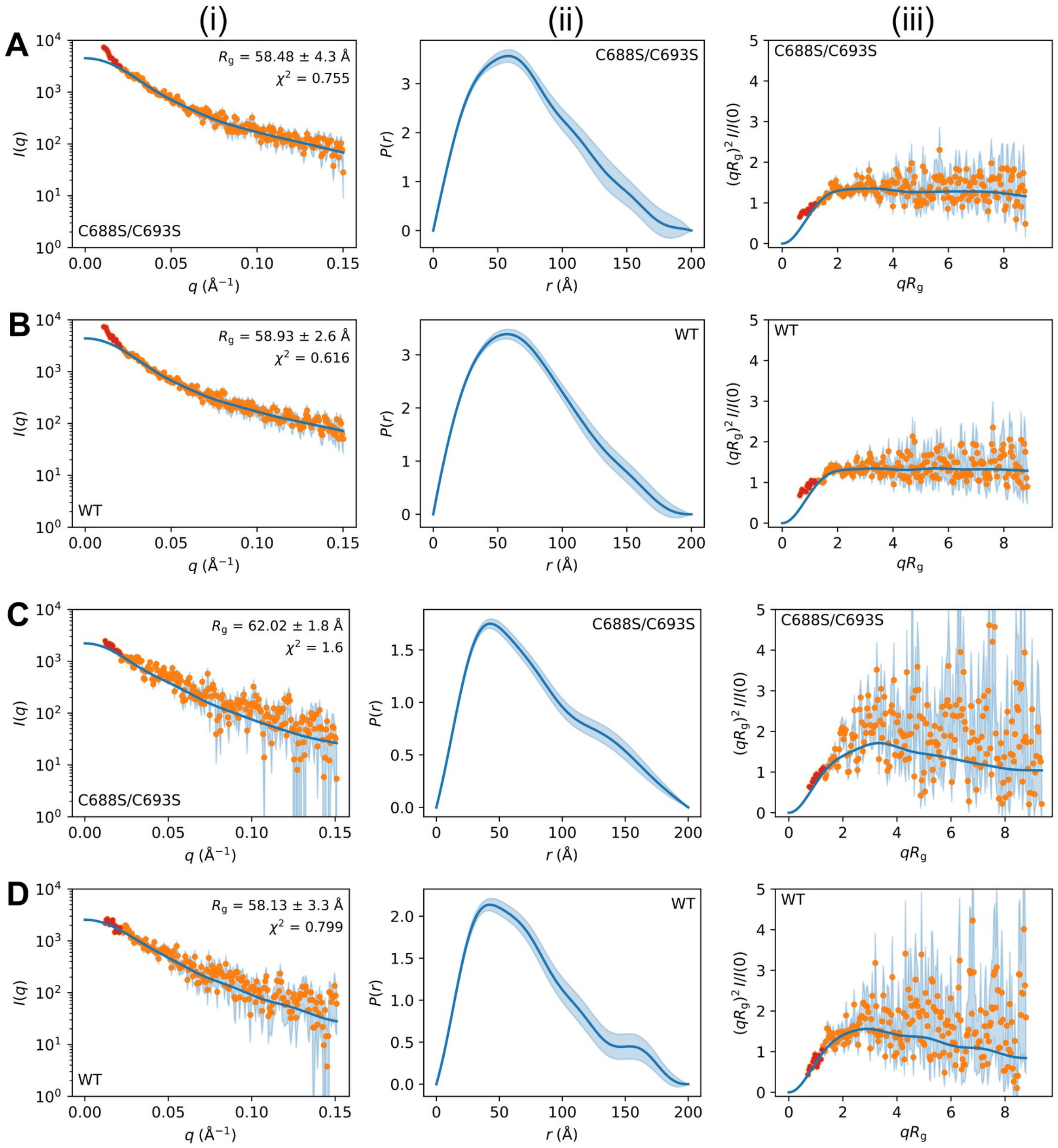

**SI Figure 6. Experimental SAXS analysis of hTE variants.**
SAXS data for (**A**) C688S/C693S at 200 µM, (**B**) WT at 200 µM, (**C**) C688S/C693S at 100 µM, and (**D**) WT at 100 µM in 50 mM sodium phosphate (pH 7) and 50 mM NaCl at 10 °C. Panels (i–iii), left to right: (i) scattering curve $I(q)$ vs $q$, (ii) distance distribution $P(r)$ from GNOM[11] using $D_{max}$= 200 Å, and (iii) dimensionless Kratky plot $(qR_g)^2I/I(0)$ vs $qR_g$, which rises slightly and plateaus without returning to 0—behavior indicative of a disordered ensemble[27-29]. Orange points denote $q$-regions without detectable interparticle effects; red points mark low-$q$ data showing a concentration-dependent upturn consistent with monomer–monomer interactions[27,28], which is less pronounced at 100 µM than 200 µM. The solid blue line is the GNOM fit; light blue shading indicates experimental uncertainties. Results from GNOM fits using $D_{max}$ values ranging from 150–240 Å are provided in SI Table 5.

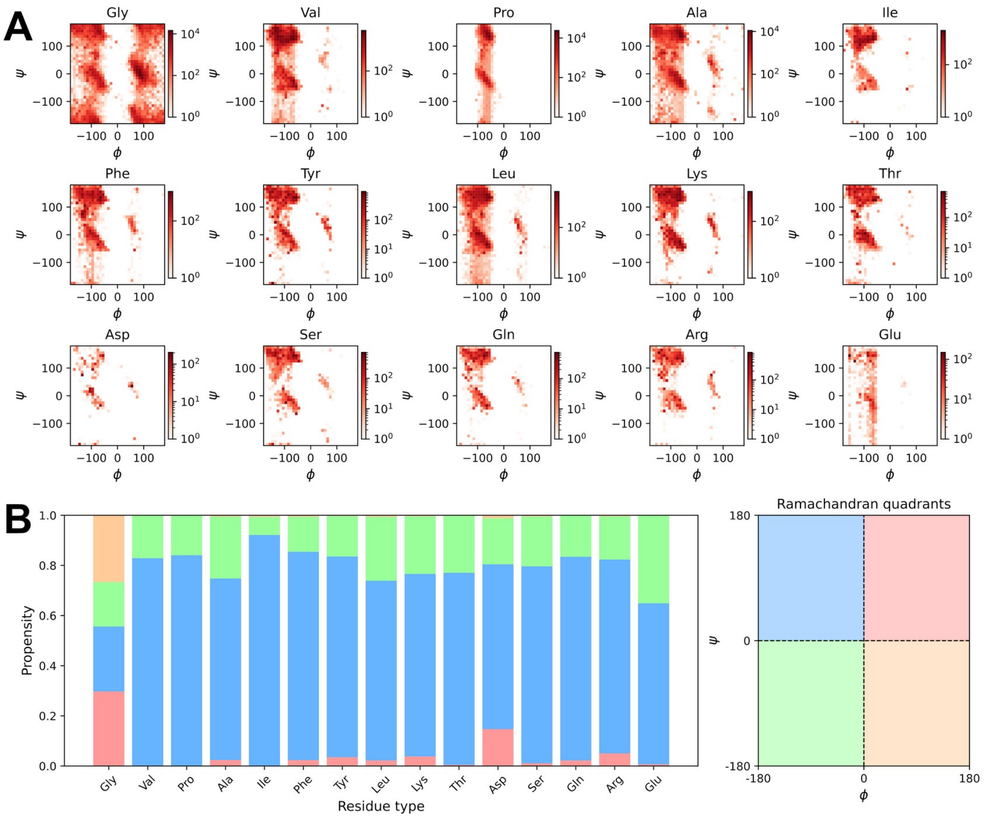

**SI Figure 7. Ramachandran space sampled by hTE conformers generated by IDPConformerGenerator.** (**A**) Heat maps of backbone torsion angles (φ, ψ) for each amino acid type. (**B**) Stacked bar graphs of reside-specific propensities to sample each quadrant of Ramachandran space color coded by region.

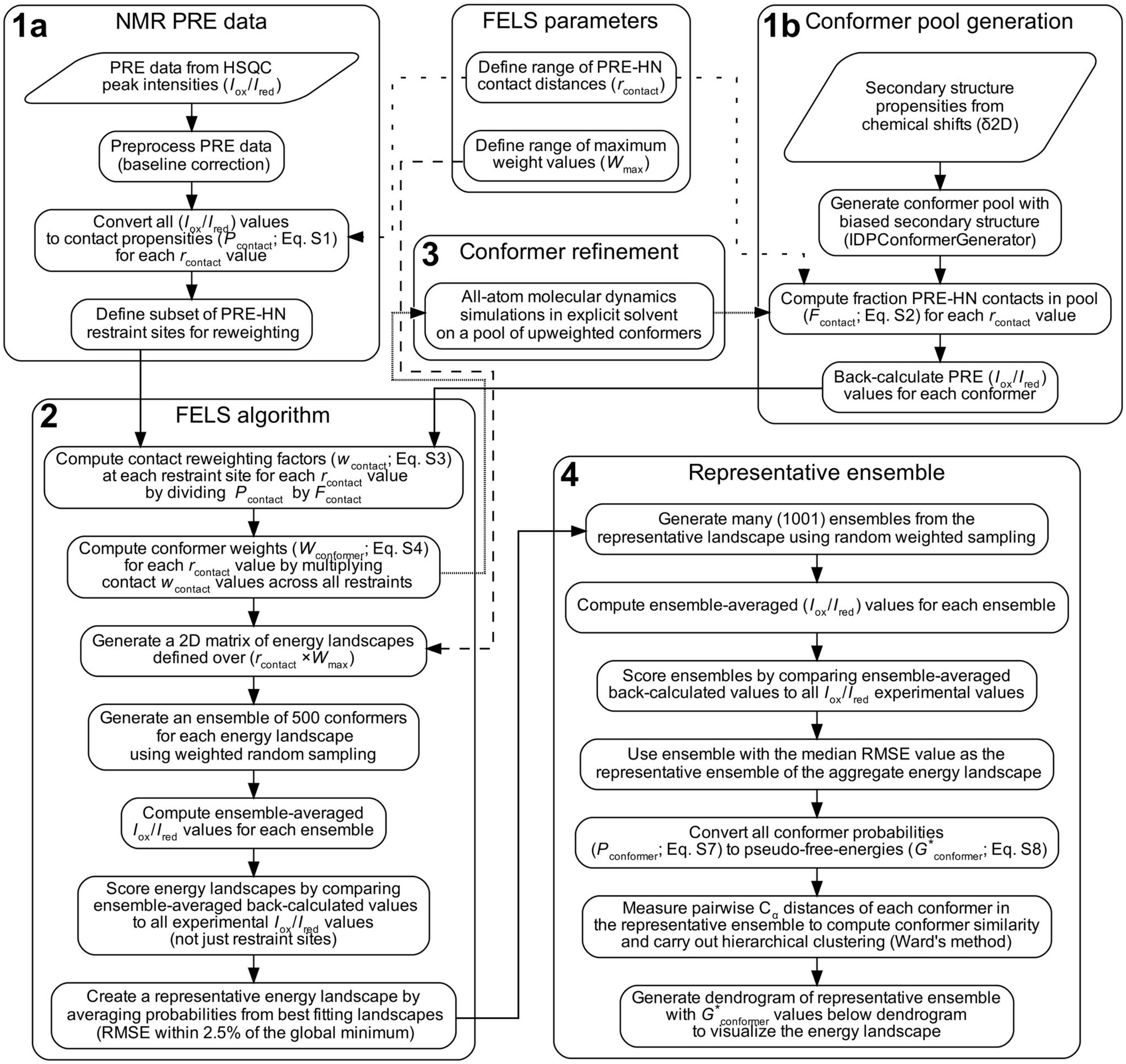

**SI Figure 8. Detailed FELS workflow.**

This schematic summarizes the full workflow from data preparation to ensemble analysis. Experimental PRE data are collected and processed to define contact propensities and restraint sites (**1a**), while an independent computational pipeline generates a large pool of conformers biased by secondary structure propensities derived from chemical shifts (**1b**). Core parameters are defined centrally, including the range of contact distances ($r_{contact}$) and maximum weight values ($W_{max}$), with short dashed arrows indicating where $r_{contact}$ is first applied in both PRE-derived contact propensities and conformer-based fraction contact calculations, and long dashed arrows indicating where $W_{max}$ governs construction of the two-dimensional energy landscape matrix. These inputs are integrated within the FELS algorithm (**2**) to generate energy landscapes and ensembles. An optional refinement pathway (**3**, stippled arrows) uses all-atom molecular dynamics simulations of upweighted conformers to improve structural realism and fitting to PRE data, with refined conformers fed back into the FELS algorithm. The final stage (**4**) focuses on generation and analysis of the representative ensemble, including structural clustering to identify dominant conformational states and energy mapping to relate ensemble populations to features of the underlying landscape.

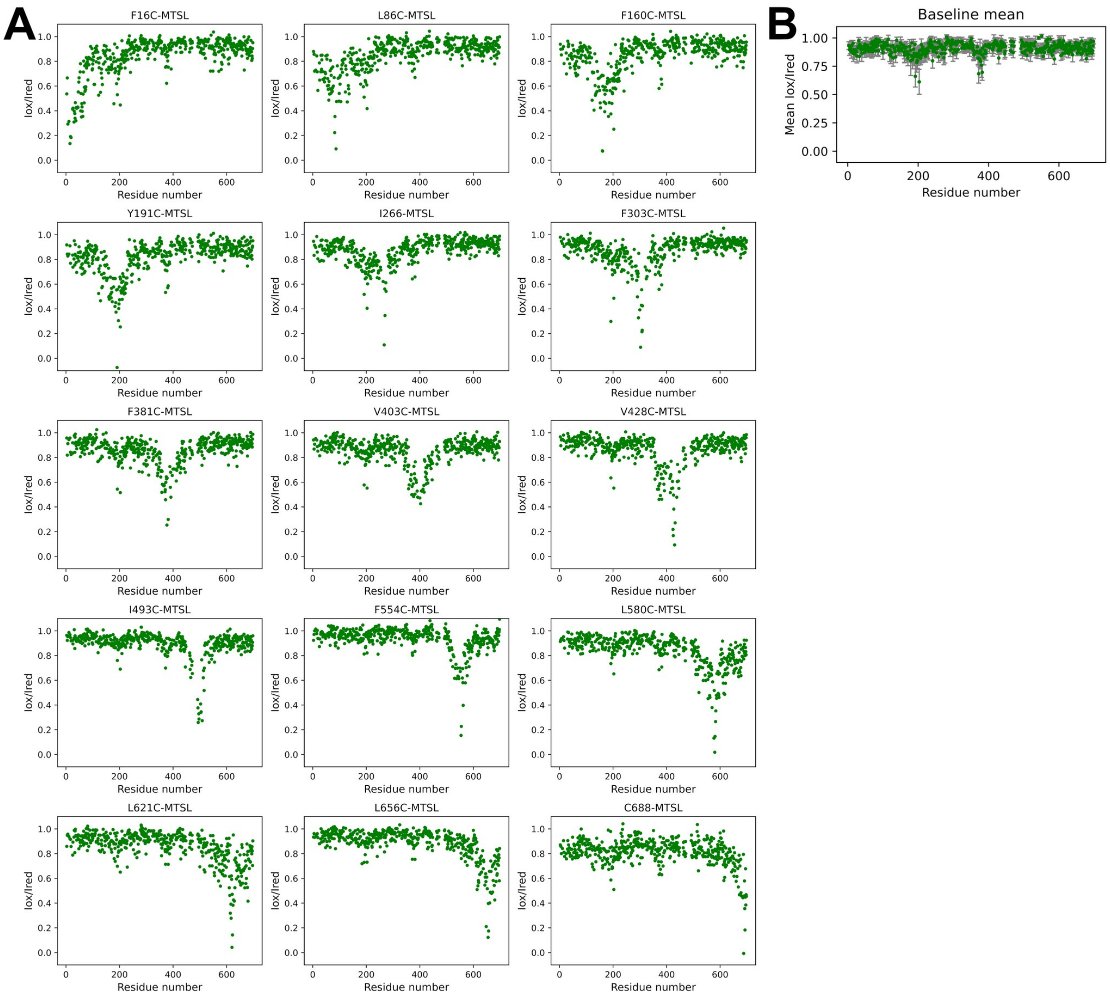

**SI Figure 9. Raw PRE data.**

(**A**) $^{1}$H-$^{15}$N HSQC peak intensity ratios ($I_{ox}/I_{red}$) plotted as a function of sequence position for each MTSL-labeled hTE mutant construct. (**B**) Mean upper baseline $I_{ox}/I_{red}$ values, calculated by averaging $I_{ox}/I_{red}$ values across all MTSL-labeled constructs for residues located more than 200 residues away from each label. Error bars represent the standard deviation from the mean.

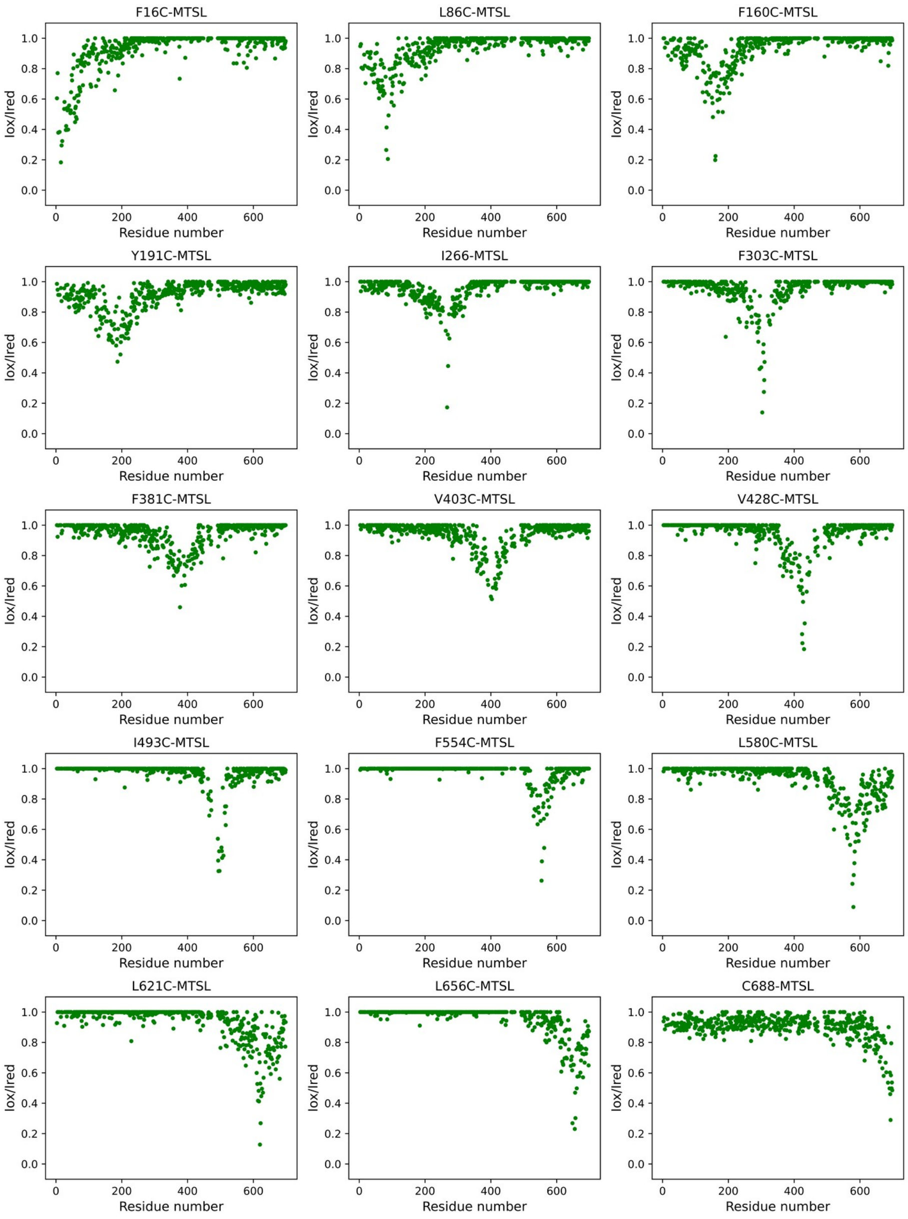

**SI Figure 10. Baseline-corrected PRE data.**
$^{1}H$-$^{15}N$ HSQC peak intensity ratios ($I_{ox}/I_{red}$) plotted as a function of sequence position for each MTSL-labeled hTE mutant construct after correction for intramolecular and site-specific relaxation effects. The baseline values shown in SI Figure 9B was subtracted from the raw PRE data in SI Figure 9A. $I_{ox}/I_{red}$ values greater than 1 were set to 1 to avoid introducing negative contact propensities.

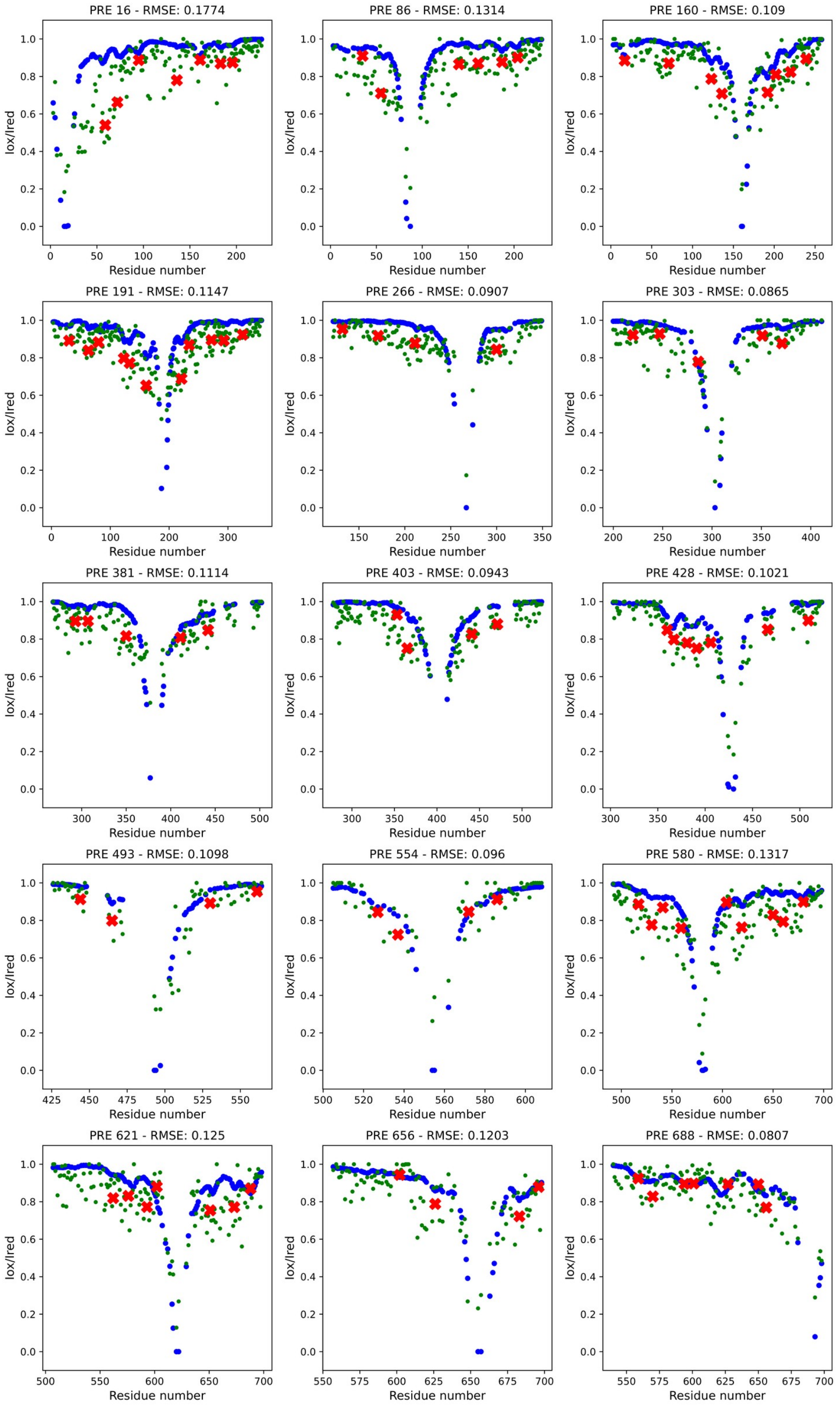

**SI Figure 11. Initial energy landscape screen fits for individual MTSL-labeled hTE sites.**
Plots show baseline-corrected experimental $I_{ox}/I_{red}$ values (small green circles) for each of the 15 MTSL-labeled hTE constructs, overlaid with back-calculated curves (larger blue circles) from the globally fitted representative ensemble using the IDPConformerGenerator pool of conformers described in SI Figure 5. Restraint sites used during ensemble fitting are marked with red Xs.

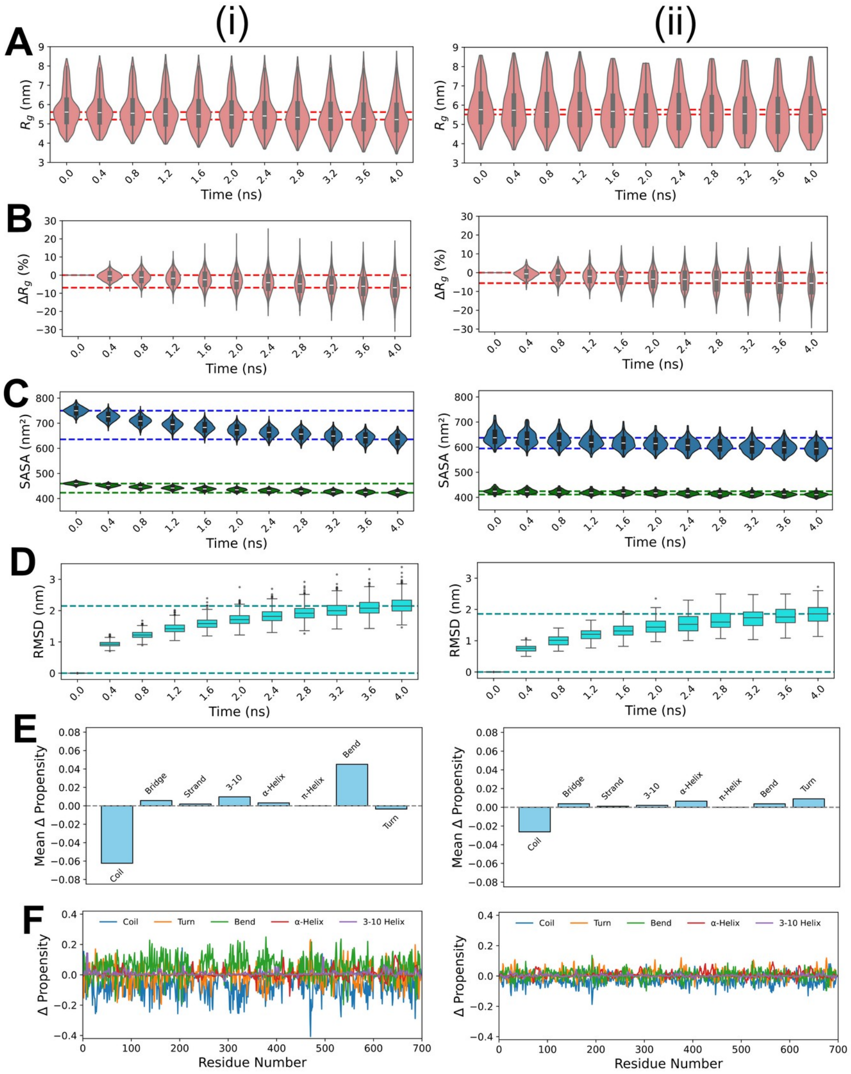

**SI Figure 12. Short all-atom MD refinement of upweighted conformers.**
(i) First round of MD refinement and (ii) second round of MD refinement for conformers upweighted in the initial energy landscape screen. Dashed parallel lines indicate the bounds of the mean values across the simulation trajectories. Violin plots show the change in conformer radius of gyration ($R_g$) during MD simulations as a function of (**A**) absolute size and (**B**) percent change. (**C**) Violin plots show the change in solvent-accessible surface area (SASA) during MD simulations, with blue representing sidechain SASA and green representing backbone SASA. (**D**) Box plots of backbone atom RMSD from the first time point of the trajectory. (**E**) Bar chart showing the mean DSSP[26] secondary structure propensity change from the first to the last time point of the trajectories. (**F**) Line chart showing the change in DSSP secondary structure as a function of sequence position.

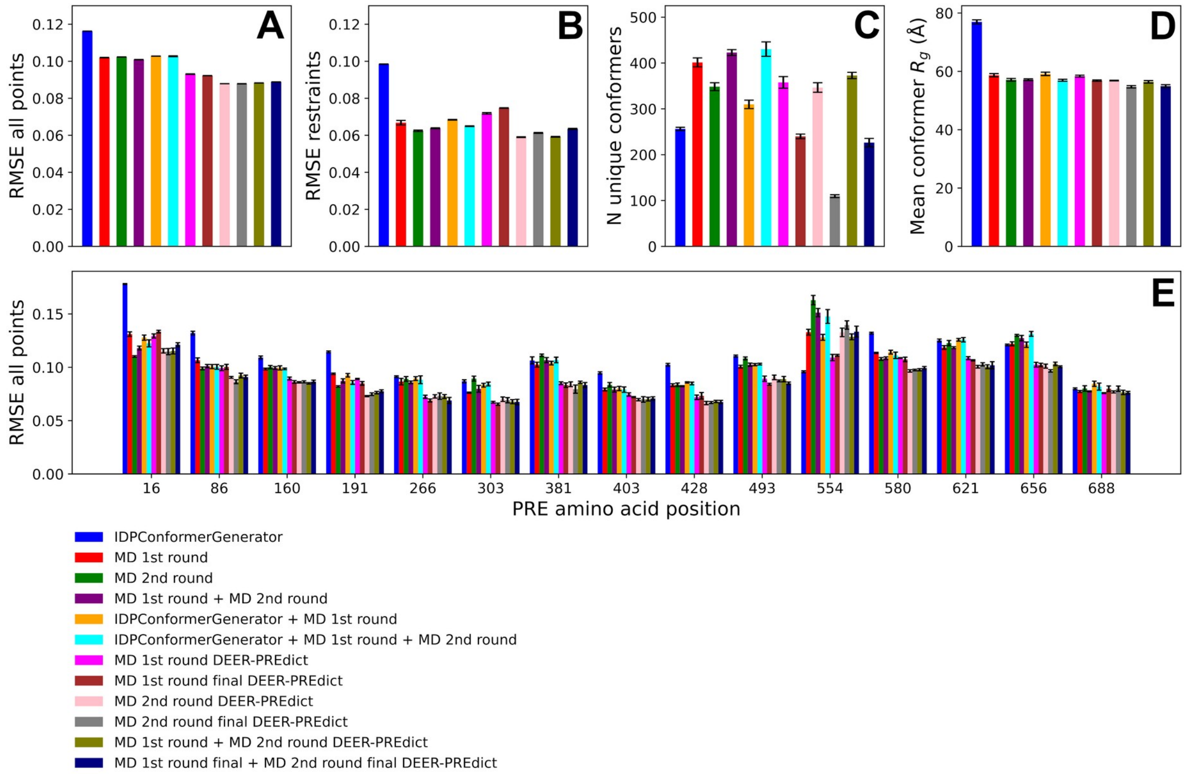

**SI Figure 13. Summary FELS runs using various conformer pool combinations.**
(**A**) RMSE of global PRE fits for each FELS run using different combinations of the initial IDPConformerGenerator pool, singly refined MD pool, and doubly refined MD pool, evaluated under both fixed-point and flexible PRE back-calculation models (DEER-PREdict)[14]. For MD pools, 10 snap shots were taken from each 4 ns trajectory to create a new pool unless indicated as "final", where only the final structure of each trajectory was used. (**B**) RMSE for restraint-site fits under the same conditions as in (**A**). (**C**) Ensemble diversity for each screened ensemble. (**D**) Mean $R_g$ values for final selected ensembles from each screen. (**E**) Comparison of single-site fits across restraint and non-restraint sites.

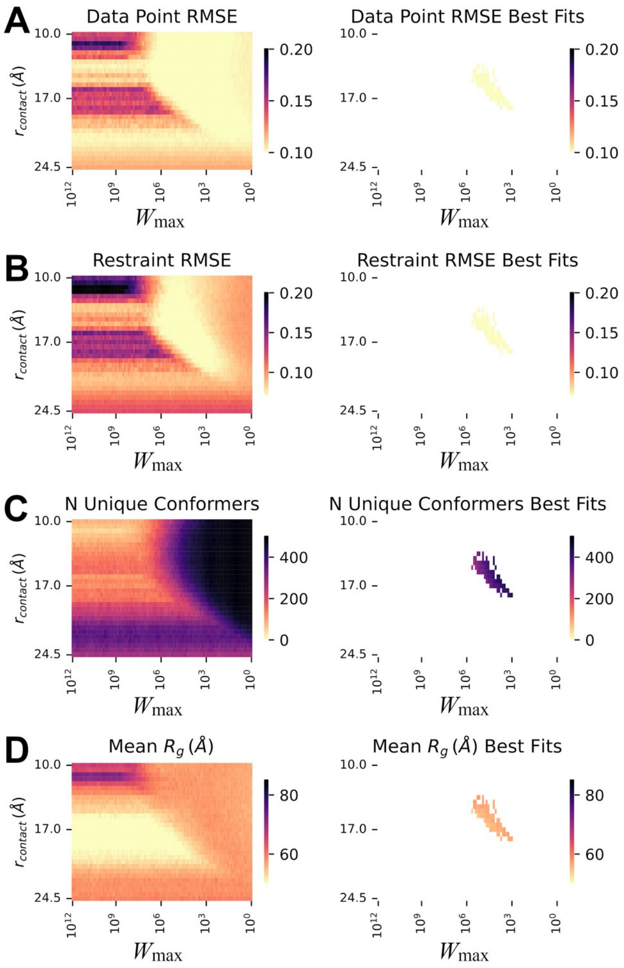

**SI Figure 14. MD-refined conformer pool energy landscape screen.**

Energy landscape screening was performed using the combined MD-refined conformer pool under the flexible PRE model to evaluate fits across the full landscape and for the best-fitting ensembles. For each panel, the left plot shows values across the entire landscape screen, while the right plot shows values for ensembles within 2.5% RMSE of the best fit for all data points. 2D heatmaps are shown, where each point represents a distinct energy landscape and its corresponding representative ensemble, displaying (**A**) RMSE across all PRE data points, (**B**) RMSE across restraint sites, (**C**) Ensemble diversity and (**D**) Mean radius of gyration ($R_g$).

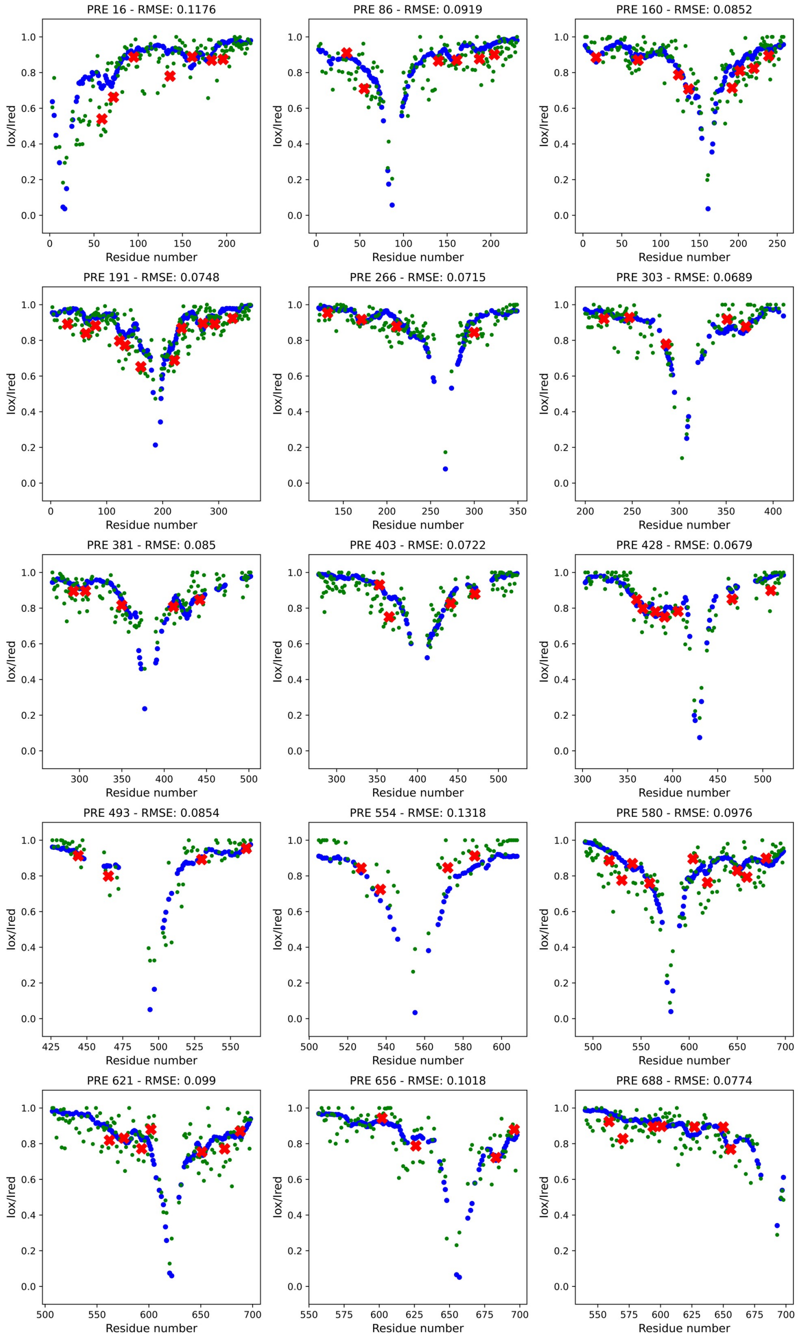

**SI Figure 15. Representative ensemble fits using MD-refined conformers and the flexible PRE model for individual MTSL-labeled hTE sites.**

Plots show baseline-corrected experimental $I_{ox}/I_{red}$ values (small green circles) for each of the 15 MTSL-labeled hTE constructs, overlaid with back-calculated curves (larger blue circles) from the globally fitted representative ensemble using the combined MD-refined conformer pool and the flexible PRE model implemented in DEER-PREdict[14]. Restraint sites used during ensemble fitting are marked with red Xs.

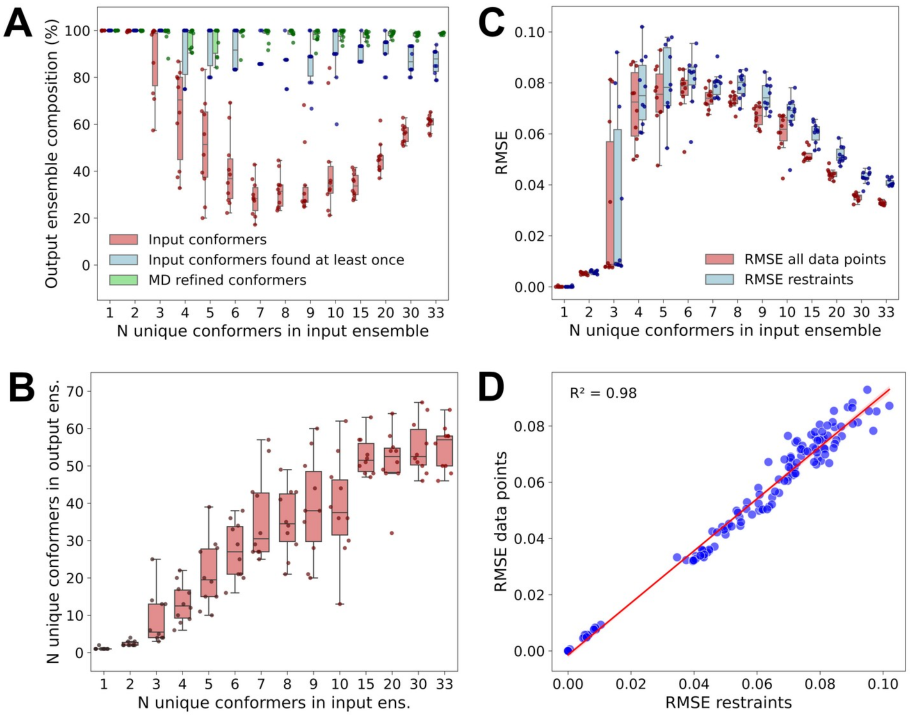

**SI Figure 16. Refined FELS proof-of-principle using MD-refined conformers and synthetic PRE data.** To validate the robustness of the energy landscape screen after refinement, synthetic PRE data were generated from input ensembles of hTE using random subsets of MD-refined structures, with each subset containing only the final structure from individual MD trajectories that had at least eight contacts at restraint sites used in the experimental screen. By including only a single structure per MD trajectory in the combined conformer pool, potential overlap was avoided, ensuring that percent recovery metrics reflected recovery at the trajectory level rather than counting multiple conformers from the same trajectory. Averaged back-calculated $I_{ox}/I_{red}$ curves from each subset served as input data for simulated energy landscape screens, which were run against a combined conformer pool containing both IDPConformerGenerator structures and single representative structures (final structure of each trajectory) from each MD-refined trajectory. (**A**) Composition of representative ensembles as a function of the number of unique conformers in the input ensemble. Box plots show the percentage of conformers in the output ensemble that were present in the input ensemble (red), the percentage of input conformers recovered at least once in the output ensemble (blue), and the percentage of MD-refined conformers recovered in the output ensemble (green). (**B**) Output ensemble diversity (number of unique conformers) as a function of the number of unique conformers in the input ensemble. (**C**) RMSE of global fits to synthetic input data as a function of the number of unique conformers in the input ensemble. (**D**) Correlation between global RMSE calculated from all data points and RMSE from restraint sites alone ($R^2 = 0.98$).

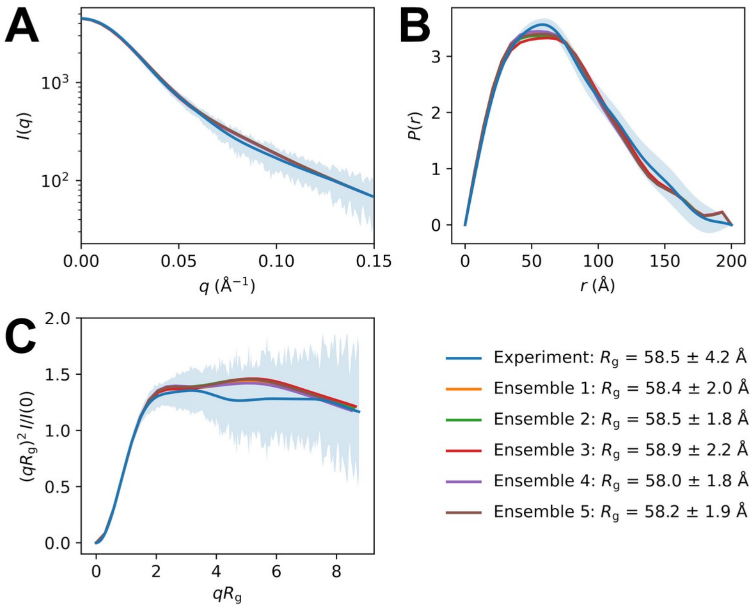

**SI Figure 17. Comparison of back-calculated SAXS data to experiment.**
(**A**) Comparison of CRYSOL[9] back-calculated SAXS curves for representative ensembles of hTE C688S/C693S generated from FELS runs against the MD-refined conformer pool using the flexible PRE model from DEER-PREdict[14]. Curves are normalized to the experimental scattering curve fitted with GNOM[11] (SI Figure 6A). (**B**) Comparison of pair-distance distributions $P(r)$ obtained from GNOM using $D_{\mathrm{max}}$ = 200 Å. (**C**) Dimensionless Kratky plot comparison. Experimental uncertainties are indicated by blue shading.

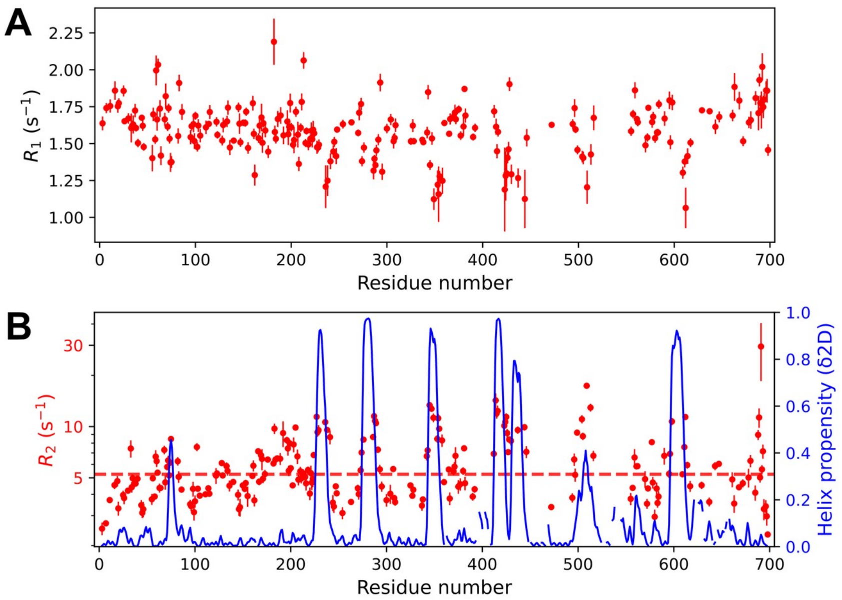

**SI Figure 18. Sequence-specific $^{15}$N backbone relaxation rates of hTE.**

(**A**) Longitudinal relaxation rates ($R_1$) and (B) transverse relaxation rates ($R_2$) measured for full-length hTE at 700 MHz and 10 °C in 50 mM sodium phosphate (pH 7) and 50 mM NaCl. Blue traces in (**B**) show δ2D-derived α-helix propensities overlaid on the $R_2$ data. The red dashed line indicates the mean $R_2$ value across the sequence. Error bars represent fitting errors obtained from exponential relaxation curve fitting.

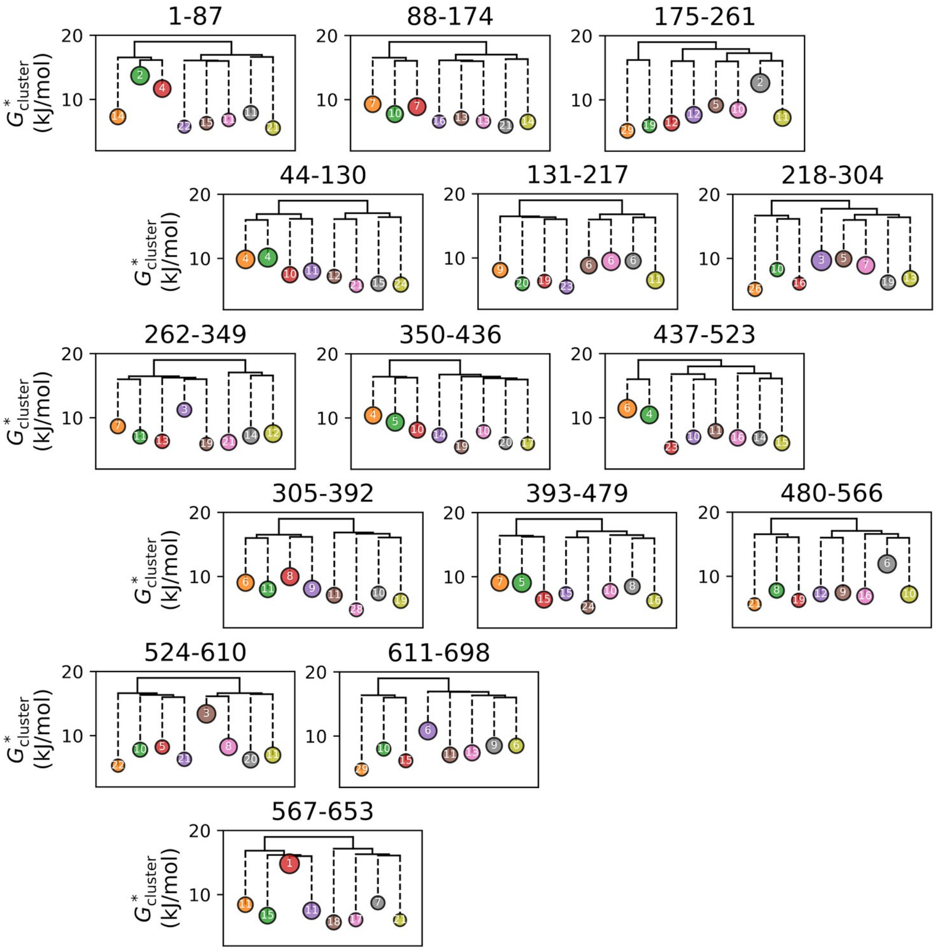

**SI Figure 19. Focused clustering analysis on overplapping sequence regions.**

Hierarchical clustering was performed on distinct regions of hTE, each approximately one-eighth the length of the sequence, using Euclidean distances derived from $C_\alpha$ distance matrices. Clustering was applied to the SAXS-validated representative ensemble generated from the combined MD-refined pool under the flexible PRE model. The simplified dendrograms show the Euclidean distance relationships between structural clusters (solid lines) and $G^*_{\mathrm{cluster}}$ values (dashed lines). The size of each circle is proportional to the mean $R_g$ of the cluster, and the number inside each circle indicates the percent fraction of the final ensemble.

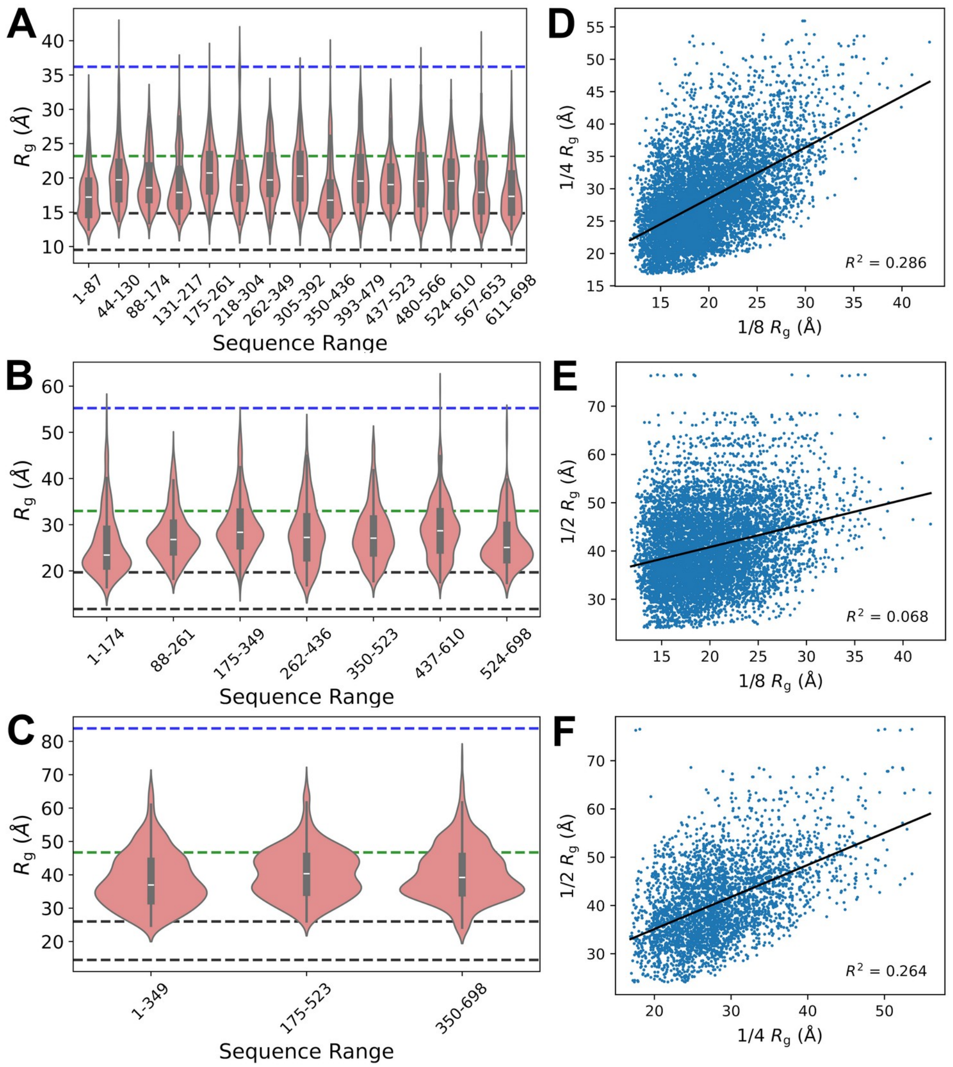

**SI Figure 20. Comparison of focused clustering across different overlapping sequence range sizes.**
Violin plots show the distribution of $R_g$ values for overlapping sequence ranges of different lengths calcuated from a representative ensemble generated from the combined MD-refined pool under the flexible PRE model from DEER-PREdict[14]. (**A**) Overlapping windows of 1/8 total hTE length. (**B**) Overlapping windows of 1/4 total hTE length.(**C**) Overlapping windows of 1/2 total hTE length. For panels A–C, dashed lines indicate the expected average size for a protein of similar length to hTE under different solvent conditions: blue for a denatured chain ($\nu$ = 0.6), green for a disordered chain ($\nu$ = 0.5), and two black lines indicating the range for globular proteins ($\nu$ between 0.3 and 0.4). The lack of correlation between the $R_g$ values of shorter and longer overlapping sequence regions is shown in (**D**) 1/4-length $R_g$ vs 1/8-length $R_g$ values.(**E**) 1/2-length $R_g$ vs 1/8-length $R_g$ values.(**F**) 1/2-length $R_g$ vs 1/4-length $R_g$ values.

#### 3. Supplementary Tables

**SI Table 1. Amino acid sequences of tropoelastin assignment constructs**

| Construct | BMRB ID | Amino acid sequence |
|---|---|---|
| hTE | 53477 | GGVPGAIPGGVPGGVFYPGAGLGALGGGALGPGGKPLKPVPGGLAGAGLGAGLGAFPAVTF<br>PGALVPGGVADAAAAYKAAKAGAGLGGVPGVGGLGVSAGAVVPQPGAGVKPGKVPGVGLP<br>GVYPGGVLPGARFPGVGVLPGVPTGAGVKPKAPGVGGAFAGIPGVGPFGGPQPGVPLGYPIK<br>APKLPGGYGLPYTTGKLPYGYGPGGVAGAAGKAGYPTGTGVGPQAAAAAAAKAAAKFGA<br>GAAGVLPGVGGAGVPGVPGAIPGIGGIAGVGTPAAAAAAAAAAKAAKYGAAAGLVPGGPG<br>FGPGVVGVPGAGVPGVGVPGAGIPVVPGAGIPGAAVPGVVSPEAAAKAAAKAAKYGARPG<br>VGVGGIPTYGVGAGGFPGFGVGVGGIPGVAGVPGVGGVPGVGGVPGVGISPEAQAAAAAKA<br>AKYGVGTPAAAAAKAAAKAAQFGLVPGVGVAPGVGVAPGVGVAPGVGLAPGVGVAPGVGV<br>APGVGVAPGIGPGGVAAAAKSAAKVAAKAQLRAAAGLGAGIPGLGVGVGVPGLGVGAGVP<br>GLGVGAGVPGFGAVPGALAAAKAAKYGAAVPGVLGGLGALGGVGIPGGVVGAGPAAAAA<br>AAKAAAKAAQFGLVGAAGLGGLGVGGLGVPGVGGLGGIPPAAAAKAAKYGAAGLGGVLG<br>GAGQFPLGGVAARPGFGLSPIFPGGACLGKACGRKRK |
| 2-8 | 53472 | GVPGAIPGGVPGGVFYPGAGLGALGGGALGPGGKPLKPVPGGLAGAGLGAGLGAFPAVTFP<br>GALVPGGVADAAAAYKAAKAGAGLGGVPGVGGLGVSAGAVVPQPGAGVKPGKVP |
| 8-14 | 27236 | GAVVPQPGAGVKPGKVPGVGLPGVYPGGVLPGARFPGVGVLPGVPTGAGVKPKAPGVGGA<br>FAGIPGVGPFGGPQPGVPLGYPIKAPKLPGGYGLPYTTGKLPYGYGPGGVAGAAGKAGYPTG<br>T |
| 2-14 | 53473 | GVPGAIPGGVPGGVFYPGAGLGALGGGALGPGGKPLKPVPGGLAGAGLGAGLGAFPAVTFP<br>GALVPGGVADAAAAYKAAKAGAGLGGVPGVGGLGVSAGAVVPQPGAGVKPGKVPGVGLP<br>GVYPGGVLPGARFPGVGVLPGVPTGAGVKPKAPGVGGAFAGIPGVGPFGGPQPGVPLGYPIK<br>APKLPGGYGLPYTTGKLPYGYGPGGVAGAAGKAGYPTGT |
| 13-19 | 53474 | GGYGLPYTTGKLPYGYGPGGVAGAAGKAGYPTGTGVGPQAAAAAAAKAAAKFGAGAAGV<br>LPGVGGAGVPGVPGAIPGIGGIAGVGTPAAAAAAAAAAKAAKYGAAAGLVPGGPGFGPGVV<br>GVPGAGVPGVGVPGAGIPVVPGAGIPGAAVPGVVSPEAAAKAAAKAAKYGARPGVG |
| 20-24 (EP1) | N/A | FPGFGVGVGGIPGVAGVPGVGGVPGVGGVPGVGISPEAQAAAAAKAAKYGVGTPAAAAAK<br>AAAKAAQFGLVPGVGVAPGVGVAPGVGVAPGVGLAPGVGVAPGVGVAPGVGVAPAIGP |
| 18-26 | 53475 | AGVPGVGVPGAGIPVVPGAGIPGAAVPGVVSPEAAAKAAAKAAKYGARPGVGVGGIPTYGV<br>GAGGFPGFGVGVGGIPGVAGVPGVGGVPGVGGVPGVGISPEAQAAAAAKAAKYGVGTPAA<br>AAAKAAAKAAQFGLVPGVGVAPGVGVAPGVGVAPGVGLAPGVGVAPGVGVAPGVGVAPGI<br>GPGGVAAAAKSAAKVAAKAQLRAAAGLGAGIPGLGVGVGVPGLGVGA |
| 24-36 | 53476 | GVGLAPGVGVAPGVGVAPGVGVAPGIGPGGVAAAAKSAAKVAAKAQLRAAAGLGAGIPGL<br>GVGVGVPGLGVGAGVPGLGVGAGVPGFGAVPGALAAAKAAKYGAAVPGVLGGLGALGGV<br>GIPGGVVGAGPAAAAAAAKAAAKAAQFGLVGAAGLGGLGVGGLGVPGVGGLGGIPPAAAA<br>KAAKYGAAGLGGVLGGAGQFPLGGVAARPGFGLSPIFPGGACLGKACGRKRK |
| ELP-29-36 | N/A | AAAAAKAAKYGAGVPGVGVPGVGVPGVGVPGVGVPGVGVPGVGVPGVGAGPAAAAAAA<br>KAAAKAAQFGLVGAAGLGGLGVGGLGVPGVGGLGGIPPAAAAKAAKYGAAGLGGVLGGA<br>GQFPLGGVAARPGFGLSPIFPGGACLGKACGRKRK |

**SI Table 2. Chemical shift assignments of full-length hTE and fragments**

| Residue | | $^{13}C_{\alpha}$ | | $^{13}CO$ | | $^{15}N$ | | $^{1}HN$ | | $H_{\alpha}$ | $C_{\beta}$ | Source |
|---|---|---|---|---|---|---|---|---|---|---|---|---|
| # | AA | hTE | Frag. | hTE | Frag. | hTE | Frag. | hTE | Frag. | Frag. | Frag. | Frag. |
| 2 | G | - | 43.0 | 173.7 | 182.0 | - | - | - | - | - | - | - |
| 3 | V | - | 60.0 | 174.8 | 174.7 | 121.6 | 120.5 | 8.35 | 8.57 | 4.30 | 32.3 | 2-8 |
| 4 | P | - | 63.6 | - | 177.5 | - | 140.2 | - | - | 4.38 | 31.9 | 2-8 |
| 5 | G | - | 45.1 | - | 173.7 | - | 110.2 | - | 8.62 | 3.93 | - | 2-8 |
| 6 | A | - | 52.2 | 177.6 | 177.6 | 123.7 | 123.6 | 8.11 | 8.07 | 4.21 | 19.4 | 2-8 |
| 7 | I | 58.3 | 58.5 | 174.8 | 174.8 | 122.4 | 122.3 | 8.34 | 8.35 | 4.26 | 38.5 | 2-8 |
| 8 | P | - | 63.7 | - | 177.5 | - | 141.2 | - | - | 4.31 | 31.9 | 2-8 |
| 9 | G | - | 45.3 | 174.7 | 174.7 | - | 110.4 | - | 8.64 | 3.75 | - | 2-8 |
| 10 | G | 44.6 | 45.0 | 173.6 | 173.7 | 108.8 | 108.7 | 8.29 | 8.30 | 3.97 | - | 2-8 |
| 11 | V | 59.5 | 59.8 | 174.7 | 174.7 | 120.5 | 120.4 | 8.14 | 8.14 | 4.26 | 32.7 | 2-8 |
| 12 | P | - | 63.6 | - | 177.5 | - | 139.6 | - | - | 4.40 | 31.9 | 2-8 |
| 13 | G | - | 45.5 | - | 174.7 | - | 110.4 | - | 8.63 | 3.74 | - | 2-8 |
| 14 | G | 44.8 | 45.2 | 173.7 | 173.7 | 108.9 | 108.8 | - | 8.32 | 3.97 | - | 2-8 |
| 15 | V | 61.8 | 62.1 | 175.3 | 175.3 | 119.6 | 119.4 | 7.97 | 7.98 | 3.95 | 33.0 | 2-8 |
| 16 | F | 57.0 | 57.4 | 174.4 | 174.4 | 124.6 | 124.4 | 8.37 | 8.38 | 4.51 | 40.1 | 2-8 |
| 17 | Y | 54.8 | 55.2 | 173.4 | 173.5 | 124.6 | 124.5 | 8.11 | 8.11 | 4.57 | 38.5 | 2-8 |
| 18 | P | - | 63.5 | 177.3 | 177.4 | - | 136.7 | - | - | 4.18 | 31.9 | 2-8 |
| 19 | G | - | 45.2 | 173.9 | 174.0 | 109.0 | 109.1 | 8.14 | 8.17 | 3.93 | - | 2-8 |
| 20 | A | - | 52.7 | 178.3 | 178.3 | 123.7 | 123.7 | 8.24 | 8.26 | 4.31 | 19.4 | 2-8 |
| 21 | G | - | 45.4 | - | 174.4 | 108.2 | 108.2 | - | 8.49 | 3.84 | - | 2-8 |
| 22 | L | - | 55.3 | - | 178.2 | - | 121.4 | - | 8.21 | 4.34 | 42.2 | 2-8 |
| 23 | G | - | 45.5 | - | 174.2 | - | 109.8 | - | 8.56 | 3.73 | - | 2-8 |
| 24 | A | - | 52.7 | 178.1 | 178.2 | 123.8 | 123.7 | - | 8.21 | 4.11 | 19.4 | 2-8 |
| 25 | L | 55.0 | 55.3 | 178.2 | 178.2 | 120.9 | 120.8 | 8.32 | 8.33 | 4.24 | 42.0 | 2-8 |
| 26 | G | 45.1 | 45.6 | 174.8 | 174.8 | 109.5 | 109.4 | 8.40 | 8.40 | 3.76 | - | 2-8 |
| 27 | G | - | 45.3 | 174.8 | 174.8 | 108.8 | 108.8 | 8.35 | 8.37 | - | - | 2-8 |
| 28 | G | 44.8 | 45.3 | 173.8 | 173.8 | 109.0 | 109.0 | 8.35 | 8.37 | 3.94 | - | 2-8 |
| 29 | A | 52.1 | 52.7 | 177.5 | 177.5 | 123.6 | 123.5 | 8.22 | 8.22 | 4.12 | 19.4 | 2-8 |
| 30 | L | 54.2 | 54.4 | 177.6 | 177.7 | 120.7 | 120.6 | 8.32 | 8.32 | 4.37 | 43.2 | 2-8 |
| 31 | G | 44.0 | 44.7 | 172.1 | 172.3 | 109.2 | 109.2 | 8.36 | 8.43 | 4.05 | - | 2-8 |
| 32 | P | 63.6 | 64.0 | 178.1 | 178.1 | - | - | - | - | 4.38 | 31.8 | 2-8 |
| 33 | G | 44.9 | 45.3 | 174.9 | 175.0 | 110.3 | 110.2 | 8.77 | 8.78 | 3.78 | - | 2-8 |
| 34 | G | 45.0 | 45.6 | 173.5 | 173.5 | 108.1 | 108.1 | 8.25 | 8.25 | 3.90 | - | 2-8 |
| 35 | K | 53.4 | 53.9 | 174.2 | 174.2 | 121.3 | 121.3 | 7.77 | 7.80 | 4.47 | 32.4 | 2-8 |
| 36 | P | 62.7 | 63.0 | 176.9 | 176.9 | - | 136.4 | - | - | 4.48 | 32.0 | 2-8 |
| 37 | L | 54.8 | 55.2 | 177.2 | 177.2 | 123.6 | 123.7 | 8.45 | 8.45 | 4.08 | 42.5 | 2-8 |
| 38 | K | 53.6 | 53.9 | 174.2 | 174.1 | 123.5 | 123.4 | 8.36 | 8.38 | 4.42 | - | 2-8 |
| 39 | P | 62.5 | 62.8 | 176.7 | 176.7 | - | 137.0 | - | - | 4.45 | 32.0 | 2-8 |
| 40 | V | 59.5 | 59.8 | 174.8 | 174.8 | 122.3 | 122.2 | 8.40 | 8.41 | 4.22 | 32.3 | 2-8 |
| 41 | P | - | 63.6 | - | 177.5 | - | 140.2 | - | - | - | 32.0 | 2-8 |

| Residue | | $^{13}C_{\alpha}$ | | $^{13}CO$ | | $^{15}N$ | | $^{1}HN$ | | $H_{\alpha}$ | $C_{\beta}$ | Source |
|---|---|---|---|---|---|---|---|---|---|---|---|---|
| # | AA | hTE | Frag. | hTE | Frag. | hTE | Frag. | hTE | Frag. | Frag. | Frag. | Frag. |
| 42 | G | - | 45.4 | - | 174.7 | - | 110.2 | - | 8.63 | 3.74 | - | 2-8 |
| 43 | G | 44.8 | 44.9 | 174.2 | 174.3 | 108.8 | 108.6 | - | 8.36 | 3.75 | - | 2-8 |
| 44 | L | 54.8 | 55.3 | 177.6 | 177.6 | 121.9 | 121.9 | 8.28 | 8.28 | 4.33 | 42.4 | 2-8 |
| 45 | A | 52.6 | 52.9 | 178.4 | 178.4 | 124.8 | 124.7 | 8.44 | 8.45 | 4.18 | 19.0 | 2-8 |
| 46 | G | 44.9 | 45.3 | 174.1 | 174.2 | 108.3 | 108.3 | 8.37 | 8.38 | 3.73 | - | 2-8 |
| 47 | A | - | 52.7 | - | 178.3 | 123.8 | 123.6 | 8.17 | 8.19 | 4.31 | 19.4 | 2-8 |
| 48 | G | - | 45.4 | - | 174.5 | - | 108.1 | - | 8.48 | 3.84 | - | 2-8 |
| 49 | L | - | 55.3 | - | 178.2 | - | 121.5 | - | 8.23 | 4.16 | 42.2 | 2-8 |
| 50 | G | - | 45.4 | - | 174.1 | - | 109.7 | - | 8.54 | 3.68 | - | 2-8 |
| 51 | A | - | 52.6 | - | 178.3 | - | 123.6 | - | 8.17 | 4.21 | 19.3 | 2-8 |
| 52 | G | - | 45.4 | - | 174.4 | - | 108.3 | - | 8.50 | 3.95 | - | 2-8 |
| 53 | L | - | 55.4 | 178.0 | 178.0 | - | 121.4 | - | 8.20 | 4.23 | 42.2 | 2-8 |
| 54 | G | 44.8 | 45.2 | 173.4 | 173.5 | 109.5 | 109.5 | 8.47 | 8.49 | 3.78 | - | 2-8 |
| 55 | A | 51.8 | 52.3 | 177.0 | 177.0 | 123.1 | 123.1 | 8.01 | 8.03 | 4.03 | 19.5 | 2-8 |
| 56 | F | 55.3 | 55.6 | 173.7 | 173.7 | 120.2 | 120.1 | 8.24 | 8.25 | 4.64 | 39.0 | 2-8 |
| 57 | P | 62.7 | 63.1 | 176.3 | 176.4 | - | - | - | - | 4.36 | 32.0 | 2-8 |
| 58 | A | 52.0 | 52.5 | 177.8 | 177.8 | 124.7 | 124.5 | 8.43 | 8.44 | 4.32 | 19.2 | 2-8 |
| 59 | V | 61.8 | 62.1 | 176.0 | 176.0 | 119.7 | 119.5 | 8.21 | 8.21 | 4.03 | 32.8 | 2-8 |
| 60 | T | 61.4 | 61.4 | 173.5 | 173.5 | 118.2 | 118.0 | 8.16 | 8.16 | 4.09 | 70.0 | 2-8 |
| 61 | F | 55.1 | 55.3 | 173.9 | 174.0 | 123.5 | 123.4 | 8.38 | 8.37 | 4.71 | 39.1 | 2-8 |
| 62 | P | 63.4 | 63.7 | 177.4 | 177.5 | - | - | - | - | 4.39 | 31.8 | 2-8 |
| 63 | G | 44.8 | 45.3 | 173.7 | 173.8 | 109.5 | 109.5 | 8.24 | 8.26 | 3.82 | - | 2-8 |
| 64 | A | 52.1 | 52.5 | 177.6 | 177.6 | 123.3 | 123.2 | 8.05 | 8.05 | 4.27 | 19.5 | 2-8 |
| 65 | L | 54.5 | 54.9 | 177.1 | 177.1 | 121.4 | 121.3 | 8.32 | 8.30 | 4.14 | 42.3 | 2-8 |
| 66 | V | 59.4 | 59.7 | 174.4 | 174.4 | 123.0 | 122.7 | 8.20 | 8.17 | 4.20 | 32.6 | 2-8 |
| 67 | P | 63.3 | 63.5 | 177.6 | 177.6 | - | - | - | - | 4.39 | 32.0 | 2-8 |
| 68 | G | 44.9 | 45.4 | 174.8 | 174.8 | 110.5 | 110.5 | 8.70 | 8.72 | 3.77 | - | 2-8 |
| 69 | G | 44.9 | 45.3 | 174.2 | 174.3 | 108.9 | 108.9 | 8.38 | 8.40 | 3.80 | - | 2-8 |
| 70 | V | 62.4 | 62.6 | 176.5 | 176.5 | 119.7 | 119.6 | 8.12 | 8.13 | 4.09 | 32.6 | 2-8 |
| 71 | A | 52.6 | 53.1 | 178.0 | 178.1 | 127.4 | 127.3 | 8.53 | 8.53 | 4.16 | 19.0 | 2-8 |
| 72 | D | 54.2 | 54.5 | 176.8 | 176.8 | 120.0 | 119.9 | 8.28 | 8.30 | 4.46 | 41.1 | 2-8 |
| 73 | A | 53.5 | 54.0 | 178.8 | 178.8 | 125.5 | 125.4 | 8.39 | 8.39 | 4.06 | 18.7 | 2-8 |
| 74 | A | 53.3 | 53.6 | 179.0 | 179.0 | 121.9 | 121.9 | 8.26 | 8.30 | 4.20 | 18.5 | 2-8 |
| 75 | A | 53.4 | 53.6 | 179.1 | 179.1 | 122.3 | 122.3 | 8.04 | 8.07 | 4.08 | 18.6 | 2-8 |
| 76 | A | 53.3 | 53.6 | 178.9 | 178.8 | 122.1 | 122.2 | 8.12 | 8.15 | 3.98 | 18.6 | 2-8 |
| 77 | Y | 58.9 | 59.4 | 176.6 | 176.6 | 119.8 | 119.8 | 8.07 | 8.09 | 4.30 | 38.3 | 2-8 |
| 78 | K | 56.9 | 57.5 | 177.1 | 177.2 | 122.0 | 122.0 | 7.97 | 8.01 | 3.85 | 32.9 | 2-8 |
| 79 | A | 52.8 | 53.2 | 178.4 | 178.4 | 123.5 | 123.6 | 8.07 | 8.10 | 4.00 | 18.7 | 2-8 |
| 80 | A | 52.8 | 53.0 | 178.6 | 178.6 | 122.5 | 122.6 | 8.09 | 8.11 | 4.02 | 18.9 | 2-8 |
| 81 | K | 56.2 | 56.7 | 176.9 | 177.0 | 120.0 | 119.9 | 8.11 | 8.13 | 3.98 | 32.6 | 2-8 |
| 82 | A | 52.7 | 53.0 | 178.5 | 178.5 | 124.2 | 124.2 | 8.16 | 8.18 | 4.27 | 19.0 | 2-8 |

| Residue | | $^{13}C_\alpha$ | | $^{13}CO$ | | $^{15}N$ | | $^{1}HN$ | | $H_\alpha$ | $C_\beta$ | Source |
|---|---|---|---|---|---|---|---|---|---|---|---|---|
| # | AA | hTE | Frag. | hTE | Frag. | hTE | Frag. | hTE | Frag. | Frag. | Frag. | Frag. |
| 83 | G | 44.9 | 45.1 | 174.1 | 174.1 | 108.0 | 108.0 | 8.31 | 8.33 | 3.75 | - | 2-8 |
| 84 | A | - | 52.7 | 178.3 | 178.3 | 123.6 | 123.7 | 8.16 | 8.16 | 4.32 | 19.4 | 2-8 |
| 85 | G | - | 45.2 | 174.4 | 174.5 | - | 108.1 | - | 8.47 | 3.95 | - | 2-8 |
| 86 | L | - | 55.4 | 178.2 | 178.2 | 121.6 | 121.6 | 8.22 | 8.24 | 4.26 | 42.3 | 2-8 |
| 87 | G | - | 45.3 | 173.8 | 174.6 | 109.6 | 109.7 | 8.58 | 8.60 | 3.85 | - | 2-8 |
| 88 | G | - | 45.1 | - | 173.7 | - | 108.6 | - | 8.28 | 3.75 | - | 2-8 |
| 89 | V | - | 59.9 | - | 174.6 | - | 121.3 | - | 8.13 | 4.23 | 32.6 | 2-8 |
| 90 | P | - | 63.5 | - | 177.5 | - | 140.2 | - | - | 4.39 | 32.1 | 2-8 |
| 91 | G | - | 45.3 | - | 174.3 | - | 109.6 | - | 8.52 | 3.88 | - | 2-8 |
| 92 | V | - | 62.5 | - | 177.0 | - | 119.4 | - | 8.12 | 4.05 | 32.5 | 2-8 |
| 93 | G | - | 45.5 | 174.6 | 174.7 | 112.9 | 112.8 | - | 8.69 | 3.98 | - | 2-8 |
| 94 | G | 44.8 | 45.4 | 174.2 | 174.3 | 108.8 | 108.8 | 8.33 | 8.34 | 3.87 | - | 2-8 |
| 95 | L | 54.9 | 55.4 | 178.1 | 178.1 | 121.5 | 121.5 | 8.27 | 8.29 | 4.24 | 42.4 | 2-8 |
| 96 | G | - | 45.4 | 174.0 | 174.2 | 110.0 | 109.9 | 8.56 | 8.57 | 3.95 | - | 2-8 |
| 97 | V | 61.9 | 62.3 | 176.5 | 176.5 | 119.3 | 119.2 | 8.05 | 8.06 | 4.18 | 32.8 | 2-8 |
| 98 | S | 58.0 | 58.3 | 174.4 | 174.4 | 119.8 | 119.8 | 8.52 | 8.53 | 4.32 | 63.8 | 2-8 |
| 99 | A | 52.5 | 52.9 | 178.2 | 178.2 | 126.6 | 126.6 | 8.48 | 8.49 | 4.17 | 19.1 | 2-8 |
| 100 | G | 44.7 | 45.1 | 173.6 | 173.7 | 108.2 | 108.2 | 8.39 | 8.41 | 3.83 | - | 2-8 |
| 101 | A | 51.9 | 52.3 | 177.6 | 177.6 | 123.8 | 123.8 | 8.10 | 8.11 | 4.23 | 19.5 | 2-8 |
| 102 | V | 61.9 | 62.3 | 176.2 | 176.1 | 120.7 | 120.5 | 8.25 | 8.26 | 4.11 | 32.6 | 2-8 |
| 103 | V | 59.6 | 59.8 | 174.3 | 174.3 | 127.2 | 127.1 | 8.42 | 8.43 | 4.23 | 32.6 | 2-8 |
| 104 | P | 62.6 | 62.9 | 176.5 | 176.5 | - | - | - | - | 4.40 | 32.1 | 2-8 |
| 105 | Q | 53.0 | 53.4 | 174.3 | 174.3 | 121.9 | 121.9 | 8.56 | 8.57 | 4.40 | 28.9 | 2-8 |
| 106 | P | - | 63.6 | - | 177.6 | - | 137.7 | - | - | 4.40 | 31.9 | 2-8 |
| 107 | G | - | 45.4 | - | 174.1 | - | 110.2 | - | 8.65 | 3.85 | - | 2-8 |
| 108 | A | - | 52.7 | 178.3 | 178.4 | 123.7 | 123.7 | - | 8.23 | 4.22 | 19.4 | 2-8 |
| 109 | G | - | 45.2 | 173.9 | 173.9 | 108.5 | 108.5 | 8.52 | 8.53 | 3.73 | - | 8-14 |
| 110 | V | 61.8 | 62.1 | 176.1 | 176.1 | 119.9 | 119.7 | 8.00 | 8.01 | 3.99 | 32.7 | 8-14 |
| 111 | K | 53.7 | 54.2 | 174.4 | 174.4 | 127.6 | 127.5 | 8.58 | 8.58 | 4.52 | 32.5 | 8-14 |
| 112 | P | - | 63.4 | - | 177.4 | - | 137.0 | - | - | 4.43 | 32.2 | 8-14 |
| 113 | G | 44.5 | 45.0 | 173.8 | 173.9 | 109.7 | 109.6 | - | 8.54 | 3.85 | - | 8-14 |
| 114 | K | 55.7 | 56.1 | 176.5 | 176.5 | 121.3 | 121.2 | 8.27 | 8.27 | 4.34 | 33.2 | 8-14 |
| 115 | V | 59.6 | 59.9 | 174.5 | 174.5 | 124.0 | 123.9 | 8.38 | 8.38 | 4.37 | 32.8 | 8-14 |
| 116 | P | - | 63.4 | - | 177.5 | - | 139.6 | - | - | 4.44 | 32.2 | 8-14 |
| 117 | G | - | 45.3 | - | 174.0 | - | 109.4 | - | 8.51 | 3.93 | - | 8-14 |
| 118 | V | - | 62.4 | - | 176.6 | - | 119.1 | - | 8.04 | 4.05 | 32.7 | 8-14 |
| 119 | G | 44.5 | 45.0 | 173.6 | 173.6 | 112.9 | 112.7 | 8.57 | 8.58 | 3.83 | - | 8-14 |
| 120 | L | 52.5 | 53.0 | 175.3 | 175.3 | 123.1 | 123.0 | 8.21 | 8.21 | 4.59 | 41.7 | 8-14 |
| 121 | P | - | 63.4 | 177.5 | 177.5 | - | 136.0 | - | - | 4.42 | 32.1 | 8-14 |
| 122 | G | 44.8 | 45.2 | 173.5 | 173.5 | 109.0 | 109.0 | 8.48 | 8.49 | 3.89 | - | 8-14 |
| 123 | V | 61.8 | 62.1 | 175.5 | 175.4 | 119.4 | 119.3 | 7.78 | 7.78 | 3.96 | 32.9 | 8-14 |

| Residue | | $^{13}C_{\alpha}$ | | $^{13}CO$ | | $^{15}N$ | | $^{1}HN$ | | $H_{\alpha}$ | $C_{\beta}$ | Source |
|---|---|---|---|---|---|---|---|---|---|---|---|---|
| # | AA | hTE | Frag. | hTE | Frag. | hTE | Frag. | hTE | Frag. | Frag. | Frag. | Frag. |
| 124 | Y | - | 55.6 | 174.2 | 174.2 | 125.7 | 125.4 | 8.49 | 8.49 | 4.66 | 38.4 | 8-14 |
| 125 | P | - | 63.4 | - | 177.5 | - | - | - | - | 4.45 | 32.2 | 8-14 |
| 126 | G | - | 45.2 | - | 174.6 | - | 109.4 | - | 8.55 | 3.75 | - | 8-14 |
| 127 | G | 44.8 | 45.3 | 173.7 | 173.7 | - | 108.6 | - | 8.34 | 3.85 | - | 8-14 |
| 128 | V | 61.5 | 61.9 | 175.7 | 175.7 | 119.2 | 119.0 | 7.88 | 7.89 | 4.01 | 33.0 | 8-14 |
| 129 | L | 52.6 | 53.0 | 175.2 | 175.2 | 128.0 | 127.8 | 8.44 | 8.44 | 4.26 | 41.5 | 8-14 |
| 130 | P | - | 63.4 | - | 177.4 | - | 135.9 | - | - | 4.31 | 32.2 | 8-14 |
| 131 | G | - | 45.2 | - | 173.9 | - | 109.3 | - | 8.53 | 3.82 | - | 8-14 |
| 132 | A | 52.0 | 52.4 | 177.5 | 177.5 | 123.7 | 123.5 | 8.08 | 8.09 | 4.06 | 19.5 | 8-14 |
| 133 | R | 55.4 | 55.9 | 175.6 | 175.6 | 120.0 | 119.8 | 8.25 | 8.24 | 4.12 | 30.9 | 8-14 |
| 134 | F | 55.1 | 55.4 | 174.0 | 174.0 | 122.1 | 121.9 | 8.33 | 8.33 | 4.70 | 39.1 | 8-14 |
| 135 | P | - | 63.5 | 177.3 | 177.3 | - | - | - | - | 4.42 | 32.0 | 8-14 |
| 136 | G | - | 45.2 | 173.9 | 174.0 | 109.0 | 109.0 | 8.13 | 8.14 | 3.83 | - | 8-14 |
| 137 | V | - | 62.5 | - | 176.7 | 119.3 | 119.3 | - | 8.07 | 4.04 | 32.7 | 8-14 |
| 138 | G | 44.7 | 45.2 | - | 173.7 | 113.1 | 112.9 | - | 8.63 | 3.84 | - | 8-14 |
| 139 | V | 61.7 | 62.1 | 176.1 | 176.1 | 119.8 | 119.4 | 8.01 | 8.00 | 4.10 | 32.6 | 8-14 |
| 140 | L | 52.4 | 52.9 | 175.1 | 175.1 | 128.1 | 127.8 | 8.49 | 8.49 | 4.41 | 41.7 | 8-14 |
| 141 | P | - | 63.5 | - | 177.5 | - | 135.9 | - | - | 4.38 | 32.1 | 8-14 |
| 142 | G | - | 45.1 | - | 173.8 | - | 109.4 | - | 8.49 | 3.82 | - | 8-14 |
| 143 | V | 59.6 | 59.9 | 174.7 | 174.6 | 121.2 | 121.1 | 8.03 | 8.01 | 4.34 | 32.6 | 8-14 |
| 144 | P | 62.9 | 63.2 | 177.1 | 177.1 | - | 139.8 | - | - | 4.55 | 32.0 | 8-14 |
| 145 | T | 61.9 | 62.1 | 175.3 | 175.3 | 114.7 | 114.6 | 8.38 | 8.39 | 4.28 | 70.0 | 8-14 |
| 146 | G | 44.8 | 45.4 | 173.9 | 174.0 | 111.5 | 111.4 | 8.53 | 8.53 | 3.87 | - | 8-14 |
| 147 | A | - | 52.7 | 178.3 | 178.3 | 124.0 | 123.9 | 8.34 | 8.34 | 4.11 | 19.4 | 8-14 |
| 148 | G | - | 45.1 | 173.8 | 173.9 | 108.3 | 108.3 | 8.51 | 8.52 | 3.71 | - | 8-14 |
| 149 | V | 61.9 | 62.1 | 176.1 | 176.1 | 119.9 | 119.6 | 7.98 | 7.99 | 3.99 | 32.8 | 8-14 |
| 150 | K | 53.7 | 54.2 | 174.4 | 174.4 | 127.7 | 127.5 | 8.54 | 8.54 | 4.38 | 32.6 | 8-14 |
| 151 | P | 62.7 | 62.9 | 176.7 | 176.7 | - | 137.1 | - | - | 4.44 | 32.7 | 8-14 |
| 152 | K | 55.6 | 56.1 | 176.1 | 176.1 | 122.4 | 122.3 | 8.50 | 8.50 | 4.23 | 33.2 | 8-14 |
| 153 | A | 50.1 | 50.5 | 175.4 | 175.4 | 127.3 | 127.1 | 8.46 | 8.45 | 4.56 | 18.2 | 8-14 |
| 154 | P | - | 63.4 | - | 177.5 | - | 135.3 | - | - | 4.44 | 32.1 | 8-14 |
| 155 | G | - | 45.2 | - | 174.2 | 109.3 | 109.1 | 8.46 | 8.48 | 3.72 | - | 8-14 |
| 156 | V | - | 62.5 | - | 176.9 | - | 119.2 | - | 8.10 | 4.05 | 32.6 | 8-14 |
| 157 | G | - | 45.3 | - | 174.7 | - | 112.7 | - | 8.68 | 3.77 | - | 8-14 |
| 158 | G | - | 45.2 | 173.9 | 174.0 | - | 108.8 | - | 8.32 | 3.84 | - | 8-14 |
| 159 | A | - | 52.7 | 177.6 | 177.6 | 123.8 | 123.6 | 8.25 | 8.26 | 4.13 | 19.2 | 8-14 |
| 160 | F | 57.3 | 57.7 | 175.5 | 175.5 | 119.5 | 119.3 | 8.32 | 8.33 | 4.54 | 39.5 | 8-14 |
| 161 | A | 52.1 | 52.5 | 177.6 | 177.6 | 126.3 | 126.0 | 8.19 | 8.20 | 4.25 | 19.4 | 8-14 |
| 162 | G | 44.5 | 45.1 | 173.4 | 173.4 | 107.5 | 107.4 | 7.71 | 7.75 | 3.82 | - | 8-14 |
| 163 | I | - | 58.6 | 174.7 | 174.6 | 122.1 | 121.9 | 8.12 | 8.12 | 4.43 | 38.6 | 8-14 |
| 164 | P | - | 63.6 | - | - | - | 140.3 | - | - | 4.42 | - | 8-14 |

| Residue | | $^{13}C_{\alpha}$ | | $^{13}CO$ | | $^{15}N$ | | $^{1}HN$ | | $H_{\alpha}$ | $C_{\beta}$ | Source |
|---|---|---|---|---|---|---|---|---|---|---|---|---|
| **#** | **AA** | **hTE** | **Frag.** | **hTE** | **Frag.** | **hTE** | **Frag.** | **hTE** | **Frag.** | **Frag.** | **Frag.** | **Frag.** |
| 165 | G | 44.9 | 45.3 | 173.9 | 174.0 | - | - | - | - | 4.01 | - | 8-14 |
| 166 | V | 61.7 | 62.0 | 176.4 | 176.4 | 118.8 | 118.7 | 8.01 | 8.00 | 4.15 | 33.3 | 8-14 |
| 167 | G | 44.1 | 44.6 | 172.2 | 172.3 | 112.6 | 112.5 | 8.46 | 8.48 | 4.00 | - | 8-14 |
| 168 | P | 62.9 | 63.5 | 177.1 | 177.1 | - | 133.9 | - | - | 4.35 | - | 8-14 |
| 169 | F | 57.5 | 57.7 | 176.4 | 176.4 | 120.1 | 119.8 | 8.47 | 8.48 | 4.44 | 39.0 | 8-14 |
| 170 | G | 44.8 | 45.2 | 174.1 | 174.2 | 110.6 | 110.4 | 8.23 | 8.23 | 3.79 | - | 8-14 |
| 171 | G | 43.9 | 44.5 | 171.3 | 171.3 | 108.6 | 108.6 | 7.88 | 7.92 | 3.86 | - | 8-14 |
| 172 | P | 62.7 | 63.4 | 177.0 | 176.9 | - | 134.2 | - | - | 4.44 | 32.2 | 8-14 |
| 173 | Q | 52.9 | 53.4 | 174.0 | 173.9 | 122.0 | 121.9 | 8.61 | 8.60 | 4.38 | 29.0 | 8-14 |
| 174 | P | - | 63.5 | - | 177.5 | - | 137.3 | - | - | 4.40 | 32.2 | 8-14 |
| 175 | G | 44.8 | 45.1 | 173.7 | 173.8 | - | 109.6 | - | 8.58 | 3.82 | - | 8-14 |
| 176 | V | 59.4 | 59.7 | 174.4 | 174.4 | 120.9 | 120.7 | 7.91 | 7.92 | 4.20 | 32.6 | 8-14 |
| 177 | P | 62.7 | 63.0 | 176.7 | 176.8 | - | - | - | - | 4.38 | 32.2 | 8-14 |
| 178 | L | 55.1 | 55.6 | 178.0 | 178.0 | 122.5 | 122.5 | 8.42 | 8.42 | 4.25 | 42.3 | 8-14 |
| 179 | G | 44.5 | 45.0 | 173.2 | 173.3 | 109.4 | 109.3 | - | 8.42 | 3.80 | - | 8-14 |
| 180 | Y | 55.6 | 55.9 | 173.9 | 173.9 | 121.2 | 121.0 | 7.95 | 7.95 | 4.85 | 40.2 | 8-14 |
| 181 | P | 62.6 | 63.0 | 176.5 | 176.5 | - | 137.6 | - | - | 4.44 | - | 8-14 |
| 182 | I | 60.8 | 61.1 | 176.3 | 176.3 | 121.6 | 121.4 | 8.25 | 8.23 | 4.06 | - | 8-14 |
| 183 | K | 55.4 | 55.9 | 175.6 | 175.6 | 125.9 | 125.6 | 8.43 | 8.44 | 4.21 | - | 8-14 |
| 184 | A | 50.1 | 50.5 | 175.3 | 175.3 | 127.5 | 127.2 | 8.41 | 8.39 | 4.50 | 18.3 | 8-14 |
| 185 | P | 62.5 | 62.8 | 176.5 | 176.5 | - | 135.5 | - | - | 4.41 | 32.1 | 8-14 |
| 186 | K | 55.7 | 56.1 | 176.3 | 176.4 | 121.9 | 121.8 | 8.46 | 8.45 | 4.18 | 33.1 | 8-14 |
| 187 | L | - | 52.9 | 175.3 | 175.3 | 125.4 | 124.7 | 8.39 | 8.35 | 4.30 | 41.8 | 8-14 |
| 188 | P | - | 63.8 | 177.6 | 177.6 | - | - | - | - | 4.41 | 31.8 | 8-14 |
| 189 | G | 45.0 | 45.4 | 174.6 | 174.7 | 110.0 | 109.9 | 8.36 | 8.36 | 3.75 | - | 8-14 |
| 190 | G | - | 45.3 | - | 173.9 | 108.8 | 108.6 | - | 8.27 | 3.68 | - | 8-14 |
| 191 | Y | 58.1 | 58.3 | 176.3 | 176.3 | 120.0 | 119.7 | - | 8.14 | 4.44 | 38.8 | 8-14 |
| 192 | G | 44.5 | 45.1 | 173.5 | 173.5 | 110.6 | 110.5 | 8.41 | 8.43 | 3.75 | - | 8-14 |
| 193 | L | - | 53.1 | 175.5 | 175.5 | 122.6 | 122.4 | 8.03 | 8.03 | 4.36 | 41.6 | 8-14 |
| 194 | P | - | 62.2 | - | - | - | - | - | - | - | 32.8 | 8-14 |
| 195 | Y | 57.7 | 57.9 | 175.9 | 176.0 | - | 119.6 | - | 7.96 | 4.66 | 38.8 | 8-14 |
| 196 | T | 61.2 | 61.5 | 174.2 | 174.3 | 116.0 | 115.6 | 8.08 | 8.09 | 4.18 | - | 8-14 |
| 197 | T | 61.9 | 62.1 | 175.0 | 175.1 | 115.7 | 115.4 | 8.15 | 8.16 | 4.20 | 70.2 | 8-14 |
| 198 | G | 44.9 | 45.3 | 173.7 | 173.7 | 111.4 | 111.2 | 8.43 | 8.43 | 3.85 | - | 8-14 |
| 199 | K | 55.6 | 56.0 | 176.4 | 176.3 | 120.8 | 120.7 | 8.19 | 8.19 | 4.10 | 33.1 | 8-14 |
| 200 | L | 52.5 | 52.8 | 175.3 | 175.3 | 124.8 | 125.1 | 8.34 | 8.37 | 4.41 | 41.7 | 8-14 |
| 201 | P | 62.6 | 63.0 | 176.3 | 176.2 | - | - | - | - | 4.42 | 31.7 | 8-14 |
| 202 | Y | 58.1 | 58.5 | 176.4 | 176.3 | 120.5 | 120.3 | 8.25 | 8.24 | 4.31 | 38.6 | 8-14 |
| 203 | G | 44.7 | 45.1 | 173.5 | 173.6 | 111.4 | 111.2 | 8.22 | 8.24 | 3.42 | - | 8-14 |
| 204 | Y | 57.7 | 58.1 | 176.0 | 175.9 | 119.7 | 119.5 | 7.93 | 7.95 | 4.46 | 39.3 | 8-14 |
| 205 | G | - | 44.6 | 171.9 | 171.9 | 110.3 | 110.2 | 8.28 | 8.30 | 3.96 | - | 8-14 |

| Residue | | $^{13}C_{\alpha}$ | | $^{13}CO$ | | $^{15}N$ | | $^{1}HN$ | | $H_{\alpha}$ | $C_{\beta}$ | Source |
|---|---|---|---|---|---|---|---|---|---|---|---|---|
| # | AA | hTE | Frag. | hTE | Frag. | hTE | Frag. | hTE | Frag. | Frag. | Frag. | Frag. |
| 206 | P | 63.5 | 63.8 | 177.9 | 177.9 | - | 134.5 | - | - | 4.41 | 32.0 | 8-14 |
| 207 | G | 44.9 | 45.3 | 174.8 | 174.8 | 109.7 | 109.7 | 8.69 | 8.70 | 3.73 | - | 8-14 |
| 208 | G | 45.0 | 45.0 | 174.1 | 174.1 | 108.6 | 108.6 | 8.20 | 8.22 | 3.83 | - | 8-14 |
| 209 | V | 62.0 | 62.2 | 176.2 | 176.1 | 119.6 | 119.3 | 8.00 | 8.02 | 3.81 | 32.6 | 8-14 |
| 210 | A | 52.5 | 52.9 | 178.4 | 178.4 | 127.5 | 127.4 | 8.49 | 8.50 | 4.15 | 19.0 | 8-14 |
| 211 | G | 44.7 | 45.3 | 174.0 | 174.1 | 108.4 | 108.4 | 8.29 | 8.30 | 3.81 | - | 8-14 |
| 212 | A | 52.4 | 52.7 | 177.7 | 177.8 | 123.8 | 123.7 | 8.13 | 8.14 | 4.07 | 19.4 | 8-14 |
| 213 | A | - | 52.8 | 178.4 | 178.4 | 122.9 | 122.8 | 8.35 | 8.36 | 4.22 | 19.1 | 8-14 |
| 214 | G | - | 45.3 | 174.2 | 174.3 | 107.9 | 107.9 | 8.29 | 8.29 | 3.91 | - | 13-19 |
| 215 | K | 55.8 | 56.2 | 176.5 | 176.5 | 121.0 | 121.0 | 8.15 | 8.15 | 4.26 | 33.0 | 13-19 |
| 216 | A | 52.4 | 52.7 | 178.0 | 178.0 | 124.9 | 124.9 | 8.38 | 8.38 | 4.24 | 19.1 | 13-19 |
| 217 | G | 44.4 | 45.2 | 173.4 | 173.4 | 107.9 | 107.9 | 8.30 | 8.30 | 3.85 | - | 13-19 |
| 218 | Y | 55.6 | 55.9 | 174.1 | 174.1 | 120.9 | 120.9 | 8.02 | 8.02 | 4.75 | 38.1 | 13-19 |
| 219 | P | 62.9 | 63.2 | 177.0 | 177.1 | - | 137.5 | - | - | 4.50 | 31.9 | 13-19 |
| 220 | T | 61.9 | 62.1 | 175.3 | 175.3 | 114.6 | 114.5 | 8.37 | 8.38 | 4.35 | 70.0 | 13-19 |
| 221 | G | 44.9 | 45.3 | 174.5 | 174.5 | 111.3 | 111.3 | 8.56 | 8.56 | 4.02 | - | 13-19 |
| 222 | T | 61.9 | 62.1 | 175.3 | 175.3 | 113.2 | 113.1 | 8.25 | 8.25 | 4.33 | 69.9 | 13-19 |
| 223 | G | 44.8 | 45.3 | 173.8 | 173.8 | 111.4 | 111.5 | 8.57 | 8.57 | 3.94 | - | 13-19 |
| 224 | V | 61.8 | 62.2 | 176.6 | 176.5 | 118.3 | 118.3 | 8.02 | 8.01 | 4.17 | 32.9 | 13-19 |
| 225 | G | 44.0 | 44.9 | 172.6 | 172.6 | 112.6 | 112.7 | 8.55 | 8.56 | 4.14 | - | 13-19 |
| 226 | P | 63.9 | 64.3 | 178.5 | 178.6 | - | 134.5 | - | - | 4.35 | 32.0 | 13-19 |
| 227 | Q | 57.0 | 57.5 | 177.4 | 177.5 | 119.9 | 119.9 | 8.57 | 8.56 | 4.22 | 28.5 | 13-19 |
| 228 | A | 53.6 | 54.1 | 179.6 | 179.7 | 124.9 | 124.9 | 8.26 | 8.26 | 4.20 | 18.3 | 13-19 |
| 229 | A | 53.8 | 54.3 | 179.6 | 179.8 | 122.9 | 122.9 | 8.39 | 8.41 | 4.21 | 18.1 | 13-19 |
| 230 | A | 53.9 | 54.3 | 179.8 | 179.9 | 122.7 | 122.7 | 8.18 | 8.17 | 4.22 | 18.0 | 13-19 |
| 231 | A | 53.8 | 54.4 | 179.6 | 179.7 | 122.7 | 122.7 | 8.15 | 8.14 | 4.22 | 17.9 | 13-19 |
| 232 | A | 53.9 | 54.6 | 179.8 | 179.9 | 122.1 | 122.0 | 8.12 | 8.11 | 4.19 | 18.0 | 13-19 |
| 233 | A | 53.5 | 54.2 | 179.5 | 179.6 | 122.5 | 122.4 | 8.14 | 8.15 | 4.21 | 17.8 | 13-19 |
| 234 | A | 53.6 | 54.2 | 179.6 | 179.8 | 122.3 | 122.2 | 8.07 | 8.06 | 4.18 | 18.0 | 13-19 |
| 235 | K | 57.7 | 58.2 | 178.0 | 178.1 | 120.0 | 120.0 | 8.00 | 7.99 | 4.11 | 32.7 | 13-19 |
| 236 | A | 53.3 | 53.9 | 178.8 | 178.9 | 122.8 | 122.7 | 8.00 | 7.99 | 4.18 | 18.2 | 13-19 |
| 237 | A | 52.9 | 53.6 | 178.6 | 178.7 | 121.4 | 121.3 | 8.04 | 8.03 | 4.20 | 18.4 | 13-19 |
| 238 | A | 52.6 | 53.8 | 178.3 | 178.3 | 121.7 | 121.6 | 7.88 | 7.88 | 4.20 | 18.3 | 13-19 |
| 239 | K | 56.5 | 57.2 | 176.8 | 176.9 | 119.3 | 119.1 | 7.91 | 7.88 | 4.16 | 32.8 | 13-19 |
| 240 | F | 57.8 | 58.1 | 176.5 | 176.6 | 119.9 | 119.7 | 8.17 | 8.16 | 4.55 | 39.3 | 13-19 |
| 241 | G | 44.8 | 45.3 | 174.0 | 174.0 | 110.2 | 110.2 | 8.27 | 8.28 | 3.90 | - | 13-19 |
| 242 | A | - | 52.9 | 178.5 | 178.5 | 123.8 | 123.8 | 8.24 | 8.24 | 4.29 | 19.2 | 13-19 |
| 243 | G | 44.9 | 45.3 | 174.1 | 174.1 | 108.3 | 108.2 | 8.45 | 8.46 | 3.90 | - | 13-19 |
| 244 | A | 52.2 | 52.6 | 177.7 | 177.7 | 124.0 | 123.9 | 8.15 | 8.15 | 4.26 | 19.2 | 13-19 |
| 245 | A | 52.5 | 52.8 | 178.3 | 178.3 | 122.8 | 122.8 | 8.34 | 8.34 | 4.24 | 19.0 | 13-19 |
| 246 | G | 44.9 | 45.2 | 173.7 | 173.7 | 107.9 | 107.9 | 8.30 | 8.30 | 3.88 | - | 13-19 |

| Residue | | $^{13}C_\alpha$ | | $^{13}CO$ | | $^{15}N$ | | $^{1}HN$ | | $H_\alpha$ | $C_\beta$ | Source |
|---|---|---|---|---|---|---|---|---|---|---|---|---|
| **#** | **AA** | **hTE** | **Frag.** | **hTE** | **Frag.** | **hTE** | **Frag.** | **hTE** | **Frag.** | **Frag.** | **Frag.** | **Frag.** |
| 247 | V | 61.8 | 62.1 | 176.1 | 176.1 | 119.5 | 119.4 | 7.94 | 7.94 | 4.09 | 32.6 | 13-19 |
| 248 | L | 52.6 | 52.9 | 175.1 | 175.1 | 128.2 | 128.2 | 8.47 | 8.47 | 4.60 | 41.5 | 13-19 |
| 249 | P | - | 63.5 | - | 177.5 | - | - | - | - | 4.36 | 32.2 | 13-19 |
| 250 | G | - | 45.2 | - | 174.3 | - | 109.4 | - | 8.49 | 3.94 | - | 13-19 |
| 251 | V | - | 62.6 | 177.0 | 177.0 | - | 119.5 | - | 8.07 | 4.14 | 32.6 | 13-19 |
| 252 | G | - | 45.3 | 174.7 | 174.7 | 112.6 | 112.8 | 8.66 | 8.67 | 3.96 | - | 13-19 |
| 253 | G | 44.8 | 45.2 | 173.9 | 173.9 | 109.1 | 109.0 | 8.32 | 8.33 | 3.94 | - | 13-19 |
| 254 | A | 52.1 | 52.6 | 178.2 | 178.2 | 123.6 | 123.6 | 8.32 | 8.33 | 4.31 | 19.1 | 13-19 |
| 255 | G | - | 45.0 | 173.7 | 173.7 | 108.4 | 108.4 | 8.48 | 8.48 | 3.92 | - | 13-19 |
| 256 | V | - | 60.1 | - | 174.6 | 121.4 | 121.3 | - | 8.09 | 4.40 | 32.4 | 13-19 |
| 257 | P | - | 63.5 | - | 177.4 | - | 140.0 | - | - | 4.35 | 32.1 | 13-19 |
| 258 | G | - | 44.9 | - | 173.7 | - | 109.5 | - | 8.53 | 3.91 | - | 13-19 |
| 259 | V | - | 59.8 | - | 174.6 | - | 121.3 | - | 8.10 | 4.40 | 32.4 | 13-19 |
| 260 | P | - | 63.7 | - | 177.6 | - | 140.0 | - | - | 4.36 | 32.0 | 13-19 |
| 261 | G | - | 45.2 | - | 173.7 | - | 110.1 | - | 8.60 | 3.91 | - | 13-19 |
| 262 | A | 52.2 | 52.4 | 177.6 | 177.5 | 123.7 | 123.6 | 8.10 | 8.10 | 4.28 | 19.1 | 13-19 |
| 263 | I | 58.4 | 58.6 | 174.7 | 174.7 | 122.6 | 122.6 | 8.35 | 8.35 | 4.43 | 38.6 | 13-19 |
| 264 | P | - | 63.6 | - | 177.4 | - | 141.0 | - | - | 4.36 | 32.2 | 13-19 |
| 265 | G | 44.7 | 45.2 | 174.3 | 174.3 | 109.4 | 109.5 | 8.51 | 8.50 | 3.95 | - | 13-19 |
| 266 | I | 61.0 | 61.3 | 177.0 | 177.0 | 120.2 | 120.0 | 8.09 | 8.09 | 4.19 | 38.4 | 13-19 |
| 267 | G | - | 45.3 | - | 174.6 | 112.9 | 113.0 | 8.66 | 8.67 | 3.96 | - | 13-19 |
| 268 | G | 44.8 | 45.1 | 174.1 | 174.1 | 108.8 | 108.7 | - | 8.28 | 3.95 | - | 13-19 |
| 269 | I | 60.8 | 61.0 | 176.2 | 176.2 | 120.5 | 120.4 | 8.12 | 8.12 | 4.15 | 38.7 | 13-19 |
| 270 | A | 52.3 | 52.6 | 178.1 | 178.1 | 128.3 | 128.2 | 8.53 | 8.54 | 4.29 | 19.1 | 13-19 |
| 271 | G | 44.9 | 45.2 | 174.2 | 174.1 | 108.4 | 108.4 | 8.38 | 8.39 | 3.94 | - | 13-19 |
| 272 | V | 62.3 | 62.7 | 177.0 | 176.9 | 119.4 | 119.3 | 8.13 | 8.11 | 4.10 | 32.5 | 13-19 |
| 273 | G | 44.8 | 45.3 | 174.1 | 174.1 | 112.6 | 112.6 | 8.69 | 8.70 | 3.99 | - | 13-19 |
| 274 | T | 60.0 | 60.2 | 173.8 | 173.8 | 113.9 | 113.8 | 7.97 | 7.96 | 4.65 | 69.7 | 13-19 |
| 275 | P | 64.1 | 64.7 | 178.6 | 178.6 | - | 138.0 | - | - | 4.38 | 31.9 | 13-19 |
| 276 | A | 53.9 | 54.3 | 179.8 | 179.8 | 122.3 | 122.3 | 8.42 | 8.43 | 4.20 | 18.4 | 13-19 |
| 277 | A | 53.9 | 54.8 | 180.2 | 180.2 | 123.1 | 123.1 | 8.21 | 8.21 | 4.21 | 18.0 | 13-19 |
| 278 | A | 53.9 | 54.5 | 180.2 | 180.3 | 122.9 | 122.9 | 8.21 | 8.21 | 4.22 | 18.0 | 13-19 |
| 279 | A | 53.9 | 54.5 | 180.3 | 180.4 | 122.7 | 122.7 | 8.24 | 8.25 | 4.23 | 17.8 | 13-19 |
| 280 | A | 54.1 | 54.6 | 180.3 | 180.3 | 122.6 | 122.7 | 8.15 | 8.16 | 4.22 | 17.9 | 13-19 |
| 281 | A | - | 54.6 | - | 180.3 | - | 122.5 | - | 8.14 | 4.22 | 17.9 | 13-19 |
| 282 | A | - | 54.7 | 180.2 | 180.2 | 122.2 | 122.4 | 8.08 | 8.13 | 4.21 | 17.9 | 13-19 |
| 283 | A | - | 54.6 | 180.2 | 180.2 | 121.9 | 121.9 | 8.08 | 8.08 | 4.19 | 17.9 | 13-19 |
| 284 | A | - | 54.6 | 179.8 | 179.9 | 122.2 | 122.2 | - | 8.10 | 4.20 | 17.8 | 13-19 |
| 285 | A | 53.8 | 54.0 | 179.8 | 179.8 | 121.8 | 121.8 | 8.00 | 8.01 | 4.19 | 18.4 | 13-19 |
| 286 | K | 58.0 | 58.6 | 177.9 | 177.9 | 119.5 | 119.5 | 7.87 | 7.88 | 4.13 | 32.8 | 13-19 |
| 287 | A | 53.3 | 53.8 | 179.1 | 179.1 | 122.0 | 121.9 | 7.93 | 7.94 | 4.19 | 18.2 | 13-19 |

| Residue | | $^{13}C_{\alpha}$ | | $^{13}$CO | | $^{15}$N | | $^{1}$HN | | $H_{\alpha}$ | $C_{\beta}$ | Source |
|---|---|---|---|---|---|---|---|---|---|---|---|---|
| # | AA | hTE | Frag. | hTE | Frag. | hTE | Frag. | hTE | Frag. | Frag. | Frag. | Frag. |
| 288 | A | 52.9 | 53.6 | 178.5 | 178.6 | 121.1 | 121.1 | 7.90 | 7.90 | 4.18 | 18.4 | 13-19 |
| 289 | K | 57.0 | 57.5 | 177.1 | 177.2 | 119.0 | 118.9 | 7.82 | 7.82 | 4.14 | 32.8 | 13-19 |
| 290 | Y | 58.1 | 58.6 | 176.8 | 176.8 | 119.0 | 118.8 | 8.09 | 8.09 | 4.54 | 38.7 | 13-19 |
| 291 | G | 45.1 | 45.4 | 174.1 | 174.2 | 109.8 | 109.8 | 8.21 | 8.22 | 3.91 | - | 13-19 |
| 292 | A | 52.6 | 53.0 | 178.1 | 178.1 | - | 123.9 | - | 8.19 | 4.30 | 18.6 | 13-19 |
| 293 | A | 52.4 | 52.8 | 177.8 | 177.8 | 122.7 | 122.6 | 8.26 | 8.26 | 4.24 | 18.5 | 13-19 |
| 294 | A | 52.4 | 52.9 | 178.3 | 178.3 | 122.6 | 122.5 | 8.17 | 8.16 | 4.23 | 19.0 | 13-19 |
| 295 | G | 44.8 | 45.3 | 173.8 | 173.9 | 107.4 | 107.4 | 8.25 | 8.25 | 3.88 | - | 13-19 |
| 296 | L | 54.6 | 54.6 | 177.2 | 177.2 | 121.4 | 121.3 | 8.01 | 8.01 | 4.34 | 42.3 | 13-19 |
| 297 | V | 59.4 | 59.6 | 174.4 | 174.3 | 122.7 | 122.7 | 8.21 | 8.21 | 4.40 | 32.6 | 13-19 |
| 298 | P | 63.2 | 63.4 | 177.4 | 177.4 | - | 140.3 | - | - | 4.22 | 31.9 | 13-19 |
| 299 | G | 44.7 | 45.2 | 174.3 | 174.3 | 110.1 | 110.2 | 8.42 | 8.43 | 3.89 | - | 13-19 |
| 300 | G | 44.1 | 44.7 | 171.9 | 171.9 | 108.9 | 108.9 | 8.22 | 8.22 | 4.12 | - | 13-19 |
| 301 | P | 63.5 | 63.6 | 177.7 | 177.6 | - | 134.0 | - | - | 4.37 | 32.1 | 13-19 |
| 302 | G | 44.6 | 45.3 | 173.7 | 173.7 | 109.5 | 109.5 | 8.54 | 8.56 | 3.92 | - | 13-19 |
| 303 | F | 57.4 | 57.7 | 175.8 | 175.8 | 119.7 | 119.5 | 8.11 | 8.10 | 4.62 | 40.0 | 13-19 |
| 304 | G | 44.1 | 44.4 | 171.7 | 171.7 | 110.5 | 110.5 | 8.31 | 8.32 | 4.06 | - | 13-19 |
| 305 | P | - | 63.7 | - | 177.7 | - | 134.0 | - | - | 4.36 | 31.9 | 13-19 |
| 306 | G | - | 45.3 | 173.8 | 173.8 | - | 109.7 | - | 8.60 | 3.92 | - | 13-19 |
| 307 | V | 61.9 | 62.4 | 176.2 | 176.2 | 120.1 | 120.0 | 7.98 | 7.98 | 4.08 | 32.7 | 13-19 |
| 308 | V | 62.3 | 62.4 | 176.5 | 176.5 | 124.7 | 124.7 | 8.32 | 8.33 | 4.08 | 32.6 | 13-19 |
| 309 | G | 44.6 | 44.9 | 173.5 | 173.5 | 113.0 | 113.0 | 8.49 | 8.50 | 3.89 | - | 13-19 |
| 310 | V | 59.5 | 59.9 | 174.7 | 174.6 | 121.2 | 121.2 | 8.11 | 8.11 | 4.40 | 32.4 | 13-19 |
| 311 | P | - | 63.8 | - | 177.6 | - | 140.0 | - | - | 4.37 | 31.9 | 13-19 |
| 312 | G | - | 45.3 | - | 173.9 | - | 110.3 | - | 8.64 | 3.93 | - | 13-19 |
| 313 | A | - | 52.5 | - | 178.2 | - | 123.5 | - | 8.21 | 4.30 | 19.1 | 13-19 |
| 314 | G | - | 45.0 | - | 173.7 | - | 108.3 | - | 8.46 | 3.89 | - | 13-19 |
| 315 | V | - | 59.9 | - | 174.6 | - | 121.1 | - | 8.08 | 4.37 | 32.8 | 13-19 |
| 316 | P | - | 63.6 | - | 177.5 | - | 140.0 | - | - | 4.36 | 32.1 | 13-19 |
| 317 | G | - | 45.2 | - | 174.0 | - | 109.7 | - | 8.55 | 3.93 | - | 13-19 |
| 318 | V | - | 62.4 | - | 176.7 | - | 119.3 | - | 8.05 | 4.12 | 32.6 | 13-19 |
| 319 | G | - | 44.9 | - | 173.6 | - | 112.9 | - | 8.60 | 3.92 | - | 13-19 |
| 320 | V | - | 59.8 | - | 174.6 | - | 121.2 | - | 8.10 | 4.40 | 32.4 | 13-19 |
| 321 | P | - | 63.6 | - | 177.6 | - | 140.0 | - | - | 4.36 | 31.9 | 13-19 |
| 322 | G | - | 45.1 | - | 173.9 | - | 110.4 | - | 8.62 | 3.91 | - | 13-19 |
| 323 | A | - | 52.6 | - | 178.2 | - | 123.5 | - | 8.19 | 4.29 | 19.1 | 13-19 |
| 324 | G | 44.6 | 44.9 | 173.6 | 173.7 | 108.3 | 108.3 | - | 8.49 | 3.90 | - | 13-19 |
| 325 | I | 58.3 | 58.6 | 174.7 | 174.7 | 122.1 | 122.0 | 8.03 | 8.03 | 4.42 | - | 13-19 |
| 326 | P | 62.9 | 63.2 | 176.6 | 176.6 | - | 140.4 | - | - | 4.40 | 32.2 | 13-19 |
| 327 | V | 62.0 | 62.4 | 176.2 | 176.2 | 121.9 | 121.9 | 8.39 | 8.40 | 4.03 | 32.8 | 18-26 |
| 328 | V | 59.4 | 59.8 | 174.4 | 174.3 | 127.1 | 127.1 | 8.41 | 8.42 | 4.41 | 32.5 | 18-26 |

| Residue | | $^{13}C_{\alpha}$ | | $^{13}CO$ | | $^{15}N$ | | $^{1}HN$ | | $H_{\alpha}$ | $C_{\beta}$ | Source |
|---|---|---|---|---|---|---|---|---|---|---|---|---|
| # | AA | hTE | Frag. | hTE | Frag. | hTE | Frag. | hTE | Frag. | Frag. | Frag. | Frag. |
| 329 | P | - | 63.7 | - | 177.6 | - | 140.1 | - | - | 4.34 | 32.0 | 18-26 |
| 330 | G | - | 45.3 | - | 173.9 | - | 110.3 | - | 8.60 | 3.93 | - | 18-26 |
| 331 | A | - | 52.5 | - | 178.2 | - | 123.5 | - | 8.21 | 4.30 | 19.1 | 18-26 |
| 332 | G | - | 45.1 | 173.6 | 173.6 | 108.3 | 108.3 | 8.47 | 8.48 | 3.90 | - | 18-26 |
| 333 | I | 58.3 | 58.7 | 174.9 | 174.8 | 122.1 | 122.1 | 8.12 | 8.12 | - | 38.4 | 18-26 |
| 334 | P | 63.4 | 63.8 | 177.6 | 177.6 | - | - | - | - | 4.36 | 32.0 | 18-26 |
| 335 | G | 44.8 | 45.3 | 173.8 | 173.8 | 110.2 | 110.4 | 8.60 | 8.63 | 3.92 | - | 18-26 |
| 336 | A | 51.9 | 52.4 | 177.4 | 177.4 | 123.6 | 123.6 | 8.07 | 8.07 | 4.28 | 19.4 | 18-26 |
| 337 | A | 51.9 | 52.3 | 177.5 | 177.5 | 123.9 | 123.8 | 8.39 | 8.40 | 4.30 | 19.2 | 18-26 |
| 338 | V | 59.6 | 59.9 | 174.6 | 174.6 | 122.0 | 122.0 | 8.30 | 8.31 | 4.38 | 32.5 | 18-26 |
| 339 | P | - | 63.5 | - | 177.5 | - | - | - | - | 4.36 | 32.1 | 18-26 |
| 340 | G | 44.7 | 45.2 | 173.8 | 173.9 | 109.7 | 109.7 | - | 8.54 | 3.94 | - | 18-26 |
| 341 | V | 62.0 | 62.4 | 176.2 | 176.2 | 120.4 | 120.4 | 8.04 | 8.04 | 4.12 | 33.0 | 18-26 |
| 342 | V | 61.6 | 62.2 | 175.9 | 175.9 | 125.8 | 125.8 | 8.44 | 8.44 | 4.16 | 33.1 | 18-26 |
| 343 | S | 56.3 | 56.3 | 173.0 | 173.0 | 123.1 | 123.0 | 8.70 | 8.70 | 4.74 | 63.5 | 18-26 |
| 344 | P | 64.5 | 65.1 | 179.1 | 179.1 | - | 137.3 | - | - | 4.35 | 31.8 | 18-26 |
| 345 | E | 58.6 | 59.2 | 178.2 | 178.2 | 119.2 | 119.2 | 8.72 | 8.71 | 4.13 | 29.1 | 18-26 |
| 346 | A | 54.1 | 54.7 | 180.6 | 180.6 | 124.6 | 124.6 | 8.11 | 8.12 | 4.13 | 18.4 | 18-26 |
| 347 | A | 54.1 | 54.5 | 180.1 | 180.1 | 123.1 | 123.1 | 8.56 | 8.57 | 4.12 | 18.0 | 18-26 |
| 348 | A | 54.2 | 54.7 | 180.4 | 180.4 | 122.4 | 122.4 | 8.16 | 8.17 | 4.21 | 18.0 | 18-26 |
| 349 | K | 58.3 | 58.8 | 178.7 | 178.8 | 120.4 | 120.4 | 8.01 | 8.02 | 4.13 | 32.5 | 18-26 |
| 350 | A | 53.9 | 54.4 | 180.0 | 180.0 | 122.6 | 122.8 | 8.04 | 8.06 | 4.18 | 18.1 | 18-26 |
| 351 | A | 53.9 | 54.2 | 179.6 | 179.6 | 122.2 | 122.2 | 8.15 | 8.17 | 4.20 | 18.0 | 18-26 |
| 352 | A | 53.7 | 54.2 | 179.6 | 179.6 | 122.0 | 121.9 | 7.97 | 7.99 | 4.20 | 18.0 | 18-26 |
| 353 | K | 57.7 | 58.4 | 177.8 | 177.8 | 119.6 | 119.6 | 7.91 | 7.92 | 4.16 | 32.7 | 18-26 |
| 354 | A | 53.3 | 53.6 | 178.8 | 178.9 | 122.2 | 122.1 | 7.95 | 7.97 | 4.17 | 18.4 | 18-26 |
| 355 | A | 52.9 | 53.4 | 178.5 | 178.5 | 121.3 | 121.3 | 7.92 | 7.93 | 4.19 | 18.4 | 18-26 |
| 356 | K | 56.9 | 57.4 | 177.0 | 177.1 | 119.1 | 119.1 | 7.86 | 7.85 | 4.14 | 32.9 | 18-26 |
| 357 | Y | 58.0 | 58.4 | 176.6 | 176.6 | 119.2 | 119.1 | 8.08 | 8.08 | 4.54 | 38.7 | 18-26 |
| 358 | G | 44.8 | 45.3 | 173.5 | 173.5 | 109.7 | 109.8 | 8.15 | 8.17 | 3.91 | - | 18-26 |
| 359 | A | 51.9 | 52.3 | 177.5 | 177.5 | 123.6 | 123.4 | 8.08 | 8.08 | 4.30 | 19.2 | 18-26 |
| 360 | R | 53.4 | 54.0 | 174.1 | 174.1 | 121.7 | 121.6 | 8.37 | 8.37 | 4.60 | 30.0 | 18-26 |
| 361 | P | - | 63.4 | - | - | - | 136.9 | - | - | 4.39 | - | 18-26 |
| 362 | G | - | - | - | 174.0 | - | - | - | - | - | - | 18-26 |
| 363 | V | - | - | - | 176.7 | - | 119.1 | - | 8.05 | - | - | 18-26 |
| 364 | G | 44.6 | 45.0 | 174.3 | 174.3 | 112.9 | 113.1 | - | 8.59 | 3.96 | - | 18-26 |
| 365 | V | 62.3 | 62.7 | 177.0 | 177.1 | 119.6 | 119.5 | 8.18 | 8.19 | 4.10 | 32.3 | 18-26 |
| 366 | G | 44.9 | 45.3 | 174.6 | 174.6 | 113.1 | 112.8 | 8.71 | 8.70 | 3.95 | - | 18-26 |
| 367 | G | 44.5 | 44.9 | 173.7 | 173.6 | 108.5 | 108.4 | 8.26 | 8.26 | 3.90 | - | 18-26 |
| 368 | I | 58.4 | 58.7 | 174.7 | 174.6 | 121.8 | 121.7 | 8.00 | 7.99 | 4.41 | - | 18-26 |
| 369 | P | 62.9 | 63.4 | 176.6 | 176.6 | - | 140.4 | - | - | 4.42 | 32.2 | 18-26 |

| Residue | | $^{13}C_{\alpha}$ | | $^{13}CO$ | | $^{15}N$ | | $^{1}HN$ | | $H_{\alpha}$ | $C_{\beta}$ | Source |
|---|---|---|---|---|---|---|---|---|---|---|---|---|
| # | AA | hTE | Frag. | hTE | Frag. | hTE | Frag. | hTE | Frag. | Frag. | Frag. | Frag. |
| 370 | T | 61.5 | 61.9 | 174.1 | 174.1 | 114.3 | 114.3 | 8.19 | 8.20 | 4.22 | 70.1 | 18-26 |
| 371 | Y | 57.4 | 57.9 | 176.0 | 176.0 | 121.9 | 121.8 | 8.19 | 8.20 | 4.58 | 39.0 | 18-26 |
| 372 | G | 44.7 | 45.2 | 173.9 | 174.0 | 110.6 | 110.6 | 8.36 | 8.37 | 3.91 | - | 18-26 |
| 373 | V | - | 62.7 | 177.0 | 176.9 | 119.3 | 119.2 | 8.11 | 8.12 | 4.10 | 32.3 | 18-26 |
| 374 | G | 44.8 | 45.2 | 174.0 | 174.0 | - | 113.0 | - | 8.63 | 3.93 | - | 18-26 |
| 375 | A | 52.3 | 52.8 | 178.3 | 178.3 | 124.0 | 123.9 | 8.29 | 8.29 | 4.28 | 19.2 | 18-26 |
| 376 | G | 44.9 | 45.3 | 174.5 | 174.5 | 108.3 | 108.2 | - | 8.50 | 3.87 | - | 18-26 |
| 377 | G | 44.4 | 45.0 | 173.3 | 173.3 | 108.1 | 108.0 | 8.13 | 8.13 | 3.81 | - | 18-26 |
| 378 | F | 55.2 | 55.6 | 174.2 | 174.2 | 121.0 | 121.0 | 8.13 | 8.13 | 4.84 | 39.1 | 18-26 |
| 379 | P | 63.2 | 63.8 | 177.1 | 177.1 | - | - | - | - | 4.35 | 31.9 | 18-26 |
| 380 | G | 44.5 | 44.9 | 173.7 | 173.7 | 108.5 | 108.5 | 7.86 | 7.87 | 3.74 | - | 18-26 |
| 381 | F | 57.7 | 58.1 | 176.3 | 176.3 | 119.8 | 119.7 | 8.18 | 8.19 | 4.58 | 39.7 | 18-26 |
| 382 | G | 44.8 | 45.3 | 173.8 | 173.9 | 111.0 | 111.0 | 8.47 | 8.48 | 3.87 | - | 18-26 |
| 383 | V | - | 62.5 | - | 176.7 | 119.3 | 119.1 | - | 8.03 | 4.11 | 32.7 | 18-26 |
| 384 | G | - | 45.3 | - | 174.3 | - | 112.9 | - | 8.58 | 3.92 | - | 18-26 |
| 385 | V | - | 62.7 | 176.9 | 176.9 | 119.4 | 119.3 | - | 8.11 | 4.11 | 32.6 | 18-26 |
| 386 | G | - | 45.3 | 174.5 | 174.5 | 112.6 | 112.6 | 8.64 | 8.65 | 3.94 | - | 18-26 |
| 387 | G | 44.5 | 45.3 | 173.6 | 173.6 | 108.6 | 108.4 | - | 8.23 | 3.90 | - | 18-26 |
| 388 | I | 58.3 | 58.6 | 174.6 | 174.6 | 122.1 | 122.0 | 8.15 | 8.15 | 4.41 | 38.6 | 18-26 |
| 389 | P | - | 63.7 | - | 177.4 | - | - | - | - | 4.33 | 32.2 | 18-26 |
| 390 | G | - | 45.2 | 173.9 | 174.0 | 109.7 | 109.7 | 8.46 | 8.47 | 3.95 | - | 18-26 |
| 391 | V | 61.8 | 62.2 | 175.9 | 175.9 | 120.0 | 119.9 | 8.03 | 8.03 | 4.11 | 33.1 | 18-26 |
| 392 | A | 52.1 | 52.5 | 178.0 | 178.0 | 128.1 | 128.0 | 8.56 | 8.57 | 4.28 | 19.2 | 18-26 |
| 393 | G | 44.7 | 44.9 | 173.6 | 173.6 | 108.2 | 108.3 | 8.38 | 8.39 | 3.91 | - | 18-26 |
| 394 | V | - | - | - | - | 121.5 | 121.6 | - | - | - | - | 18-26 |
| 395 | P | - | - | - | - | - | - | - | - | - | - | - |
| 396 | G | - | - | - | 174.2 | - | - | - | - | - | - | 18-26 |
| 397 | V | - | - | - | 177.0 | - | 119.6 | - | 8.13 | - | - | 18-26 |
| 398 | G | - | - | - | 174.6 | - | 113.1 | - | 8.72 | - | - | 18-26 |
| 399 | G | - | - | - | 173.7 | - | 108.6 | - | 8.33 | - | - | 18-26 |
| 400 | V | - | - | - | 174.7 | - | 121.6 | - | 8.09 | - | - | 18-26 |
| 401 | P | - | - | - | - | - | - | - | - | - | - | - |
| 402 | G | - | - | - | 174.2 | - | - | - | - | - | - | 18-26 |
| 403 | V | - | - | - | 177.0 | - | 119.6 | - | 8.13 | - | - | 18-26 |
| 404 | G | - | - | - | 174.6 | - | 113.1 | - | 8.72 | - | - | 18-26 |
| 405 | G | - | - | - | 173.7 | - | 108.6 | - | 8.33 | - | - | 18-26 |
| 406 | V | - | - | - | 174.7 | - | 121.6 | - | 8.09 | - | - | 18-26 |
| 407 | P | - | - | - | - | - | - | - | - | - | - | - |
| 408 | G | - | - | - | - | - | - | - | - | - | - | - |
| 409 | V | - | 62.3 | - | 176.7 | - | - | - | - | 4.15 | 32.5 | 18-26 |
| 410 | G | 44.8 | 45.3 | 173.6 | 173.7 | 112.8 | 112.7 | 8.59 | 8.59 | 3.92 | - | 18-26 |

| Residue | | $^{13}C_{\alpha}$ | | $^{13}CO$ | | $^{15}N$ | | $^{1}HN$ | | $H_{\alpha}$ | $C_{\beta}$ | Source |
|---|---|---|---|---|---|---|---|---|---|---|---|---|
| # | AA | hTE | Frag. | hTE | Frag. | hTE | Frag. | hTE | Frag. | Frag. | Frag. | Frag. |
| 411 | I | 60.3 | 60.7 | 176.1 | 176.1 | 119.8 | 119.6 | 8.07 | 8.06 | 4.26 | 38.9 | 18-26 |
| 412 | S | 56.2 | 56.8 | 173.1 | 173.1 | 123.0 | 122.9 | 8.68 | 8.68 | 4.73 | 63.3 | 18-26 |
| 413 | P | 64.8 | 65.3 | 179.4 | 179.4 | - | 137.3 | - | - | 4.29 | 31.8 | 18-26 |
| 414 | E | 59.0 | 59.6 | 178.6 | 178.6 | 119.3 | 119.2 | 8.90 | 8.90 | 4.10 | 28.9 | 18-26 |
| 415 | A | 54.2 | 54.6 | 180.9 | 180.8 | 125.4 | 125.3 | 8.19 | 8.19 | 4.25 | 18.3 | 18-26 |
| 416 | Q | 58.1 | 58.6 | 178.4 | 178.4 | 120.5 | 120.4 | 8.52 | 8.53 | 4.09 | 28.5 | 18-26 |
| 417 | A | 54.0 | 54.7 | 180.2 | 180.2 | 123.2 | 123.4 | 8.26 | 8.27 | 4.21 | 17.9 | 18-26 |
| 418 | A | 54.1 | 54.5 | 180.1 | 180.1 | - | 122.3 | - | 8.09 | 4.21 | 17.8 | 18-26 |
| 419 | A | 53.9 | 54.5 | 180.1 | 180.1 | 121.8 | 121.9 | 8.02 | 8.03 | 4.19 | 18.1 | 18-26 |
| 420 | A | - | 54.3 | 179.8 | 179.8 | 122.3 | 122.3 | 8.09 | 8.13 | 4.17 | 18.0 | 18-26 |
| 421 | A | 53.8 | 53.9 | 179.7 | 179.7 | 121.7 | 121.8 | 8.00 | 8.01 | 4.17 | 18.3 | 18-26 |
| 422 | K | 57.9 | 58.4 | 177.8 | 177.8 | 119.3 | 119.3 | 7.85 | 7.86 | 4.26 | 32.9 | 18-26 |
| 423 | A | 53.2 | 53.6 | 178.8 | 178.9 | 122.0 | 122.0 | 7.91 | 7.93 | 4.17 | 18.4 | 18-26 |
| 424 | A | 52.9 | 53.5 | 178.4 | 178.4 | 121.0 | 121.0 | 7.87 | 7.88 | 4.19 | 18.5 | 18-26 |
| 425 | K | 56.8 | 57.4 | 176.8 | 176.8 | 118.9 | 118.8 | 7.81 | 7.81 | 4.24 | 32.9 | 18-26 |
| 426 | Y | 57.7 | 58.1 | 176.5 | 176.5 | 118.5 | 118.4 | 8.00 | 8.00 | 4.60 | 38.8 | 18-26 |
| 427 | G | - | 45.3 | 174.0 | 174.0 | 109.9 | 109.9 | 8.19 | 8.21 | 3.96 | - | 18-26 |
| 428 | V | - | 62.6 | 176.9 | 177.0 | 119.4 | 119.4 | 8.14 | 8.15 | 4.12 | 32.7 | 18-26 |
| 429 | G | 44.8 | 45.3 | 174.1 | 174.1 | 112.8 | 112.8 | 8.68 | 8.72 | 3.98 | - | 18-26 |
| 430 | T | 60.1 | 60.2 | 173.6 | 173.7 | 114.4 | 114.1 | 7.98 | 7.95 | 4.68 | 69.8 | 18-26 |
| 431 | P | 63.9 | 64.6 | 178.1 | 178.3 | - | - | - | - | 4.37 | 32.0 | 18-26 |
| 432 | A | 53.4 | 54.0 | 179.2 | 179.3 | 122.7 | 122.5 | 8.39 | 8.40 | 4.18 | 18.6 | 18-26 |
| 433 | A | 53.4 | 54.3 | 179.4 | 179.5 | 122.9 | 122.9 | 8.20 | 8.18 | 4.19 | 18.2 | 18-26 |
| 434 | A | 53.8 | 53.6 | 179.7 | 179.8 | 122.4 | 122.4 | 8.14 | 8.13 | 4.17 | 18.3 | 18-26 |
| 435 | A | 53.4 | 54.1 | 179.4 | 179.6 | 122.6 | 122.6 | 8.24 | 8.25 | 4.21 | 18.0 | 18-26 |
| 436 | A | 53.5 | 54.0 | 179.7 | 179.9 | 122.7 | 122.7 | 8.10 | 8.08 | 4.20 | 18.1 | 18-26 |
| 437 | K | 57.7 | 58.3 | 178.1 | 178.3 | 120.5 | 120.5 | 8.09 | 8.09 | 4.13 | 32.5 | 18-26 |
| 438 | A | 53.4 | 54.1 | 179.3 | 179.6 | 123.2 | 123.0 | 8.09 | 8.08 | 4.20 | 18.2 | 18-26 |
| 439 | A | 52.9 | 54.0 | 179.0 | 179.3 | 122.4 | 122.3 | 8.19 | 8.18 | 4.22 | 18.1 | 18-26 |
| 440 | A | 53.4 | 54.1 | 179.1 | 179.3 | 122.5 | 122.3 | 8.04 | 8.01 | 4.20 | 18.2 | 18-26 |
| 441 | K | 57.1 | 57.9 | 177.6 | 177.8 | 119.9 | 119.7 | 8.04 | 8.01 | 4.10 | 32.6 | 18-26 |
| 442 | A | 52.9 | 53.6 | 178.5 | 178.7 | 123.1 | 122.8 | 8.08 | 8.05 | 4.15 | 18.5 | 18-26 |
| 443 | A | 52.7 | 53.3 | 178.2 | 178.4 | 121.7 | 121.3 | 8.03 | 7.99 | 4.20 | 18.6 | 18-26 |
| 444 | Q | 55.9 | 56.4 | 176.1 | 176.2 | 118.3 | 118.1 | 7.98 | 7.96 | 4.16 | 29.0 | 18-26 |
| 445 | F | 57.9 | 58.2 | 176.3 | 176.3 | 120.0 | 119.7 | 8.14 | 8.11 | 4.60 | 39.3 | 18-26 |
| 446 | G | 44.9 | 45.3 | 173.7 | 173.7 | 110.1 | 110.1 | 8.23 | 8.23 | 3.87 | - | 18-26 |
| 447 | L | 54.7 | 55.1 | 177.2 | 177.2 | 121.4 | 121.2 | 8.03 | 8.03 | 4.34 | 42.4 | 18-26 |
| 448 | V | 59.5 | 59.9 | 174.4 | 174.3 | 123.4 | 123.2 | 8.28 | 8.28 | 4.40 | 32.6 | 18-26 |
| 449 | P | - | - | - | - | - | 139.8 | - | - | - | - | 18-26 |
| 450 | G | 44.8 | - | 174.0 | - | 109.6 | 109.5 | 8.52 | - | - | - | 18-26 |
| 451 | V | 62.2 | 62.5 | 176.8 | 176.7 | 119.6 | 119.5 | 8.05 | 8.04 | 4.12 | 32.4 | 18-26 |

| Residue | | $^{13}C_{\alpha}$ | | $^{13}CO$ | | $^{15}N$ | | $^{1}HN$ | | $H_{\alpha}$ | $C_{\beta}$ | Source |
|---|---|---|---|---|---|---|---|---|---|---|---|---|
| # | AA | hTE | Frag. | hTE | Frag. | hTE | Frag. | hTE | Frag. | Frag. | Frag. | Frag. |
| 452 | G | 44.8 | 45.2 | 173.7 | 173.7 | 113.0 | 113.0 | 8.60 | 8.61 | 3.94 | - | 18-26 |
| 453 | V | 61.4 | 61.8 | 175.7 | 175.6 | 119.6 | 119.4 | 8.01 | 8.01 | 4.11 | 32.8 | 18-26 |
| 454 | A | 50.1 | 50.4 | 175.4 | 175.3 | 130.0 | 129.9 | 8.54 | 8.54 | 4.59 | 18.1 | 18-26 |
| 455 | P | - | 63.4 | - | 177.6 | - | 135.2 | - | - | 4.38 | 32.1 | 18-26 |
| 456 | G | 44.8 | 45.2 | 174.0 | 174.1 | 109.6 | 109.5 | 8.52 | 8.53 | 3.94 | - | 18-26 |
| 457 | V | 62.2 | 62.5 | 176.8 | 176.7 | 119.6 | 119.5 | 8.05 | 8.04 | 4.12 | 32.4 | 18-26 |
| 458 | G | 44.8 | 45.2 | 173.7 | 173.7 | 113.0 | 113.0 | 8.60 | 8.61 | 3.94 | - | 18-26 |
| 459 | V | 61.4 | 61.8 | 175.7 | 175.6 | 119.6 | 119.4 | 8.01 | 8.01 | 4.11 | 32.8 | 18-26 |
| 460 | A | 50.1 | 50.4 | 175.4 | 175.3 | 130.0 | 129.9 | 8.54 | 8.54 | 4.59 | 18.1 | 18-26 |
| 461 | P | - | - | - | 177.6 | - | - | - | - | 4.38 | - | 18-26 |
| 462 | G | 44.8 | 45.2 | 174.0 | 174.1 | 109.6 | 109.5 | 8.52 | 8.53 | 3.94 | - | 18-26 |
| 463 | V | 62.2 | 62.5 | 176.8 | 176.7 | 119.6 | 119.5 | 8.05 | 8.05 | 4.12 | 32.4 | 18-26 |
| 464 | G | 44.8 | 45.2 | 173.7 | 173.7 | 113.0 | 113.0 | 8.60 | 8.61 | 3.94 | - | 18-26 |
| 465 | V | 61.4 | 61.8 | 175.7 | 175.6 | 119.6 | 119.4 | 8.01 | 8.01 | 4.11 | 32.8 | 18-26 |
| 466 | A | 50.1 | 50.4 | 175.4 | 175.3 | 130.0 | 129.9 | 8.54 | 8.54 | 4.59 | 18.1 | 18-26 |
| 467 | P | - | - | - | 177.6 | - | 135.2 | - | - | - | - | 18-26 |
| 468 | G | - | - | - | - | - | - | - | - | - | - | - |
| 469 | V | - | - | - | 176.7 | - | 119.5 | - | 8.04 | - | - | 18-26 |
| 470 | G | 44.6 | 45.3 | 173.7 | 173.7 | 112.8 | 112.7 | - | 8.58 | - | - | 18-26 |
| 471 | L | 54.2 | 54.6 | 176.9 | 176.9 | 121.7 | 121.7 | 8.14 | 8.13 | 4.33 | 42.6 | 18-26 |
| 472 | A | 50.1 | 50.6 | 175.3 | 175.3 | 126.6 | 126.6 | 8.41 | 8.42 | 4.57 | 18.0 | 18-26 |
| 473 | P | - | 63.4 | - | 177.6 | - | 135.3 | - | - | 4.38 | 32.1 | 18-26 |
| 474 | G | 44.8 | 45.2 | 174.0 | 174.1 | 109.6 | 109.5 | 8.52 | 8.53 | 3.94 | - | 18-26 |
| 475 | V | 62.2 | 62.5 | 176.8 | 176.7 | 119.6 | 119.5 | 8.05 | 8.04 | 4.12 | 32.4 | 18-26 |
| 476 | G | 44.8 | 45.2 | 173.7 | 173.7 | 113.0 | 113.0 | 8.60 | 8.61 | 3.94 | - | 18-26 |
| 477 | V | 61.4 | 61.8 | 175.7 | 175.6 | 119.6 | 119.4 | 8.01 | 8.01 | 4.11 | 32.8 | 18-26 |
| 478 | A | 50.1 | 50.4 | 175.4 | 175.3 | 130.0 | 129.9 | 8.54 | 8.54 | 4.59 | 18.1 | 18-26 |
| 479 | P | - | 63.4 | - | 177.6 | - | 135.2 | - | - | 4.38 | 32.1 | 18-26 |
| 480 | G | 44.8 | 45.2 | 174.0 | 174.1 | 109.6 | 109.5 | 8.52 | 8.53 | 3.94 | - | 18-26 |
| 481 | V | 62.2 | 62.5 | 176.8 | 176.7 | 119.6 | 119.5 | 8.05 | 8.04 | 4.12 | 32.4 | 18-26 |
| 482 | G | 44.8 | 45.2 | 173.7 | 173.7 | 113.0 | 113.0 | 8.60 | 8.61 | 3.94 | - | 18-26 |
| 483 | V | 61.4 | 61.8 | 175.7 | 175.6 | 119.6 | 119.4 | 8.01 | 8.01 | 4.11 | 32.8 | 18-26 |
| 484 | A | 50.1 | 50.4 | 175.4 | 175.3 | 130.0 | 129.9 | 8.54 | 8.54 | 4.59 | 18.1 | 18-26 |
| 485 | P | - | 63.4 | - | 177.6 | - | 135.2 | - | - | 4.38 | 32.1 | 18-26 |
| 486 | G | 44.8 | 45.2 | 174.0 | 174.1 | 109.6 | 109.5 | 8.52 | 8.53 | 3.94 | - | 18-26 |
| 487 | V | 62.2 | 62.5 | 176.8 | 176.7 | 119.6 | 119.5 | 8.05 | 8.04 | 4.12 | 32.4 | 18-26 |
| 488 | G | 44.8 | 45.2 | 173.7 | 173.7 | 113.0 | 113.0 | 8.60 | 8.61 | 3.94 | - | 18-26 |
| 489 | V | 61.4 | 61.8 | 175.7 | 175.6 | 119.6 | 119.4 | 8.01 | 8.01 | 4.11 | 32.8 | 18-26 |
| 490 | A | 50.1 | 50.4 | 175.4 | 175.3 | 130.0 | 129.9 | 8.54 | 8.54 | 4.59 | 18.1 | 18-26 |
| 491 | P | - | 63.3 | - | 177.6 | - | - | - | - | 4.39 | 32.2 | 18-26 |
| 492 | G | 44.8 | 45.2 | 173.9 | 173.9 | - | 109.2 | - | 8.50 | 3.92 | - | 18-26 |

| Residue | | $^{13}C_{\alpha}$ | | $^{13}CO$ | | $^{15}N$ | | $^{1}HN$ | | $H_{\alpha}$ | $C_{\beta}$ | Source |
|---|---|---|---|---|---|---|---|---|---|---|---|---|
| # | AA | hTE | Frag. | hTE | Frag. | hTE | Frag. | hTE | Frag. | Frag. | Frag. | Frag. |
| 493 | I | 60.6 | 61.2 | 176.5 | 176.5 | 120.0 | 119.9 | 8.05 | 8.04 | 4.22 | 38.9 | 18-26 |
| 494 | G | 44.3 | 44.7 | 172.1 | 172.1 | 113.2 | 113.2 | 8.45 | 8.45 | 4.07 | - | 18-26 |
| 495 | P | 63.5 | 63.9 | 178.2 | 178.2 | - | - | - | - | 4.38 | 31.8 | 18-26 |
| 496 | G | 45.0 | 45.4 | 175.1 | 175.1 | 109.7 | 109.7 | 8.75 | 8.76 | 3.96 | - | 18-26 |
| 497 | G | 45.2 | 45.4 | 174.6 | 174.7 | 109.0 | 109.0 | 8.25 | 8.25 | 3.96 | - | 18-26 |
| 498 | V | 63.0 | 63.4 | 176.8 | 176.9 | 120.5 | 120.5 | 8.05 | 8.06 | 3.99 | 32.5 | 18-26 |
| 499 | A | 52.8 | 53.4 | 178.5 | 178.6 | 126.8 | 126.7 | 8.50 | 8.51 | 4.26 | 18.7 | 18-26 |
| 500 | A | 52.9 | 53.5 | 178.6 | 178.7 | 123.0 | 123.0 | 8.25 | 8.25 | 4.21 | 18.5 | 18-26 |
| 501 | A | 52.8 | 53.5 | 178.6 | 178.7 | 122.9 | 122.8 | 8.23 | 8.24 | 4.21 | 18.5 | 18-26 |
| 502 | A | 52.9 | 53.5 | 179.0 | 179.1 | 123.1 | 122.9 | 8.20 | 8.19 | 4.24 | 18.5 | 18-26 |
| 503 | K | 56.9 | 57.6 | 177.7 | 177.8 | 120.7 | 120.7 | 8.25 | 8.24 | 4.24 | 32.7 | 18-26 |
| 504 | S | 59.0 | 59.6 | 175.2 | 175.3 | 116.5 | 116.4 | 8.28 | 8.27 | 4.35 | 63.4 | 18-26 |
| 505 | A | 53.0 | 53.6 | 178.7 | 178.9 | 125.4 | 125.4 | 8.35 | 8.35 | 4.26 | 18.6 | 18-26 |
| 506 | A | 53.0 | 53.6 | 178.9 | 179.1 | 122.7 | 122.5 | 8.19 | 8.19 | 4.23 | 18.5 | 18-26 |
| 507 | K | 57.1 | 57.9 | 177.7 | 177.9 | 121.0 | 120.9 | 8.14 | 8.10 | 4.39 | 32.6 | 18-26 |
| 508 | V | 63.4 | 64.2 | 177.1 | 177.3 | 121.7 | 121.5 | 8.10 | 8.09 | 3.87 | 32.6 | 18-26 |
| 509 | A | 53.0 | 53.5 | 178.4 | 178.6 | 126.2 | 125.9 | 8.30 | 8.29 | 4.24 | 18.6 | 18-26 |
| 510 | A | 52.8 | 53.7 | 178.9 | 179.1 | 122.8 | 122.7 | 8.25 | 8.23 | 4.23 | 18.5 | 18-26 |
| 511 | K | 57.0 | 57.8 | 177.4 | 177.6 | 120.2 | 120.0 | 8.18 | 8.15 | 4.16 | 32.6 | 18-26 |
| 512 | A | 52.9 | 53.7 | 178.7 | 179.0 | 124.0 | 123.7 | 8.20 | 8.17 | 4.21 | 18.4 | 18-26 |
| 513 | Q | 56.2 | 57.0 | 176.8 | 177.0 | 119.4 | 119.3 | 8.24 | 8.23 | 4.22 | 28.9 | 18-26 |
| 514 | L | 55.7 | 56.5 | 178.1 | 178.4 | 123.0 | 122.6 | 8.16 | 8.11 | 4.22 | 42.0 | 18-26 |
| 515 | R | 56.4 | 57.1 | 176.7 | 177.0 | 121.4 | 121.1 | 8.26 | 8.22 | 4.22 | 30.5 | 18-26 |
| 516 | A | 52.6 | 53.4 | 178.2 | 178.5 | 124.5 | 124.1 | 8.25 | 8.21 | 4.23 | 18.6 | 18-26 |
| 517 | A | 52.4 | 53.0 | 177.9 | 178.0 | 122.8 | 122.6 | 8.23 | 8.20 | 4.24 | 18.4 | 18-26 |
| 518 | A | 52.3 | 52.9 | 178.3 | 178.4 | 122.5 | 122.2 | 8.17 | 8.11 | 4.27 | 19.1 | 18-26 |
| 519 | G | 44.8 | 45.4 | 174.4 | 174.5 | 107.6 | 107.5 | 8.28 | 8.25 | 3.96 | - | 18-26 |
| 520 | L | 54.9 | 55.3 | 178.2 | 178.1 | 121.6 | 121.5 | 8.17 | 8.15 | 4.38 | 42.5 | 18-26 |
| 521 | G | - | 45.4 | 173.8 | 173.9 | 109.9 | 109.8 | 8.53 | 8.53 | 3.92 | - | 18-26 |
| 522 | A | 52.2 | 52.5 | 178.2 | 178.3 | 123.7 | 123.6 | 8.23 | 8.23 | 4.31 | 19.2 | 18-26 |
| 523 | G | 44.6 | 45.1 | 173.7 | 173.7 | 108.2 | 108.2 | 8.44 | 8.46 | 3.90 | - | 18-26 |
| 524 | I | 58.4 | 58.8 | 174.8 | 174.8 | 122.2 | 122.1 | 8.09 | 8.09 | 4.42 | 38.4 | 18-26 |
| 525 | P | - | - | - | - | - | - | - | - | - | - | - |
| 526 | G | - | - | 174.1 | - | - | - | - | - | - | - | - |
| 527 | L | - | - | - | - | 121.5 | - | 8.17 | - | - | - | - |
| 528 | G | - | - | - | - | - | - | - | - | - | - | - |
| 529 | V | - | - | - | - | - | - | - | - | - | - | - |
| 530 | G | - | 45.2 | - | 173.9 | - | 112.8 | - | 8.62 | - | - | 24-36 |
| 531 | V | - | 62.4 | - | 176.7 | - | 119.0 | - | 8.04 | - | 32.6 | 24-36 |
| 532 | G | - | 44.9 | - | 173.6 | - | 112.7 | - | 8.58 | - | - | 24-36 |
| 533 | V | - | 59.9 | - | 174.6 | - | 121.3 | - | 8.07 | - | 32.3 | 24-36 |

| Residue | | $^{13}C_{\alpha}$ | | $^{13}CO$ | | $^{15}N$ | | $^{1}HN$ | | $H_{\alpha}$ | $C_{\beta}$ | Source |
|---|---|---|---|---|---|---|---|---|---|---|---|---|
| **#** | **AA** | **hTE** | **Frag.** | **hTE** | **Frag.** | **hTE** | **Frag.** | **hTE** | **Frag.** | **Frag.** | **Frag.** | **Frag.** |
| 534 | P | - | - | - | - | - | - | - | - | - | - | - |
| 535 | G | - | - | - | - | - | 109.3 | - | 8.53 | - | - | 24-36 |
| 536 | L | - | 55.2 | - | 178.0 | - | 121.4 | - | 8.20 | - | 42.3 | 24-36 |
| 537 | G | - | 45.1 | - | 174.1 | - | 110.0 | - | 8.56 | - | - | 24-36 |
| 538 | V | - | - | - | 176.9 | - | 119.2 | - | 8.10 | - | - | 24-36 |
| 539 | G | - | - | - | - | - | 112.8 | - | - | - | - | 24-36 |
| 540 | A | - | - | - | - | - | 123.5 | - | 8.17 | - | - | 24-36 |
| 541 | G | - | - | - | - | - | 108.1 | - | 8.45 | - | - | 24-36 |
| 542 | V | - | - | - | - | - | - | - | - | - | - | - |
| 543 | P | - | - | - | - | - | - | - | - | - | - | - |
| 544 | G | - | - | - | - | - | 109.3 | - | 8.53 | - | - | 24-36 |
| 545 | L | - | 55.2 | - | 178.0 | - | 121.4 | - | 8.20 | - | 42.3 | 24-36 |
| 546 | G | - | 45.1 | - | 174.1 | - | 110.0 | - | 8.56 | - | - | 24-36 |
| 547 | V | - | - | - | 176.9 | - | 119.2 | - | 8.10 | - | - | 24-36 |
| 548 | G | - | 45.2 | - | 173.8 | - | 112.8 | - | - | - | - | 24-36 |
| 549 | A | - | 52.5 | - | 178.2 | - | 123.5 | - | 8.17 | - | 19.1 | 24-36 |
| 550 | G | - | 44.9 | 173.7 | 173.7 | 108.4 | 108.2 | - | 8.45 | - | - | 24-36 |
| 551 | V | - | 60.0 | 174.7 | 174.6 | 121.3 | 121.2 | 8.06 | 8.06 | - | 32.4 | 24-36 |
| 552 | P | - | 63.6 | - | 177.4 | - | - | - | - | - | 32.0 | 24-36 |
| 553 | G | 44.6 | 45.1 | 174.0 | 174.0 | - | 109.7 | - | 8.54 | 3.88 | - | 24-36 |
| 554 | F | 58.1 | 58.5 | 176.4 | 176.4 | 120.5 | 120.3 | 8.21 | 8.20 | 4.50 | 39.6 | 24-36 |
| 555 | G | 44.6 | 45.1 | 173.5 | 173.5 | 111.5 | 111.5 | 8.45 | 8.45 | 3.81 | - | 24-36 |
| 556 | A | 51.9 | 52.3 | 177.6 | 177.6 | 123.7 | 123.7 | 8.08 | 8.08 | 4.26 | 19.1 | 24-36 |
| 557 | V | 59.9 | 60.2 | 174.8 | 174.7 | 121.4 | 121.4 | 8.28 | 8.28 | 4.34 | 32.1 | 24-36 |
| 558 | P | 63.4 | 63.9 | 177.8 | 177.8 | - | - | - | - | 4.32 | 31.9 | 24-36 |
| 559 | G | 44.9 | 45.3 | 174.4 | 174.4 | 110.0 | 109.9 | 8.64 | 8.63 | 3.91 | - | 24-36 |
| 560 | A | 53.0 | 53.5 | 178.8 | 178.8 | 123.7 | 123.7 | 8.14 | 8.14 | 4.20 | 19.0 | 24-36 |
| 561 | L | 55.4 | 55.9 | 178.0 | 178.0 | 121.0 | 121.0 | 8.27 | 8.28 | 4.24 | 42.0 | 24-36 |
| 562 | A | 52.6 | 53.2 | 178.6 | 178.6 | 124.2 | 124.2 | 8.25 | 8.25 | 4.17 | 18.6 | 24-36 |
| 563 | A | 52.9 | 53.3 | 178.5 | 178.5 | 122.6 | 122.6 | 8.22 | 8.22 | 4.19 | 18.5 | 24-36 |
| 564 | A | 52.9 | 53.3 | 178.7 | 178.7 | 122.8 | 122.8 | 8.15 | 8.15 | 4.19 | 18.6 | 24-36 |
| 565 | K | 56.8 | 57.3 | 177.0 | 177.0 | 120.1 | 120.1 | 8.16 | 8.16 | 4.10 | 32.9 | 24-36 |
| 566 | A | 52.4 | 52.9 | 178.1 | 178.0 | 123.8 | 123.8 | 8.15 | 8.14 | 4.19 | 18.7 | 24-36 |
| 567 | A | 52.4 | 52.8 | 178.0 | 177.9 | 122.7 | 122.7 | 8.13 | 8.12 | 4.18 | 18.7 | 24-36 |
| 568 | K | 56.4 | 56.7 | 176.6 | 176.5 | 119.9 | 119.9 | 8.08 | 8.07 | 4.15 | 32.7 | 24-36 |
| 569 | Y | 57.8 | 58.1 | 176.5 | 176.5 | 120.3 | 120.3 | 8.19 | 8.19 | 4.53 | 38.7 | 24-36 |
| 570 | G | 44.8 | 45.2 | 173.5 | 173.5 | 110.7 | 110.8 | 8.26 | 8.27 | 3.85 | - | 24-36 |
| 571 | A | 51.9 | 52.3 | 177.3 | 177.3 | 123.7 | 123.6 | 8.08 | 8.07 | 4.25 | 19.3 | 24-36 |
| 572 | A | 51.8 | 52.3 | 177.5 | 177.5 | 123.5 | 123.5 | 8.32 | 8.32 | 4.26 | 19.0 | 24-36 |
| 573 | V | 59.5 | 59.8 | 174.5 | 174.5 | 121.7 | 121.7 | 8.24 | 8.25 | 4.33 | 32.4 | 24-36 |
| 574 | P | - | - | - | - | - | - | - | - | - | - | - |

| Residue | | $^{13}C_{\alpha}$ | | $^{13}CO$ | | $^{15}N$ | | $^{1}HN$ | | $H_{\alpha}$ | $C_{\beta}$ | Source |
|---|---|---|---|---|---|---|---|---|---|---|---|---|
| # | AA | hTE | Frag. | hTE | Frag. | hTE | Frag. | hTE | Frag. | Frag. | Frag. | Frag. |
| 575 | G | 44.8 | - | 174.1 | 174.1 | 109.5 | 109.4 | - | - | - | - | 24-36 |
| 576 | V | 62.1 | 62.5 | 176.5 | 176.4 | 119.7 | 119.6 | 8.04 | 8.03 | - | 32.7 | 24-36 |
| 577 | L | 55.0 | 55.3 | 177.9 | 177.9 | 126.0 | 126.1 | 8.51 | 8.51 | 4.31 | 42.0 | 24-36 |
| 578 | G | 44.9 | 45.2 | 174.6 | 174.6 | 110.1 | 110.1 | 8.48 | 8.49 | 3.92 | - | 24-36 |
| 579 | G | 44.8 | 45.2 | 174.4 | 174.4 | 108.6 | 108.5 | 8.27 | 8.30 | - | - | 24-36 |
| 580 | L | 55.1 | 55.7 | 178.3 | 178.2 | 121.5 | 121.5 | 8.33 | 8.33 | - | 42.1 | 24-36 |
| 581 | G | - | 45.3 | 174.1 | 174.0 | 109.7 | 109.6 | 8.55 | 8.53 | - | - | 24-36 |
| 582 | A | 52.3 | 52.6 | 178.1 | 178.1 | 123.8 | 123.7 | 8.19 | 8.18 | - | 19.0 | 24-36 |
| 583 | L | 54.9 | 55.4 | 178.1 | 178.1 | 120.9 | 121.0 | 8.31 | 8.31 | - | 42.0 | 24-36 |
| 584 | G | 45.1 | 45.3 | 174.5 | 174.5 | 109.3 | 109.3 | 8.37 | 8.38 | - | - | 24-36 |
| 585 | G | - | 45.1 | - | 174.0 | 108.8 | 108.6 | - | 8.27 | 3.92 | - | 24-36 |
| 586 | V | - | 62.4 | 176.7 | 176.7 | - | 118.8 | - | 8.07 | 4.09 | 32.6 | 24-36 |
| 587 | G | - | 44.9 | 173.6 | 173.5 | 112.5 | 112.6 | 8.55 | 8.55 | 3.88 | - | 24-36 |
| 588 | I | 58.2 | 58.6 | 174.8 | 174.8 | 121.7 | 121.7 | 8.09 | 8.09 | 4.41 | 38.6 | 24-36 |
| 589 | P | 63.4 | 63.7 | 177.6 | 177.6 | - | - | - | - | 4.35 | 31.9 | 24-36 |
| 590 | G | 44.9 | 45.2 | 174.7 | 174.7 | 110.5 | 110.5 | 8.66 | 8.66 | 3.93 | - | 24-36 |
| 591 | G | 44.9 | 45.0 | 173.8 | 173.8 | 109.0 | 108.7 | - | 8.30 | 3.93 | - | 24-36 |
| 592 | V | 62.0 | 62.4 | 176.4 | 176.4 | 119.8 | 119.8 | 8.15 | 8.15 | 4.08 | 32.7 | 24-36 |
| 593 | V | 62.4 | 62.7 | 176.8 | 176.7 | 125.4 | 125.5 | 8.42 | 8.42 | 4.05 | 32.4 | 24-36 |
| 594 | G | 44.7 | 45.0 | 173.6 | 173.6 | 113.1 | 113.2 | 8.59 | 8.59 | 3.91 | - | 24-36 |
| 595 | A | 52.4 | 52.6 | 178.4 | 178.4 | 123.7 | 123.6 | 8.29 | 8.29 | 4.31 | 19.4 | 24-36 |
| 596 | G | 45.0 | 45.4 | 173.1 | 173.1 | 108.5 | 108.5 | 8.55 | 8.56 | 4.06 | - | 24-36 |
| 597 | P | 64.3 | 64.7 | 178.7 | 178.7 | - | - | - | - | - | 31.9 | 24-36 |
| 598 | A | 53.9 | 54.4 | 179.8 | 179.9 | 122.8 | 122.8 | 8.41 | 8.40 | - | - | 24-36 |
| 599 | A | 54.0 | - | 180.1 | 180.1 | 123.2 | 123.3 | 8.23 | 8.23 | - | - | 24-36 |
| 600 | A | 53.9 | - | 180.2 | 180.1 | 122.7 | 122.7 | 8.24 | 8.24 | - | - | 24-36 |
| 601 | A | - | - | 180.1 | 180.1 | - | 122.6 | - | 8.19 | - | - | 24-36 |
| 602 | A | - | - | 180.4 | 180.3 | 122.2 | 122.2 | 8.10 | 8.09 | - | - | 24-36 |
| 603 | A | - | 54.6 | 180.1 | 180.1 | 122.5 | 122.5 | 8.19 | 8.19 | - | - | 24-36 |
| 604 | A | 54.2 | 54.7 | 180.3 | 180.3 | 122.6 | 122.4 | 8.09 | 8.08 | - | 17.8 | 24-36 |
| 605 | K | 58.4 | 58.9 | 178.6 | 178.6 | 120.5 | 120.4 | 8.03 | 8.03 | - | 32.3 | 24-36 |
| 606 | A | 53.9 | 54.5 | 179.9 | 179.9 | 122.6 | 122.5 | 8.03 | 8.02 | - | 17.9 | 24-36 |
| 607 | A | 53.7 | 54.2 | 179.6 | 179.6 | 122.0 | 122.1 | 8.16 | 8.16 | - | 17.9 | 24-36 |
| 608 | A | 53.7 | 54.2 | 179.6 | 179.6 | 122.0 | 122.0 | 7.97 | 7.95 | - | 17.9 | 24-36 |
| 609 | K | 58.0 | 58.1 | 178.0 | 178.0 | 119.5 | 119.6 | 7.96 | 7.94 | - | 32.5 | 24-36 |
| 610 | A | 53.4 | 53.8 | 178.9 | 178.9 | 122.4 | 122.4 | 8.01 | 8.00 | - | 18.1 | 24-36 |
| 611 | A | 53.0 | 53.4 | 178.5 | 178.6 | 121.1 | 121.0 | 7.94 | 7.93 | - | 18.4 | 24-36 |
| 612 | Q | 56.3 | 56.7 | 176.4 | 176.4 | 118.0 | 117.9 | 7.90 | 7.89 | - | 28.9 | 24-36 |
| 613 | F | 58.0 | 58.5 | 176.5 | 176.5 | 119.6 | 119.5 | 8.10 | 8.09 | - | 39.3 | 24-36 |
| 614 | G | 44.9 | 45.4 | 173.9 | 173.9 | 109.5 | 109.5 | 8.19 | 8.20 | - | - | 24-36 |
| 615 | L | 54.8 | 55.2 | 177.6 | 177.6 | 121.4 | 121.3 | 8.03 | 8.02 | - | 42.2 | 24-36 |

| Residue | | $^{13}C_{\alpha}$ | | $^{13}CO$ | | $^{15}N$ | | $^{1}HN$ | | $H_{\alpha}$ | $C_{\beta}$ | Source |
|---|---|---|---|---|---|---|---|---|---|---|---|---|
| # | AA | hTE | Frag. | hTE | Frag. | hTE | Frag. | hTE | Frag. | Frag. | Frag. | Frag. |
| 616 | V | 62.4 | 62.7 | 176.8 | 176.8 | 120.9 | 120.9 | 8.18 | 8.17 | - | 32.7 | 24-36 |
| 617 | G | 44.8 | 45.2 | 174.0 | 174.0 | 112.6 | 112.6 | 8.49 | 8.50 | - | - | 24-36 |
| 618 | A | 52.3 | 52.9 | 177.9 | 177.8 | 123.9 | 123.9 | 8.24 | 8.24 | - | 19.2 | 24-36 |
| 619 | A | 52.4 | 52.8 | 178.4 | 178.3 | 122.6 | 122.6 | 8.38 | 8.38 | - | 19.0 | 24-36 |
| 620 | G | 44.6 | 45.3 | 174.4 | 174.4 | 107.4 | 107.5 | 8.28 | 8.28 | - | - | 24-36 |
| 621 | L | - | - | - | 178.2 | 121.5 | 121.5 | - | 8.18 | - | - | 24-36 |
| 622 | G | - | - | - | - | - | 109.6 | - | 8.56 | - | - | 24-36 |
| 623 | G | - | - | - | - | - | 108.6 | - | - | - | - | 24-36 |
| 624 | L | - | - | - | - | - | - | - | - | - | - | - |
| 625 | G | - | - | - | - | - | - | - | - | - | - | - |
| 626 | V | - | - | - | 177.1 | - | 119.1 | - | 8.11 | - | - | 24-36 |
| 627 | G | - | - | - | 174.6 | - | 112.7 | - | 8.67 | - | - | 24-36 |
| 628 | G | 44.8 | - | 174.2 | 174.2 | 108.8 | 108.5 | - | 8.28 | - | - | 24-36 |
| 629 | L | 54.9 | 55.2 | 178.0 | 177.9 | 121.3 | 121.2 | 8.22 | 8.21 | - | 42.1 | 24-36 |
| 630 | G | 44.7 | - | 173.6 | 173.6 | 109.5 | 109.5 | 8.50 | 8.49 | - | - | 24-36 |
| 631 | V | 59.5 | - | 174.6 | - | 121.1 | 121.0 | 8.06 | - | - | - | 24-36 |
| 632 | P | - | 63.4 | - | 177.5 | - | - | - | - | - | 32.2 | 24-36 |
| 633 | G | - | 45.1 | - | 174.3 | - | 109.4 | - | 8.49 | - | - | 24-36 |
| 634 | V | 62.4 | 62.6 | 177.0 | 177.0 | - | 119.4 | - | 8.12 | - | 32.6 | 24-36 |
| 635 | G | 44.9 | 45.3 | 174.6 | 174.6 | 112.8 | 112.8 | 8.69 | 8.69 | - | - | 24-36 |
| 636 | G | 44.8 | 45.2 | 174.4 | 174.4 | 108.7 | 108.6 | 8.32 | 8.31 | - | - | 24-36 |
| 637 | L | 54.9 | 55.5 | 178.2 | 178.2 | 121.9 | 121.7 | 8.31 | 8.31 | - | 42.1 | 24-36 |
| 638 | G | 44.8 | 45.3 | 174.6 | 174.6 | 109.7 | 109.8 | 8.59 | 8.60 | - | - | 24-36 |
| 639 | G | 44.5 | 45.0 | 173.6 | 173.6 | 108.6 | 108.5 | 8.23 | 8.23 | 3.90 | - | 24-36 |
| 640 | I | 58.3 | 58.6 | 174.4 | 174.4 | 122.1 | 122.0 | 8.12 | 8.11 | 4.42 | 38.3 | 24-36 |
| 641 | P | - | - | - | - | - | - | - | - | - | - | - |
| 642 | P | 62.9 | 63.4 | 177.2 | 177.1 | - | - | - | - | 4.34 | 32.1 | 24-36 |
| 643 | A | 52.5 | 52.9 | 178.1 | 178.1 | 123.8 | 123.8 | 8.51 | 8.51 | 4.19 | 18.6 | 24-36 |
| 644 | A | 52.4 | - | 178.2 | - | 122.9 | 122.9 | 8.24 | 8.25 | - | - | 24-36 |
| 645 | A | 52.4 | - | 178.1 | - | 123.3 | 123.7 | 8.25 | - | - | - | 24-36 |
| 646 | A | 52.7 | 53.0 | 178.3 | 178.3 | 123.0 | 123.0 | 8.25 | 8.25 | - | 18.6 | 24-36 |
| 647 | K | 56.4 | 56.9 | 176.8 | 176.7 | 120.4 | 120.4 | 8.18 | 8.19 | - | 32.8 | 24-36 |
| 648 | A | 52.4 | 52.6 | 177.9 | 177.8 | 124.4 | 124.3 | 8.21 | 8.20 | - | 18.7 | 24-36 |
| 649 | A | 52.4 | 52.8 | 177.9 | 177.9 | 123.2 | 123.2 | 8.21 | 8.21 | - | 18.7 | 24-36 |
| 650 | K | 56.3 | 56.7 | 176.5 | 176.5 | 120.2 | 120.2 | 8.15 | 8.15 | - | 32.7 | 24-36 |
| 651 | Y | 57.7 | 58.0 | 176.5 | 176.4 | 120.3 | 120.3 | 8.20 | 8.20 | - | 38.6 | 24-36 |
| 652 | G | 44.9 | 45.3 | 173.9 | 173.9 | 110.5 | 110.5 | 8.27 | 8.28 | - | - | 24-36 |
| 653 | A | 52.9 | - | 177.8 | - | 124.0 | 123.8 | 8.21 | 8.20 | - | - | 24-36 |
| 654 | A | 52.4 | 52.8 | 178.3 | 178.3 | 122.5 | 122.6 | 8.39 | 8.39 | - | 19.0 | 24-36 |
| 655 | G | 44.9 | 45.3 | 174.4 | 174.4 | 107.6 | 107.5 | 8.27 | 8.27 | - | - | 24-36 |
| 656 | L | 54.9 | - | 178.2 | - | 121.6 | 121.5 | 8.20 | 8.17 | - | - | 24-36 |

| Residue | | $^{13}C_{\alpha}$ | | $^{13}CO$ | | $^{15}N$ | | $^{1}HN$ | | $H_{\alpha}$ | $C_{\beta}$ | Source |
|---|---|---|---|---|---|---|---|---|---|---|---|---|
| # | AA | hTE | Frag. | hTE | Frag. | hTE | Frag. | hTE | Frag. | Frag. | Frag. | Frag. |
| 657 | G | 44.8 | - | 174.7 | 174.6 | 109.7 | - | 8.56 | - | - | - | 24-36 |
| 658 | G | 44.9 | - | 174.2 | 174.2 | 108.6 | 108.5 | 8.24 | 8.24 | 3.92 | - | 24-36 |
| 659 | V | 62.3 | 62.4 | 176.6 | 176.5 | 119.6 | 119.5 | 8.08 | 8.07 | 4.07 | 32.7 | 24-36 |
| 660 | L | 55.0 | 55.3 | 177.9 | 177.9 | 125.7 | 125.8 | 8.49 | 8.49 | 4.31 | 42.0 | 24-36 |
| 661 | G | 44.9 | - | 174.7 | 174.6 | 110.1 | 110.0 | 8.43 | 8.44 | - | - | 24-36 |
| 662 | G | 44.8 | 45.2 | 174.0 | 174.0 | 108.9 | 108.7 | - | 8.29 | - | - | 24-36 |
| 663 | A | 52.4 | 52.7 | 178.4 | 178.3 | 123.7 | 123.6 | 8.36 | 8.35 | 4.28 | 19.1 | 24-36 |
| 664 | G | 44.8 | 45.3 | 173.9 | 173.7 | 108.1 | 108.0 | 8.47 | 8.47 | 3.87 | - | 24-36 |
| 665 | Q | 55.3 | 55.8 | 175.3 | 175.2 | 119.1 | 119.2 | 8.07 | 8.06 | 4.10 | 29.4 | 24-36 |
| 666 | F | 55.1 | 55.6 | 173.7 | 173.7 | 121.6 | 121.5 | 8.31 | 8.31 | 4.86 | 38.8 | 24-36 |
| 667 | P | 62.9 | 63.2 | 177.0 | 177.0 | - | - | - | - | 4.36 | 31.8 | 24-36 |
| 668 | L | 55.1 | 55.5 | 178.2 | 178.2 | 122.7 | 122.7 | 8.50 | 8.51 | 4.29 | 42.0 | 24-36 |
| 669 | G | 44.9 | 45.1 | 174.6 | 174.6 | 109.9 | 110.0 | 8.54 | 8.54 | 3.89 | - | 24-36 |
| 670 | G | 44.8 | 45.1 | 174.0 | 174.0 | 108.8 | 108.6 | - | 8.29 | 3.91 | - | 24-36 |
| 671 | V | 61.9 | 62.2 | 175.9 | 175.9 | 119.3 | 119.1 | 8.08 | 8.07 | 4.02 | 32.9 | 24-36 |
| 672 | A | 51.9 | 52.3 | 177.2 | 177.2 | 127.4 | 127.4 | 8.38 | 8.38 | 4.17 | 19.1 | 24-36 |
| 673 | A | 52.0 | 52.3 | 177.4 | 177.4 | 123.3 | 123.3 | 8.22 | 8.23 | - | 19.2 | 24-36 |
| 674 | R | 53.4 | 53.9 | 174.1 | 174.1 | 121.5 | 121.5 | 8.31 | 8.31 | 4.57 | 30.0 | 24-36 |
| 675 | P | 63.2 | 63.6 | 177.5 | 177.5 | - | - | - | - | 4.32 | 31.8 | 24-36 |
| 676 | G | 44.8 | 45.2 | 173.9 | 173.9 | 110.0 | 109.9 | 8.58 | 8.58 | 3.78 | - | 24-36 |
| 677 | F | 57.9 | 58.4 | 176.3 | 176.2 | 120.3 | 120.1 | 8.18 | 8.17 | 4.51 | 39.7 | 24-36 |
| 678 | G | 44.7 | 45.1 | 173.8 | 173.7 | 111.2 | 111.2 | 8.46 | 8.45 | 3.83 | - | 24-36 |
| 679 | L | 54.4 | 54.8 | 177.4 | 177.3 | 121.3 | 121.2 | 8.09 | 8.08 | 4.36 | 42.5 | 24-36 |
| 680 | S | 56.2 | 56.5 | 172.5 | 172.5 | 118.3 | 118.3 | 8.41 | 8.41 | 4.69 | 63.3 | 24-36 |
| 681 | P | 62.9 | 63.1 | 176.2 | 176.2 | - | - | - | - | 4.34 | 32.0 | 24-36 |
| 682 | I | 60.6 | 60.9 | 175.7 | 175.7 | 120.5 | 120.4 | 8.11 | 8.10 | 4.01 | 38.8 | 24-36 |
| 683 | F | 54.9 | 55.2 | 174.0 | 174.0 | 125.4 | 125.5 | 8.41 | 8.41 | 4.90 | 39.0 | 24-36 |
| 684 | P | 63.3 | 63.5 | 177.5 | 177.4 | - | - | - | - | 4.35 | 31.7 | 24-36 |
| 685 | G | 45.0 | 45.3 | 174.7 | 174.7 | 109.5 | 109.5 | 8.32 | 8.31 | 3.92 | - | 24-36 |
| 686 | G | 44.8 | 45.1 | 173.8 | 173.7 | 108.9 | 108.8 | 8.30 | 8.28 | 3.91 | - | 24-36 |
| 687 | A | 52.0 | 52.4 | 177.1 | 177.1 | 123.6 | 123.5 | 8.25 | 8.25 | 4.27 | 19.1 | 24-36 |
| 688 | C | 56.2 | 56.5 | 173.5 | 173.5 | 119.2 | 119.1 | 8.43 | 8.43 | 4.50 | 41.6 | 24-36 |
| 689 | L | 54.3 | 54.6 | 177.3 | 177.3 | 126.4 | 126.3 | 8.49 | 8.49 | 4.49 | 43.2 | 24-36 |
| 690 | G | 44.9 | 45.3 | 175.0 | 175.0 | 111.5 | 111.4 | 8.59 | 8.59 | 3.79 | - | 24-36 |
| 691 | K | 56.9 | 57.3 | 177.2 | 177.2 | 121.2 | 121.1 | 8.54 | 8.51 | 4.19 | 32.9 | 24-36 |
| 692 | A | 52.5 | 53.0 | 178.2 | 178.2 | 122.5 | 122.4 | 8.44 | 8.43 | 4.29 | 18.1 | 24-36 |
| 693 | C | 55.4 | 55.7 | 175.5 | 175.5 | 116.8 | 116.8 | 7.90 | 7.88 | 4.50 | 41.5 | 24-36 |
| 694 | G | 45.1 | 45.4 | 173.9 | 173.8 | 109.9 | 109.8 | 8.42 | 8.42 | 3.91 | - | 24-36 |
| 695 | R | 55.7 | 56.1 | 176.3 | 176.2 | 120.8 | 120.7 | 8.11 | 8.09 | 4.32 | 30.7 | 24-36 |
| 696 | K | 55.9 | 56.4 | 176.5 | 176.4 | 123.3 | 123.3 | 8.49 | 8.47 | 4.26 | 32.9 | 24-36 |
| 697 | R | 55.7 | 56.1 | 175.3 | 175.3 | 123.7 | 123.7 | 8.50 | 8.49 | 4.28 | 30.7 | 24-36 |

| Residue | | $^{13}C_{\alpha}$ | | $^{13}CO$ | | $^{15}N$ | | $^{1}HN$ | | $H_{\alpha}$ | $C_{\beta}$ | Source |
|---|---|---|---|---|---|---|---|---|---|---|---|---|
| # | AA | hTE | Frag. | hTE | Frag. | hTE | Frag. | hTE | Frag. | Frag. | Frag. | Frag. |
| 698 | K | 57.5 | 57.9 | 181.3 | 181.3 | 128.0 | 128.0 | 8.10 | 8.09 | 4.07 | 33.4 | 24-36 |

**SI Table 3. Unambiguous backbone chemical shift assignment statistics**

| Construct | HN (%) | CO (%) | $C_\alpha$ (%) | $H_\alpha$ (%) |
|---|---|---|---|---|
| hTE | 77.9 | 79.4 | 71.3 | N/A |
| 2-8 | 100 | 99.1 | 100 | 96.7 |
| 8-14 | 99 | 98.4 | 100 | 98.1 |
| 2-14 | 96.2 | 93.7 | 90 | 61.2 |
| 13-19 | 100 | 99.4 | 99.4 | 99.1 |
| 18-26 | 91.5 | 90.8 | 84.2 | 78.8 |
| 20-24 (EP1)[a] | 77.2 | 66.9 | 54 | 35.1 |
| 24-36 | 96.7 | 95.2 | 93.1 | 82.5 |
| ELP-29-36[a] | 83.8 | 81.6 | 80.9 | 69.2 |

*a* – These mutant or hybrid constructs were used for comparison purposes to aid in the assignment of constructs with WT sequences. None of the chemical shifts from these constructs were used for δ2D secondary structure prediction for the purposes of generating IDPConformerGenerator restraints.

**SI Table 4. IDPConformerGenerator parameters**

| Option/Flag | Value | Description |
|---|---|---|
| -db | dpconfgen_database_2023.json | Database containing non-redundant structural motifs from PDB |
| -seq | hTE_Sx2.seq | FASTA file containing hTE sequence with C688 and C693 mutated to serine |
| -etbb | 250 | Backbone energy threshold |
| -etss | 500 | File with x-mer probabilities for motif selection |
| -xp | xmer_probs.csv<br>1 0<br>2 0<br>3 2<br>4 2<br>5 2<br>6 2<br>7 1<br>8 1<br>9 0 | File with x-mer probabilities for motif selection |
| -csss | hTE_d2D.json | d2D Secondary structure propensity file from chemical shifts |
| -long | - | Flag to indicate long fragment generation |
| --long-ranges | 1-116,117-222,223-360,361-468,469-584,585-698 (50%)<br><br>1-58,59-116,117-170,171-222,223-292,293-360,361-415,416-468,469-526,527-584,585-638,639-698 (50%) | Defines ranges for each fragment during model building |
| --dloop-off | - | Turns off loop closure during fragment stitching |
| -et | pairs | Energy threshold for pairwise interactions |

**SI Table 5. GNOM fits to hTE variant experimental SAXS data**

| | C688S/693S 100 µM | | WT 100 µM | | C688S/693S 200 µM | | WT 200 µM | |
|---|---|---|---|---|---|---|---|---|
| **Dmax (Å)** | $R_g$ (Å) | $\chi^2$ | $R_g$ (Å) | $\chi^2$ | $R_g$ (Å) | $\chi^2$ | $R_g$ (Å) | $\chi^2$ |
| 150 | 49.9 ± 1.2 | 1.61 | 48.1 ± 2.5 | 0.81 | 52.4 ± 1.0 | 0.78 | 52.5 ± 1.0 | 0.64 |
| 160 | 52.6 ± 1.3 | 1.60 | 50.3 ± 2.1 | 0.82 | 54.1 ± 1.3 | 0.77 | 55.1 ± 1.6 | 0.62 |
| 170 | 55.3 ± 1.1 | 1.60 | 52.9 ± 1.9 | 0.82 | 56.7 ± 1.9 | 0.75 | 57.9 ± 2.7 | 0.60 |
| 180 | 57.7 ± 1.2 | 1.60 | 55.7 ± 2.0 | 0.81 | 57.1 ± 3.9 | 0.75 | 58.3 ± 2.9 | 0.60 |
| 190 | 59.9 ± 1.5 | 1.60 | 57.7 ± 2.1 | 0.82 | 57.5 ± 3.7 | 0.75 | 58.5 ± 2.6 | 0.61 |
| 200 | 62.0 ± 1.8 | 1.60 | 58.1 ± 3.3 | 0.80 | 58.5 ± 4.3 | 0.76 | 58.9 ± 2.6 | 0.62 |
| 210 | 64.5 ± 7.1 | 1.49 | 58.7 ± 4.1 | 0.81 | 59.4 ± 4.8 | 0.76 | 59.4 ± 2.9 | 0.62 |
| 220 | 66.0 ± 2.8 | 1.59 | 59.4 ± 4.7 | 0.82 | 60.3 ± 5.2 | 0.76 | 62.0 ± 1.3 | 0.70 |
| 230 | 68.1 ± 2.5 | 1.60 | 62.9 ± 3.1 | 0.87 | 61.7 ± 4.4 | 0.77 | 63.0 ± 1.5 | 0.70 |
| 240 | 70.5 ± 2.7 | 1.60 | 65.0 ± 2.4 | 0.91 | 63.3 ± 3.3 | 0.78 | 64.3 ± 1.7 | 0.70 |
